# APOE+ Tumor-Associated Macrophages and CD4-DOCK4 T Cells Reveal Distinct Microenvironmental Features in HER2-Low and HER2-0 Hormone Receptor-Positive Breast Cancer

**DOI:** 10.1101/2025.09.04.674072

**Authors:** Carlos Wagner S. Wanderley, Daniel E. Michaud, Kenichi Shimada, Adam Nelson, Jingxin Fu, Stuart J. Schnitt, Sara M Tolaney, Elizabeth A. Mittendorf, Romualdo Barroso-Sousa, Adrienne Waks, Paolo Tarantino, Jennifer L. Guerriero

## Abstract

Novel anti-HER2 antibody-drug conjugates (ADCs), such as trastuzumab deruxtecan (T-DXd), have shown efficacy in tumors with varying HER2 expression, including HER2-low and even tumors with minimal HER2 presence. This has sparked interest in the biology underlying the HER2 expression spectrum. Using molecular and multiplexed imaging, we revealed distinct immune and stromal features in treatment-naive, hormone receptor-positive (HR+) HER2-low versus HER2-0 tumors. HER2-0 tumors exhibit inflammatory and tissue remodeling gene signatures, with enrichment of APOE⁺ tumor-associated macrophages (TAMs) and DOCK4⁺ CD4 T cells. In contrast, HER2-low tumors are more immunosuppressed, with elevated cell cycle, metabolic, and estrogen signaling pathways, suggesting increased proliferative activity. These findings underscore key biological differences between HR+ HER2-low and HER2-0 breast cancers, and may inform more tailored therapeutic strategies.

**Statement of significance:** This study revealed the distinct biological profiles of HR+ HER2-low and HER2-0 breast tumors. HER2-0 tumors exhibit inflammatory and tissue remodeling signatures, whereas HER2-low tumors have elevated cell cycle, metabolic, and estrogen signaling. These insights may help refine therapeutic approaches to improve outcomes for breast cancer patients.

## Introduction

The biological heterogeneity of breast cancer challenges treatment but also provides opportunities for innovation in precision medicine. For over two decades, the eligibility for traditional anti-HER2 targeted therapy was determined based on HER2 status in a binary fashion: Positive if harboring HER2 overexpression (immunohistochemistry [IHC] 3+) or epidermal growth factor receptor two gene 2 (*ERBB2)* amplification (positive fluorescence *in situ* hybridization [FISH]), or negative if tumors have low or no HER2 expression (IHC 0, 1+ or 2+ with *ERBB2* FISH unamplified)(1). However, the results of phase three clinical trials have demonstrated that the novel anti-HER2 antibody drug-conjugate (ADC) trastuzumab deruxtecan (T-DXd) improves progression-free and overall survival of patients with HER2-low (IHC 1+ or 2+ with *ERBB2* FISH unamplified) (1–3). As a result, there is a need to better understand the molecular and immunological features underlying the HER2 expression spectrum, to inform more refined patient stratification and the development of novel therapeutic approaches.

The genomic features of breast cancer play a critical role in tumor growth, metastasis, immune evasion, and therapy response(4). In hormone receptor-positive (HR+) breast tumors, the tumor microenvironment (TME) typically exhibits lower tumor-infiltrating lymphocytes (TIL) density and a higher proportion of tumor-associated macrophages (TAMs) compared to triple-negative breast cancer(5). HER2-low tumors account for approximately 45-55% of breast carcinomas, of which 60-80% are HR+(6). Previous studies have shown that HR+ HER2-low tumors have an immune-depleted profile and marginal differences in TIL density compared with HER2-0 tumors (7–9). However, to date, no comprehensive analysis has compared the TME of HER2-low and HER2-0 HR+ breast tumors.

In this study, using molecular and multiplexed imaging we conducted an in-depth characterization of the immune and stromal features of treatment-naïve, early-stage HR+ HER2- low and HER2-0 breast tumors, providing insights into their distinct biological profiles and potential implications for future therapeutic strategies.

## Results

### HER2-low and HER2-0 HR+ breast cancers exhibit distinct immune and cell cycle gene expression profiles

A total of 25 patients with primary, treatment-naïve, HR+ breast cancer, categorized as either HER2-low (IHC 1+, 2+/FISH Neg., n=17) or HER2-0 (IHC 0, n=8) were included. Three HR+ HER2-positive (IHC 3+, 2+/FISH Pos.) patients were used as references for *ERBB2* gene expression analysis. Age, menopausal status, and histological grade were well-balanced between the two groups. Detailed demographics and clinicopathological characteristics according to HER2 status are reported in **Supplementary Table 1**. To investigate the TME landscape, we performed bulk RNA sequencing (RNA-seq), single-nucleus sequencing (sNuc- seq), and multi-plex cyclic immunofluorescence (CyCIF) (**Fig. 1A**). Bulk tumor RNA-seq analysis confirmed that *ERBB2* expression varied according to the HER2 status. Specifically, HER2-low tumors exhibited higher *ERBB2* gene expression compared to HER2-0 (p=0.01) but lower than HER2-positive tumors (p=0.008) (**Fig. 1B**). Principal component analysis (PCA) revealed a general overlap between the groups, with some potential differences observed (**Supplementary Fig. 1)**. This was further supported by the identification of 203 differentially expressed genes (DEGs) between HER2-0 and HER2-low tumors (**Fig. 1C, and Supplementary Table 2**). Gene set enrichment analysis (GSEA) of biological processes (BP) revealed that differentially expressed genes (DEGs) in HER2-0 tumors were associated with immune responses, including “adaptive immune response” (adjusted p-value or padj<0.001), “positive regulation of immune response” (padj<0.001), and “activation of immune response” (padj<0.001) among others (**Fig. 1D**). In contrast, DEGs in HER2-low tumors primarily belonged to cell cycle processes, including “E2F targets” (padj<0.001), “cell cycle checkpoints” (padj<0.001), and “mitotic nuclear division” (padj<0.001) (**Fig. 1D**). Tumor immune estimation recourse (TIMER2.0)(10) was used to assess the relative fraction of immune cells and revealed that CD4 T cells (p=0.029) and macrophages (p=0.003) were increased in HER2-0 compared to HER2-low breast tumors (**Supplementary Fig. 2**).

**Figure 1.**
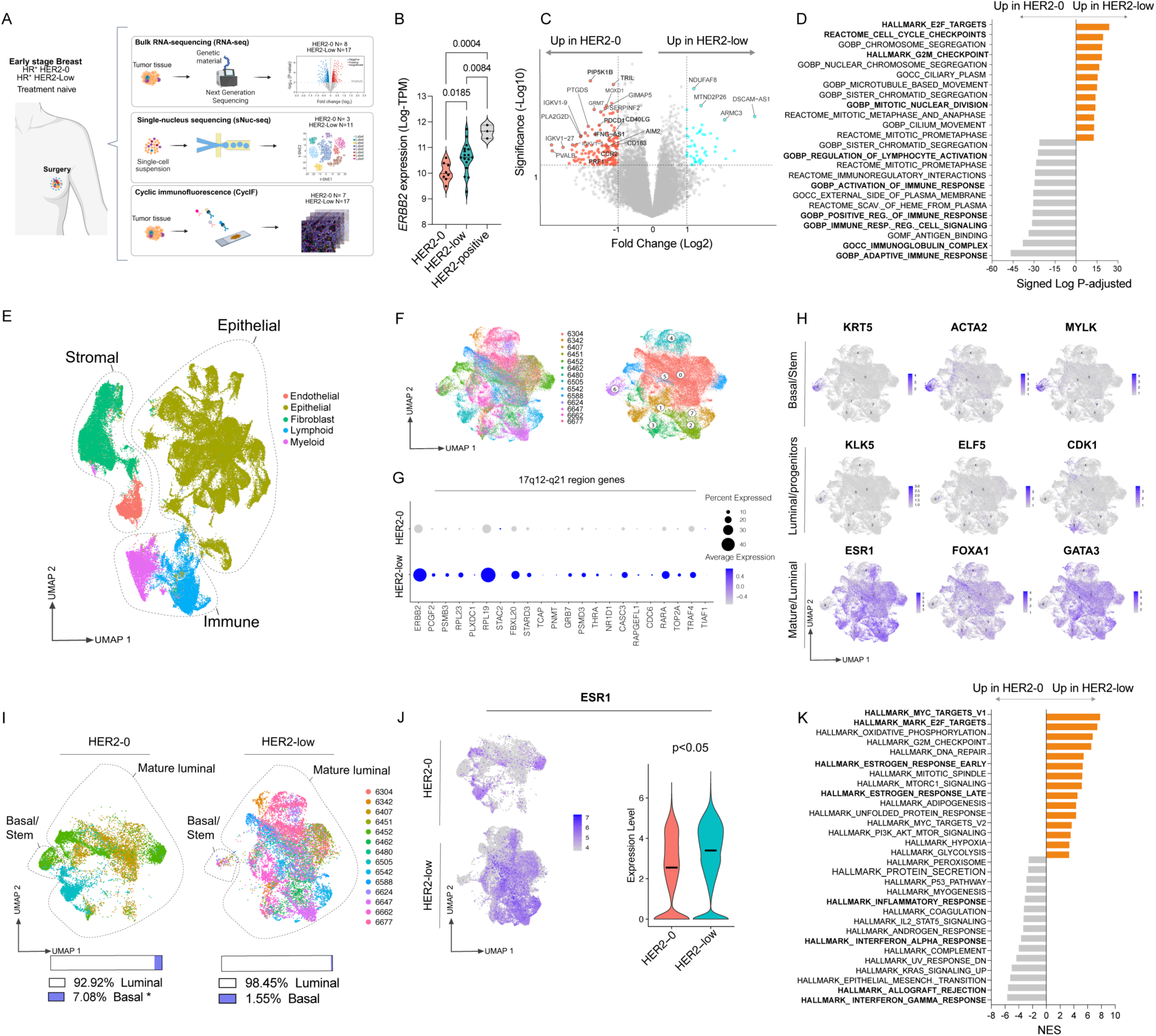
HER2-low and HER2-0 HR+ breast cancers exhibit distinct immune and cell cycle gene expression profiles. (A) Schematic of experimental design and clinicopathological information of samples (created with BioRender.com). (B-D) Bulk RNA-seq analysis comparing HER2-low (n=17) and HER2-0 (n=8) tumors. (B) The expression level of *ERBB2* by Log- transcripts per million (Log-TPM), grouped by HER2 status. (C) Volcano plot exhibiting 203 DEGs among HER2-low vs. HER2-0 tumors is shown. (D) Gene set enrichment analysis (GSEA) reveals up- and down-regulated pathways in HER2-low compared to HER2-0. (E-L) Single nucleus RNA sequencing (sNuc-Seq) analysis comparing HER2-low (n=11) and HER2-0 (n=3). (E) UMAP visualization of 90,248 single nuclei, revealing epithelial, stromal (fibroblast and endothelial cells), and immune (lymphoid and myeloid) cells. (F) UMAP of epithelial tumor cells, color coded by samples (left) and 8 clusters (right). (G) Dot plots showing 17q12-q21 amplicon genes in isolated epithelial cells. The dot size indicates the proportion of expressing cells, colored by the average log scale expression values. (H) UMAP plots of normalized expression of markers used to identify tumor cell lineage states. (I) UMAP plots are split by HER2 status and colored by patient. The proportion of tumor cells is classified according to tumor cell lineage states highlighted in the panel H. (J) ESR1 gene expression in epithelial cells. (K) HALLMARK pathways significantly up- or down-regulated in HER2-low compared to HER2-0 tumor cells based on sNuc-seq data. Significance was assessed using ANOVA, followed by the Tukey or Wilcoxon rank-sum tests, was used appropriately for statistical analysis (p<0.05).

To investigate the contributions of individual cell types to the differential gene signatures, we performed single-nucleus RNA sequencing (sNuc-seq). Using the Seurat R package(11), we performed fine clustering of 90,248 single nuclei from 14 breast tumor resections (HER2-low n=11 and HER2-0 n=3). The distribution of these nuclei was visualized using uniform manifold approximation and projection (UMAP) and categorized based on their cell types using canonical markers of epithelial, immune (myeloid and lymphoid cells), and stromal cells (fibroblasts and endothelial cells) (see Methods) (**Fig. 1E and Supplementary Fig 3A**). Initially, we focused on the epithelial cells from HER2-0 and HER2-low tumors (**Fig. 1F**) and explored the copy number gene variation (CNV). The HER2 gene (*ERBB2*) is located on chromosome 17q12 and is frequently co-amplified with adjacent genes in this region. Using inferred CNV (inferCNV) analysis, we observed that HER2-low epithelial cells showed a gain in the region of genes previously observed co-amplified with *ERBB2,* including *STARD3*, *GRB7*, *RARA*, and *TOP2A* (**Supplementary Fig 3B-C and Supplementary Table 3**). We further confirmed the different expression levels of *ERBB2* and these co-amplified genes through single gene analysis (**Fig. 1G**). Additionally, based on a cell-origin gene signature(12), we identified that seven out of eight clusters from both HR+ HER2-0 and HER2-low patients express mature luminal markers. However, the cluster with basal signature was predominantly found in HER2-0 tumors (HER2-0 = 7.08% vs. HER2-low = 1.55%, p<0.05) (**Fig. 1H-I**). Furthermore, although both are HR+, with comparable levels of ER expression by IHC (averaging 93% in HER2-0 and 90% in HER2-low) the gene expression levels of *ESR1* were higher in HER2-low (**Fig. 1J and supplementary Fig. 4)**. Then, to better understand the different biological processes between HER2-0 and HER2- low tumor cells, we performed a GSEA on the epithelial cell clusters. We observed that immune-related biological processes were enriched in HER2-0, including “interferon-gamma response” (padj<0.001) and “inflammatory response” (padj<0.001). In contrast, HER2-low was enriched with pathways associated with cell cycle, metabolism, and estrogen signaling, such as “MYC targets” (padj<0.001), “E2F targets” (padj<0.001), “oxidative phosphorylation” (padj<0.001), “glycolysis” (padj<0.001), “estrogen early response” (padj<0.001), and “estrogen late response” (padj<0.001) (**Fig. 1K**).

Collectively, these findings suggest that tumor cells in HER2-0 breast tumors exhibit inflammatory characteristics, whereas HER2-low tumors exhibit enhanced estrogen signaling, cell cycle regulation, and energy metabolism.

### HER2-low and HER2-0 HR+ breast tumors display distinct immune profiles

We hypothesized that the enrichment of immune-related pathways identified in our GSEA analysis of bulk RNA-seq data from whole tumor (**Fig. 1D**) and sNuc-seq data from tumor epithelial cells (**Fig. 1K**) from HER2-0 compared to HER2-low tumors could be associated with increased immune infiltration and/or activation in HER2-0 tumors. Therefore, we reclustered the previously identified immune cell populations to examine the immune milieu at a higher resolution (**Fig. 2A-B and Supplementary Fig. 5A**). We identified and characterized the frequency of immune cells in HER2-0 and HER2-low breast tumors (**Fig. 2C-F, Supplementary Fig. 5B**). Out of all immune cells, we observed an increased proportion of B cells in HER2-0 compared to HER2-Low (1.17% vs. 0.23%, p=0.03) (**Supplementary Fig. 5C**). However, analysis of immune subsets out of all cells, a surrogate of total numbers of cells within a tumor, revealed significant differences between HER2-0 and HER2-low tumors (**Fig. 2F**). We identified B cells (3.7% vs. 1.57%, p=0.009), CD4 (4.5% vs. 0.9%, p=0.03), CD8 (8.3% vs. 2.6%, p=0.02) conventional dendritic cells 1 and 2 (cDC1 0.27% vs. 0.09%, p=0.01 and cDC2, 0.3% vs. 0.07%, p=0.01), macrophages (8.9% vs. 5.7%, p=0.03), NK cells (0.45% vs. 0.11%, p=0.04), plasmacytoid dendritic cells (pDCs, 0.37% vs. 0.08% vs., p=0.005), and plasma cells (2.7% vs. 0.3%, p=0.02), p=0.02) were more abundant in HER2-0 than in HER2-low samples (**Fig. 2F**).

**Figure 2.**
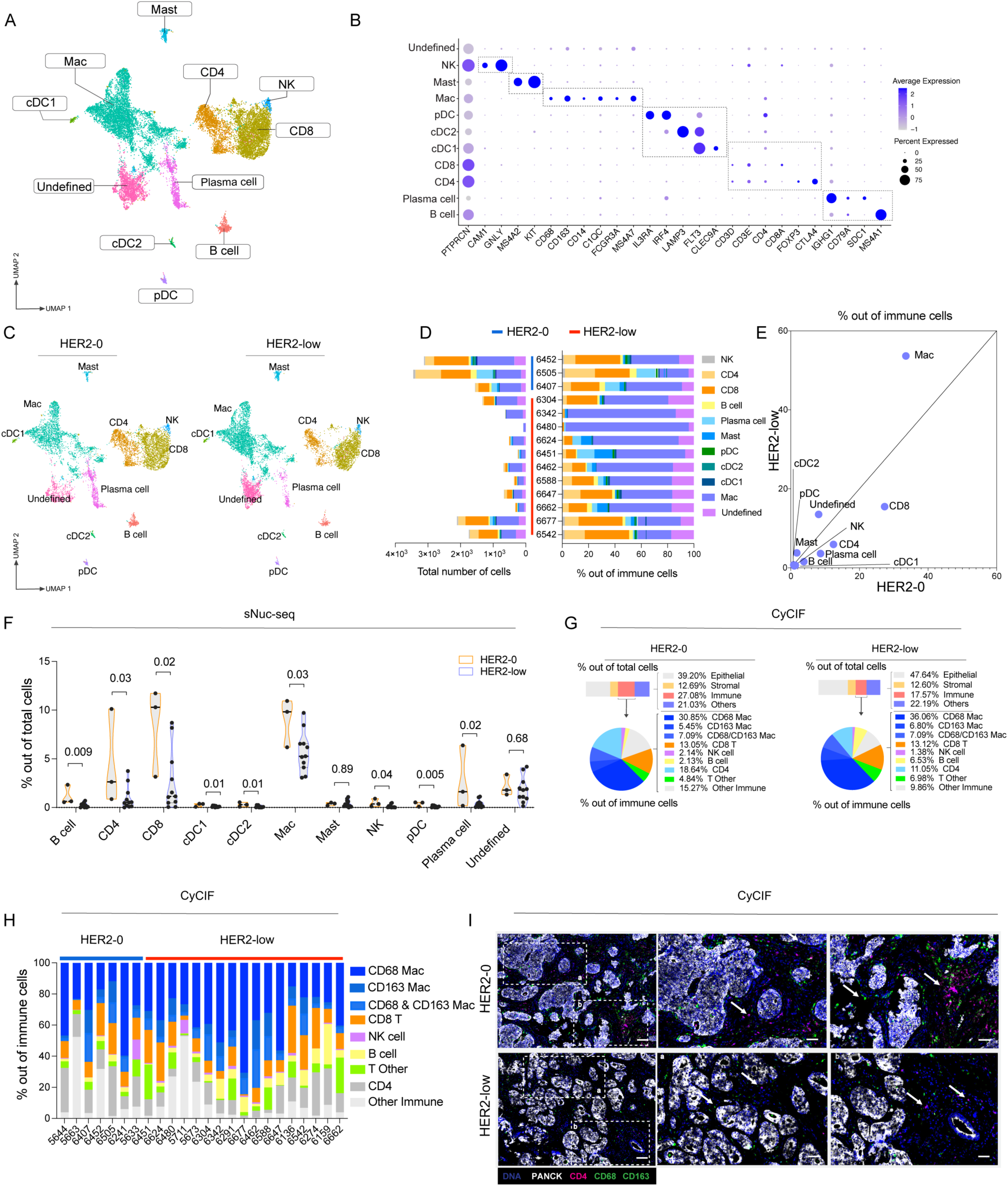
HER2-low and HER2-0 HR+ breast tumors display distinct immune profiles. (A- F) sNuc-Seq data analysis of HER2-low (n=11) and HER2-0 (n=3). (A) UMAP plots show immune cell (PTPRC+) subpopulations colored by cell type. (B) Dot plots showing marker genes for cell types in Fig. 2A. Dot size indicates the proportion of expressing cells, colored by the average of log-TPM values. (C) UMAP plot showing the distribution of immune cells colored by cell type in HER2-0 and HER2-low samples. (D) The absolute number (left) and immune cell type proportion (right) are assigned to each patient from immune cells (PTPRC+). (E) Scatter plot comparing average proportions of immune cell types for HER2-0 (x-axis) and HER2-low (y- axis) samples (out of immune cells PTPRC+). (F) Fractions of the indicated immune cell subtypes from the total of cells. (G-I) Cyclic immunofluorescence (CyCIF) analysis of HER2 low (n=17) and HER2-0 (n=7) tumors. (G) The average fractions of epithelial, stromal, immune, and others out of total cells (top) and immune cell subtype proportion from CD45+ immune cell compartment (bottom). (H) The fraction of immune cells out of CD45+ cells shown by patient. (I) Representative images of a breast tumors, DNA (blue), PanCK (white), CD4 (red), CD68 (green), and CD163 (green) staining, to reveal the immune landscape of HER2-low and HER2-0 tumors. Scale bars, where 100 µm (left) and 50 µm (middle and right) are in the insets. A two-tailed t-test was used for statistical analysis (p<0.05).

To confirm the differences in immune cells observed in our sNuc-Seq dataset, we utilized single- cell, multiplex CyCIF. Similar to the sNuc-Seq analysis, the frequency of immune cell subsets out of all immune cells displayed moderate differences between HER2-0 and HER2-low tumors (**Supplementary Fig. 6A**). However, out of all cells, HER2-0 tumors exhibited an average of 10% more immune cells than HER2-low tumors (27.08% in HER2-0 vs. 17.57% in HER2-low, p=0.09), and we observed a significant increase in CD4 T cells infiltration in HER2-0 compared to HER2-low tumors (**Fig. 2G-I and Supplementary Fig. 6B)**. These findings suggest that HER2-0 and HER2-low breast tumors have distinct immune profiles.

### HER2-0 and HER2-low HR+ breast tumors exhibit distinct macrophage subsets

Tumor associated macrophages (TAMs) represent a highly heterogeneous cell population, exhibiting diverse phenotypic and functional profiles that are dependent on the microenvironmental context (13). In the present study, we subclustered TAMs from the sNuc- seq dataset, revealing 10 unique subtypes across HER2-0 and HER2-low tumors (**Fig. 3A and Supplementary Figure 7A**). To further explore TAM heterogeneity and plasticity, we visualized their transcriptional profiles using previously established TAM gene signatures (13) and visualized their transcriptional profiles using a radar plot (**Fig. 3B**). Corroborating the functional diversity and plasticity of TAMs, we observed, for example, that APOE-TAMs were enriched for the lipid-associated (LA)_TAM signature, as previously described(13), but also overlapped with immunosuppressive regulatory and proliferative TAMs, suggesting a dual role in immunosuppression and self-renewal (**Fig. 3B**). DOCK4-TAMs aligned with intermediate monocytes, regulatory TAMs, and LA-TAMs signatures, suggesting a metabolically active and suppressive profile. FLT3-TAMs were primarily enriched in classical tumor-infiltrating monocyte (TIM) and regulatory TAM signatures. This profile suggests a monocytic origin with immunosuppressive features, potentially representing newly recruited monocytes differentiating within the TME. COL6A1-TAMs exhibited strong enrichment for angiogenic TAM, lipid- associated (LA)-TAM, and regulatory TAM programs. This transcriptional pattern indicates a pro- tumoral phenotype, potentially involved in vascular remodeling, lipid metabolism, and immune suppression. F13A1-TAMs were selectively enriched for the resident tissue macrophage (RTM) signature, with limited expression of other TAM-related programs. This suggests a locally adapted, tissue-resident phenotype, possibly maintaining homeostatic or remodeling functions within the tumor niche (**Fig. 3B**). We identified that APOE-TAMs (0.5% vs. 0.19%, p=0.0002), FLT3-TAMs (2.1% vs. 0.9%, p=0.02), and F13A1-TAMs (1.4% vs. 0.6%, p=0.008) were more abundant in HER2-0 compared to HER2-low tumors (**Fig 3C-E and Supplementary Fig. 7B-D)**. Furthermore, global analysis of TAMs in HER2-0 and HER2-low tumors revealed an enrichment of immune pathways, including “positive regulation of immune response” (padj=0.001) and “lymphocyte activation” (padj=0.001) in HER2-0 samples compared to HER2- low (**Fig. 3F**).

**Figure 3.**
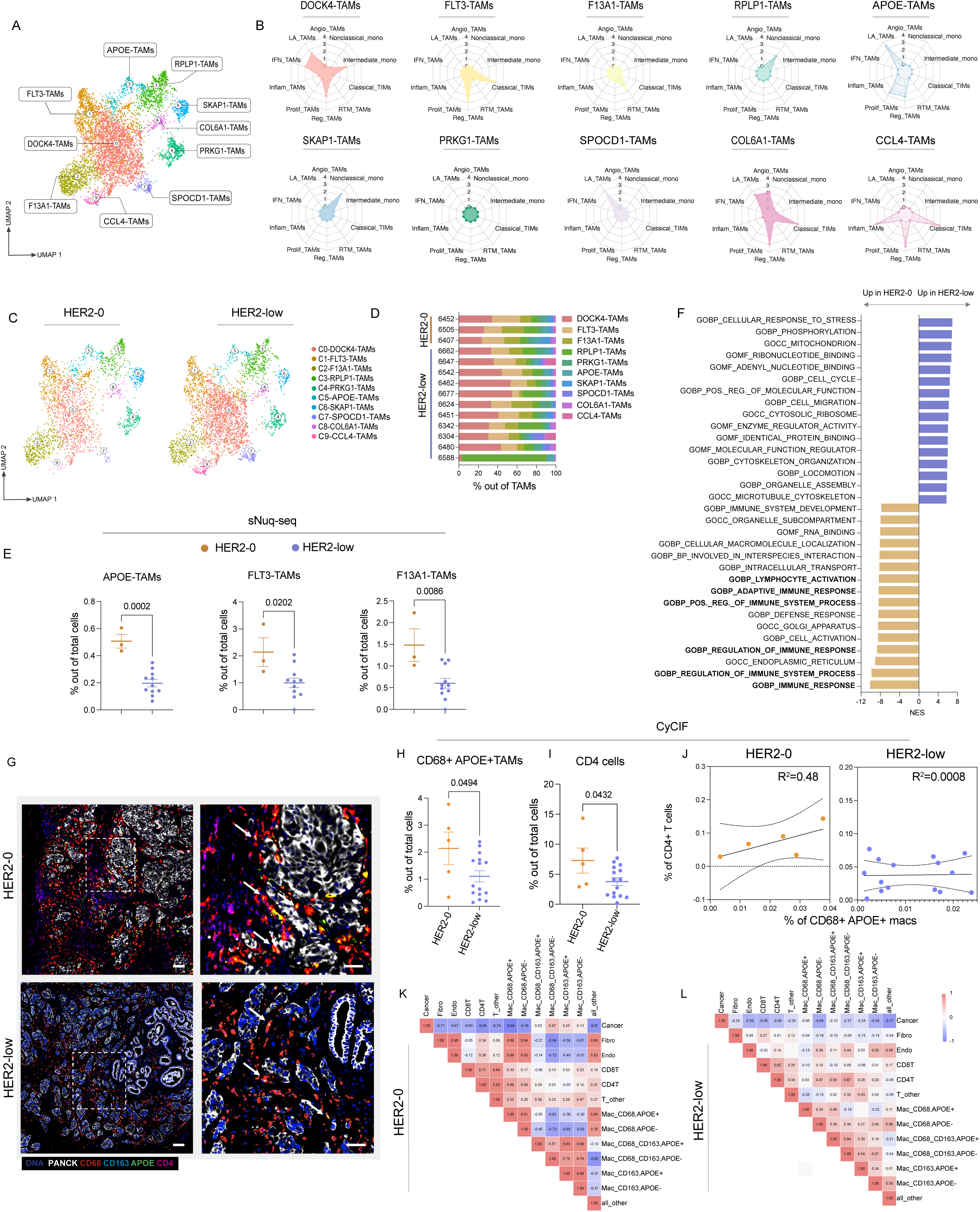
HER2-0 and HER2-low HR+ breast tumors exhibit distinct macrophage subsets. (A-F) Single nucleus RNA sequencing (sNuc-Seq) was performed (HER2 low, n=11; HER2-0, n=3). (A) UMAP plots display tumor-associated macrophage (TAM) subpopulations annotated by their top gene expression profiles. (B) Relative enrichment of phenotypes scores among TAMs subpopulations. Up to four signatures with an adjusted p-value<0.05 were shown. Nonclassical monocyte (Nonclassical mono), Intermediated monocyte (Intermediated mono), Classical tumor infiltrating monocyte (Classical TIM), Resident tissue macrophage (RTM TAMS), Regulatory TAMs (Reg TAMs), Proliferative TAMs (Prolif TAMs), Inflammatory TAMs (Inflamma TAMs), Interferon TAMs (IFN TAMs), Lipid associated TAMs (LA TAMs), and Angiogenic TAMs (angio TAMs). (C) UMAP plot showing the distribution of TAMs colored by subtype in HER2-0 and HER2-low tumors. (D) The proportion of TAM subtypes in each patient. (E) Graphs representing the percentage of TAM subsets out of the total of cells. (F) Gene set enrichment analysis (GSEA) shows the gene ontology biological process (GOBP) pathways that are up-or down-regulated in HER2-low compared to HER2-0. (G-L) CyCIF analysis (HER2 low, n=15; HER2-0, n=5). (G) Representative images, DNA (blue), panCK (white), CD68 (red), CD163 (light blue), APOE (green), and CD4 (purple) staining. Co-localization of CD68 (red) and APOE (green) staining (yellow). Scale bars, where 100 µm (left) and 50 µm (right) are in the insets. (H) Graphs representing the percentage of CD68+APOE+ TAMs and (I) CD4 T cells from the total of cells observed by CyCIF. (J) Proportion of CD68+APOE+ TAMs to CD4 T cells regression analysis. (K-L) Heatmap of Pearson’s correlation coefficient. The two-tailed t-test, Wilcoxon rank-sum test, and Pearson’s correlation were used for statistical analysis (p<0.05).

The APOE-TAM cluster is of significant interest as they were well conserved across several tumors and associated with lipid metabolism and immune regulation(14). APOE-TAMs expressed high levels of CD68, along with lipid-associated genes such as APOC1, LIPA, and TREM2, as well as key antigen presentation genes including HLA-DRA and HLA-DRB1, suggesting a potential role in modulating CD4 T cells responses (**Supplementary Fig. 7E-F and supplementary table 4)**. Therefore, we conducted CyCIF analysis to examine the spatial relationship between APOE-TAMs and CD4 T cells within the tumor. Indeed, CD68+APOE+ macrophages were enriched in HER2-0 tumors compared to HER2-low (**Fig. 3G-H and Supplementary Fig. 8**). Among the other cell types analyzed by CyCIF, we confirmed that CD4 T cells were enriched in HER2-0 compared with HER2-low tumors (**Fig. 3I**). Interestingly, we observed a weak but positive correlation between APOE-TAMs and CD4 T cells (R^2^=0.48 and Pearson’s-r = 0.69) in HER2-0 tumors, which was not detected in HER2-low tumors (R^2^=0.0008 and Pearson’s-r = 0.03) (**Fig 3J-L**). These findings highlight the distinct immune environments of HER2-0 and HER2-low tumors, with APOE+ TAMs and CD4 T cells emerging as critical players in the TME of HER2-0 tumors.

### Analysis of T cell clusters reveals a unique CD4 subset in HER2-0 tumors

Emerging evidence suggests distinct subsets of CD4 and CD8 T cells are found across tumors(15). Our analysis identified 13 distinct T-cell clusters in HER2- 0 and HER2-low samples. CD4 and CD8 T cells were classified based on their gene signatures to reveal functional states, such as naïve, exhausted, and memory (**Fig. 4A-C and Supplementary Fig. 9A-C).** We identified a significant increase in the proportion of CD4-DOCK4 within the T cell compartment (1.9 and 0.6%, p= 0.03) (**Supplementary Fig. 9D-F**) and out of total cells (0.27% vs. 0.04%, p=0.0052) (**Fig. 4D**) in HER2-0 compared to HER2-low tumors. Other subsets of T cells were also enriched in HER2-0 tumors including CD4-DLGAP1, which expressed CTLA-4 and TIGIT, the CD4-PDE7B, which expressed TOX and CXCL13, the CD4/CD8 double positive subset, and CD8 cytotoxic cells (**Fig 4B and D**). The CD4-DOCK4 subset was the mostly highly enriched lymphocyte population in HER2-0 tumors and expressed *GZMK* and *GZMA* as well as gene sets associated with IFNy (padj<0.001) and TNFa signaling (padj<0.001) (**Fig. 4B and E**), suggesting a subtype of CD4 T cells that expresses genes associated with cytotoxicity (15). These data revealed the unique functional states of CD4 T cells, particularly the cytotoxic CD4-DOCK4 subset, which is prominent in HER2-0 tumors.

**Figure 4.**
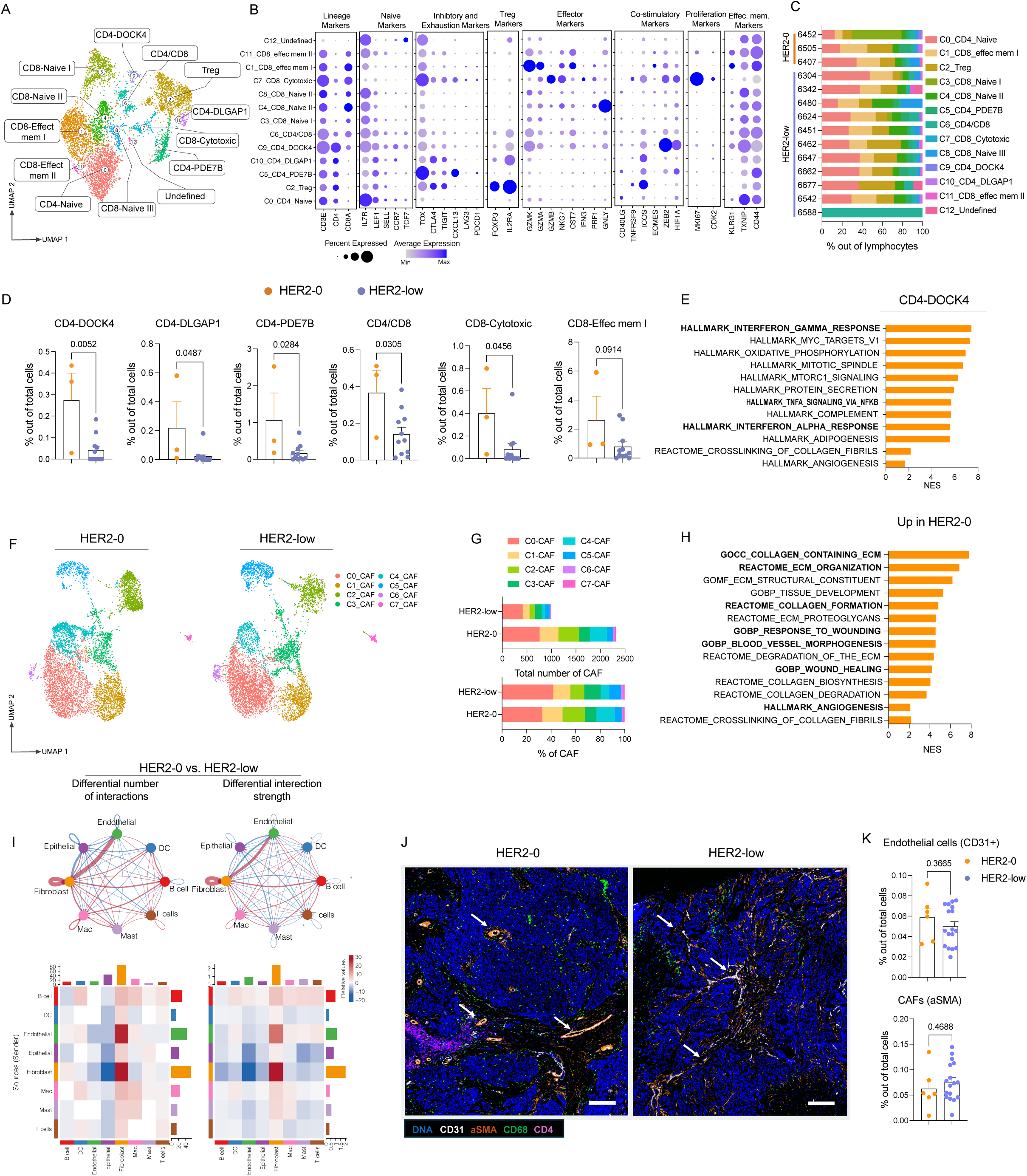
T cells exhibit distinct CD4 functional states, and cancer-associated fibroblasts display unique features in HER2-low and HER2-0, HR+ breast cancer. (A-I) sNuc-Seq analysis (HER2 low, n=11; HER2-0, n=3). (A) UMAP plots show CD4 and CD8 T cell subpopulations labeled with inferred T cell types. (B) Dot plot showing select T cell markers gene expression values (log scale) and percentage of nuclei expressing these genes within each cluster. (C) T cell subsets are shown by patient. (D) Bar graphs reveal the percentage of T cell subsets from the total cell population. (E) Gene set enrichment analysis (GSEA) shows HALLMARK pathways upregulated in the DOCK4-CD4 cluster compared with all other clusters pooled. (F) UMAP plot showing the distribution of cancer-associated fibroblast (CAF) colored by subtype in HER2-0 and HER2-low samples. (G) The average absolute number (top) and the proportion of the CAF subtype (bottom). (H) Gene set enrichment analysis (GESEA) reveals tissue remodeling associated pathways from Gene ontology biological process (GOBP), HALLMARKS, and REACTOME upregulated in CAFs from HER2-0 compared to HER2-low. (I) Circle plot showing a differential number of interactions (above left) and differential interaction strength (above right) in the cell-cell communication network of HER2-0 vs. HER2-low tumors. The circle color is coded according to the cell type, and the line color indicates the cell source as the sender. Red-colored edges represent increased signaling, and blue-colored edges represent decreased signaling in HER2-0 compared to HER2-low. A thicker line represents the number (above left) or the strength (above right) of interactions. The heatmaps show the number (lower left) and strength (lower right) of differential interactions in HER2-0 vs. HER2-low. The top-colored bar plot represents the sum of each column of the absolute values displayed in the heatmap (incoming signaling). The right-colored bar plot represents the sum of each row of the absolute values (outgoing signaling). The bar height indicates the degree of change regarding the number of interactions or strength between the two conditions. In the color bar, red and blue represent increased and decreased signaling in HER2-0 compared to HER2-low. (J-K) Cyclic immunofluorescence (CyCIF) was performed (HER2 low, n=15; HER2-0, n=5). (J) Representative images, DNA (blue), αSMA (orange), CD31 (white), CD4 (purple), and CD68 (green) staining. Scale bars, 100 µm in the insets. (K) Bar graph representing the global percentage of endothelial cells (top) and CAFs (bottom) from the total cells. The two-tailed t-test and the Wilcoxon rank-sum test were used for statistical analysis (p<0.05).

### Cancer-associated fibroblasts may support tissue remodeling in HR+ HER2-0 breast tumors

As we had identified differences observed in the myeloid and lymphoid immune populations between HER2-0 and HER2-low, we next sought to understand how stromal components may differ between tumor types. Cancer-associated fibroblasts (CAFs) are key stromal cells that remodel the extracellular matrix (ECM) and promote tumor progression. In breast cancer, CAFs promote ECM remodeling and facilitate angiogenesis(16, 17). To investigate the contribution of CAFs, we subclustered these cells from the total cell population and identified eight CAF clusters (**Supplementary Fig. 10A-B** and **Fig 4F-G**). C4-CAFs were enriched in HER2-0 tumors compared to HER2-low samples as a frequency out of CAFs (20.4% vs. 7.43%, p=0.03) as well as out of total cells (3.56% vs. 1.2%, p=0.01) (**Supplementary Fig. 10C)**. The C4-CAF cluster was characterized by the expression of COL1A1 and COL3A1, suggesting a role in wound-repair, (**Supplementary Fig. 10C**). Gene set enrichment analysis further supported this phenotype as multiple pathways involved in extracellular matrix (ECM) remodeling (padj=0.001), response to wounding (padj=0.001), and blood vessel morphogenesis (padj=0.001) were enhanced in CAFs from HER2-0 tumors compared to HER2-low (**Fig. 4H**).

To assess the interactions between the different tumor cell populations, we next analyzed potential cell networks within HER2-0 and HER2-low tumors using the CellChat package (18). We observed significant interactions in both the number and strength of interactions between CAFs and endothelial cells, both of which were higher in HER2-0 tumors compared to HER2- low (**Fig. 4I**). Furthermore, comparing the sum of communication probabilities among all pairs of cell groups in the inferred network, we observed that markers of ECM remodeling (*EGF* and *FN1),* vascular development (*EPHA* and *EPHB*), and immune response (CD45 and complement) were higher in HER2-0 compared to HER2-low (**Supplementary Fig. 10D-E)**. However, using CyCIF we observed that the frequency of endothelial cells (CD31+) and CAFs (αSMA+) was similar in both groups, suggesting that the difference lies not in abundance but in specialized biological functions (**Fig. 4J-K)**. Together, these results indicated that CAFs potentially contribute to tissue remodeling in HR+ HER2-0 breast tumors.

## Discussion

The present study highlights the key immune and stromal differences between treatment-naive HER2-0 and HER2-low, HR+ tumors. HER2-0 tumors exhibited prominent inflammatory and tissue remodeling signatures, whereas HER2-low tumors showed enhanced metabolic activity, cell cycle progression, and estrogen signaling, indicating active cellular proliferative programs.

Recent analyses of large retrospective cohorts have highlighted the differences in genetic and tumor immune features between HR+/HER2-0 and HR+/HER2-low breast cancer tumors(7–9, 19). In a cohort of 973 patients, among the HR+ tumors, HER2-0 and HER2-low had a similar density of TIL whereas within HR- tumors, HER2-0 was associated with an increased number of TIL compared with HER2-low tumors (7). In a separate cohort of 529 patients, HR+ HER2-0 tumors demonstrated increased expression of genes associated with an adaptive immune response compared with HER2-low tumors(9). Similarly, we observed that tumor cells, TAMs, and CAFs from HER2-0 samples were significantly enriched in genes related to immune signaling and responses compared to the HER2-low cohort. Furthermore, our study revealed a significant accumulation of APOE-TAMs in HER2-0 tumors. APOE expression has been detected in a subset of TAMs found across several cancers and is defined as lipid-associated macrophages(14). The accumulation of lipids or lipid-induced pathways in macrophages is associated with pro-tumor activities supporting tumor invasiveness and progression(14). In murine models of gastric carcinoma, colorectal carcinoma, and hepatocellular carcinoma, pharmacological inhibition or genetic deletion of APOE potentiated anti-PD1 therapy (20). As DOCK4-T cells with cytotoxic potential are associated with APOE-TAMs in HER2-0 breast cancer, it is possible that strategies aimed at targeting APOE-TAMs may enhance anti-tumor T cell responses.

Chronic inflammatory processes, tissue remodeling, and stroma modification in solid cancers are recognized as essential for cancer development and progression(21). CAFs, the primary stromal cells of solid tumors, play a central role in the multifactorial network involved in each one of these processes(22). In breast cancer, CAF-mediated collagen cross-linking has been shown to stiffen the ECM and promotes the invasion of a premalignant lesions(16). Another study, using a 3D *in vitro* model of blood vessel formation, it was demonstrated that CAFs derived from human breast tumors support the development of vasculature independently of soluble factors, including VEGF(17). CAFs generate more significant deformations within the ECM, stimulating the self-assembly of endothelial cells into a vascular network(17). Our data revealed that CAFs from HER2-0 tumors showed enrichment of pathways involved in ECM remodeling. Particularly, the pathways involved in collagen ECM synthesis were upregulated in HER2-0 compared to HER2-low tumors. These observations suggest that CAFs support a tissue remodeling program in HR+ HER2-0 breast tumors.

Crosstalk between HER2 and ER pathways has been implicated in tumor growth, immune suppression, and therapy resistance(23). This interaction occurs through common downstream signaling pathways such as the PI3K/AKT pathways, which is key regulators of cellular proliferation and immune evasion (23, 24). Notably, most ER-high expressing tumors are HER2- low(6), supporting the notion of an interplay between these pathways with a potential impact on the TME immune composition. Our group previously demonstrated that increased ER signaling in cancer cells was negatively associated with immune-related gene modules such as TNFα/NF- κB signaling and type-I interferon (IFN-I) response(25). Here, we observed that even among tumors with high ER expression (>50%), HER2-low tumors exhibited an enrichment of pathways associated with early and late ER responses compared to HER2-0 tumors. These data suggest that the ER/HER2 pathway may contribute to the suppression of the immune response observed in HR+ HER2-low tumors.

Our study has limitations, including its retrospective design and the unequal distribution of patients between HER2 groups. Furthermore, the present study focused on early-stage breast cancer, and therefore, our conclusions cannot be applied to a metastatic setting. Despite these limitations, the strength of this study lies in its comprehensive, multi-modal approach to deeply characterize the immune and stromal complexity of HER2-0 and HER2-low breast cancer, providing valuable insights into the distinct tumor microenvironments of these emerging subtypes.

Taken together, our findings offer a comprehensive view of the tumor landscapes in HR+ HER2- low and HER2-0 breast cancer. HER2-0 tumors are characterized by the enrichment of inflammatory and tissue remodeling pathways, along with marked infiltration of APOE+ macrophages and DOCK4-CD4 T cells, suggesting a more active and potentially targetable immune microenvironment. In contrast, HER2-low tumors exhibit increased ER signaling and metabolic and proliferative programs. These insights deepen our understanding of the distinct biology underlying the HR+ HER2-low and HER2-0 subtypes and may inform therapeutic strategies.

## Methods

### Cohort characteristics

Breast tumors used in this study were collected with written informed consent from all patients under DFCI protocol 93-085 with approval from all relevant human research ethics committees (Dana-Farber Cancer Institute Ethics Committee). Consent included the use of all de-identified patient data for publication, which was carried out in accordance with the Declaration of Helsinki. Fresh tissues were split in half with a sterile razor and either placed in a cryovial for immediate flash freezing in liquid nitrogen or placed in 10% neutral buffered formalin for formalin fixation for 24 hours prior to paraffin embedding. A total of 25 patients with primary, treatment-naïve, HR+ breast cancer was included in this study. Estrogen receptor, progesterone receptor, and HER2 expression were confirmed by IHC and scored by a clinical pathologist. HER2 status was determined by IHC and, when applicable, by FISH. Additionally, three HER2-positive tumors (IHC 3+ or IHC 2+/FISH-positive) were analyzed as reference samples for ERBB2 gene expression only. Of the patients assessed, one with HER2–ultra-low disease was excluded. Age, menopausal status, and histological grade were well balanced between the HER2-low and HER2-0 groups. Detailed patient demographics and clinicopathologic characteristics according to HER2 status are provided in **Supplementary Table 1**.

### Bulk RNA-seq analysis

RNA was extracted from 5-micron thick FFPE sections using the High Pure FFPET RNA Isolation Kit (Roche, 424530). The RNA was quantified using the Nanodrop One instrument (Thermo Fisher Scientific). RNA was sequenced using 2 x 150 bp reads on an Illumina NovaSeq sequencer. Sequencing reads were aligned to the human reference genome GRCh38 using STAR and hit counts were generated.

### Immune cell infiltration

To estimate the relative abundance of immune cell populations within the bulk RNA-seq profiles of HER2-low and HER2-0 breast tumors, we utilized the TIMER2.0 web platform (Tumor Immune Estimation Resource, http://timer.cistrome.org). Gene expression data from 25 tumor samples (17 HER2-low and 8 HER2-0) were uploaded to TIMER2.0 for deconvolution analysis. We employed the "Immune Estimation" module to infer the infiltration levels of major immune cell types, including CD4⁺ T cells, CD8⁺ T cells, B cells, neutrophils, macrophages, and dendritic cells, using the TIMER algorithm.

### Single-nuclei GEM generation and gene expression library construction

Low-retention microcentrifuge tubes (Fisher Scientific, Hampton, NH, USA) were used throughout the procedure to minimize nuclei loss. Briefly, tissues were manually dissociated by chopping with fine spring scissors for 10 minutes, homogenized in TST solution, filtered through a 30-µm MACS SmartStrainer (Miltenyi Biotec, Germany), and pelleted by centrifugation for ten minutes at 500g at 4°C. Nuclei pellets were washed with 1 mL of resuspension buffer (1X PBS + 1% BSA) and filtered again, centrifuged at 500g for 5 minutes at 4°C and then resuspended in 100 µL of resuspension buffer and trypan blue-stained nuclei were counted manually using INCYTO C-Chip Neubauer Improved Disposable Hemacytometers (VWR International Ltd., Radnor, PA, USA) under a brightfield microscope. This process was performed for each sample independently. The yield and quality of nuclei were adequate to process further for single-nuclei RNA sequencing. A droplet-based microfluidic Chromium Controller (10X Genomics, Inc., Pleasanton, CA, USA) was used to generate single-nuclei Gel Beads-in-emulsions (GEMs), as per the Chromium Single Cell 5’ Reagent Kits (Cat no. 1000263) User Guide RevE (v2 Chemistry Dual Index; 10X Genomics demonstrated protocol). Approximately 10,000 nuclei were loaded with 10X-barcoded gel beads and partitioning oil including RT master mix for GEM generation. Next, GEMs were incubated to produce 10x barcoded, full-length double-stranded cDNA from polyadenylated mRNA. The Agilent High Sensitivity DNA Kit assessed the resulting cDNA with the Agilent Bioanalyzer 2100 (Agilent Technologies, Lexington, MA, USA). Approximately 50 ng of cDNA were processed for gene expression library construction. and libraries were also quantified on the Agilent Bioanalyzer 2100. Next, libraries were assessed on the Agilent Bioanalyzer 2100. All the libraries were normalized and pooled for sequencing on Illumina NovaSeq platforms (Illumina, Inc., San Diego, CA, USA).

### Single-nuclei RNAseq (nucSeq) data pre-processing

Pooled samples were demultiplexed using 10x Cellranger mkfastq. Read pairs were aligned to the hg38 reference sequence using a 10x Cellranger count, which utilizes STAR. Cellbender remove-background was run on aligned, barcoded reads to distinguish cell-containing droplets from empty droplets and retrieve background-free gene expression profiles. Cellbender-filtered matrices were analyzed in RStudio using Seurat v4.4.0. Here, data was further filtered by retaining cells with ≥ 200 detected genes, and genes if they were expressed in ≥ 3 cells. Cells with < 20% mitochondrial read were retained. All samples were merged into a single Seurat object and cells were integrated using Harmony. Dimensionality reduction was performed using the Uniform Manifold Approximation and Projection (UMAP) algorithm from matplotlib to cluster cells and preserve similarities in the original gene expression space. Individual clusters were annotated using canonical marker expression.

### Differential gene expression, module, and cellular interaction prediction analysis

Differential gene expression analysis between HER2-0 and HER2-low cells was performed using the FindAllMarkers function within Seurat v4.4.0. Gene set enrichment analysis was performed using differentially expressed genes and the Hallmark, Gene Ontology (GO), and Reactome gene sets derived from the “msigdbr” R package. Cellular interaction analysis was performed using the “CellChat” R package using clusters derived from Seurat v4.4.0.

### Profiling macrophage phenotypic states

To classify TAMs subtypes, we used gene signatures defined by Ma et al., 2022 (13) encompassing Nonclassical monocytes (Nonclassical mono), Intermediated monocytes (Intermediated mono), Classical tumor infiltrating monocytes (Classical TIM), Resident tissue macrophages (RTM TAMS), Regulatory TAMs (Reg TAMs), Proliferative TAMs (Prolif TAMs), Inflammatory TAMs (Inflamma TAMs), Interferon TAMs (IFN TAMs), Lipid associated TAMs (LA TAMs), and Angiogenic TAMs (angio TAMs). These signatures were compiled into a gene set list and used to annotate clusters identified from our Seurat object, which was previously batch- corrected using harmony. Differential expression analysis across clusters was performed using the wilcoxauc function and ranked gene lists were generated based on log fold change and AUC. Gene set enrichment analysis (GSEA) was performed using the fgsea package (v1.3.0). The enrichment of TAM subtypes per cluster was further visualized using radar plots generated with the fmsb and RColorBrewer packages in R.

### Copy number variation inference

Copy number variations (CNVs) were inferred from single-cell RNA sequencing data using the inferCNV (v1.23.0) tool (https://github.com/broadinstitute/inferCNV), executed via the Terra platform (Broad Institute). This analysis focused on detecting large-scale CNVs across the genome, particularly on chromosome 17, which harbors the *ERBB2* (HER2) gene locus. Expression data from tumor cells were compared with non-malignant immune T cells as reference populations to identify tumor-specific CNV patterns. The inferCNV workflow was configured with standard parameters (cutoff = 0.1, cluster_by_groups = TRUE, and denoise = TRUE). Heatmaps were generated to visualize the CNV profiles, highlighting amplifications and deletions, particularly within the 17q12 region. This approach allowed us to infer *ERBB2* amplification status and assess chromosomal instability in tumor subpopulations.

### Cyclic immunofluorescence (CYCIF) staining

CyCIF was performed on 5-micron thick FFPE sections. Briefly, the sections were dewaxed and underwent antigen retrieval using a Leica Bond Rx instrument. Sections were then blocked with SuperBlock Blocking Buffer (Thermo Fisher; 27515) containing secondary antibodies for all primary antibody host species used in the staining panel overnight at 4C. Sections were mounted to coverslips using 70% glycerol and imaged using the CyteFinder II (RareCyte) whole-slide imager at 20X magnification. After imaging, the slides were de-coverslipped in PBS and photobleached using sodium hydroxide and hydrogen peroxide in PBS for 30 minutes under LED light. Staining, imaging, and bleaching were performed repeatedly for the subsequent cycles until completed.

### Cyclic immunofluorescence (CYCIF) data pre-processing

Whole slide images for each cycle were overlaid, and cells were segmented using the MCMICRO computational platform. Overlaid whole slide images were visualized using OMERO and regions of interest (ROI) were selected for further analysis based upon segmented cell presence at the final cycle. Fluorescence intensity for each marker within ROIs was collected for each segmented cell using Gater (https://github.com/labsyspharm/gater).

## Data Availability Statement

The transcriptomic and single-nucleus RNA sequencing (sNuc-seq) data generated in this study have been deposited in the Gene Expression Omnibus (GEO) under accession number. All custom code used for data processing and analysis is available on GitHub. Additional data supporting the findings of this study are available from the corresponding author upon reasonable request.

## Statistics

Clinical and baseline genomic categorical variables were compared between the HER2-low and HER2-0 groups using the chi-square or Fisher’s test when appropriated. For differential gene expression analysis within individual clusters, either a two-sided Student’s *t*-test or the non- parametric Wilcoxon rank-sum test was applied, depending on the data distribution. Comparisons involving more than two groups (e.g., across multiple cell types or sample categories) were conducted using one-way analysis of variance (ANOVA), followed by Tukey’s post-hoc test to adjust for multiple comparisons. Correlation analyses between continuous variables were performed using Pearson’s correlation coefficients. A *p*-value of <0.05 was considered statistically significant unless otherwise specified. All statistical analyses were conducted using R (version 4.4.1) or GraphPad Prism (version 10.4.2), as appropriate

## Author contributions

Concept and design: C.W.S.W., D.M., Acquisition, analysis, or interpretation of data: C.W.S.W., D.M., A.N., K.S., J.F., R.B-S., A.W., P.T. J.L.G., Drafting of the manuscript: C.W.S.W., Critical revision of the manuscript for important intellectual content: D.M., K.S., A.N., S.M.T., E.A.M, R.B- S., A.W., P.T., S.J.S, J.L.G. Statistical analysis: C.W.S.W., D.M., K.S. Obtained funding: J.L.G. Administrative, technical, or material support: A.W., Supervision: P.T., J.L.G.

## Acknowledgments

This work was supported by the Susan G. Komen Foundation (ASPIRE1039903, CCR18547597; Fundação de Amparo à Pesquisa do Estado de São Paulo (FAPESP 2021/12898-8), The Harvard Ludwig Center, NIH DF/HCC SPORE in Breast Cancer (P50 CA168504), NIH NCI (R37 CA269499), and The Concern Foundation. A.N. was supported by the Canadian Institute of Health Research Postdoctoral Fellowship (FRN: 194068). The authors used AI tools to assist with English language editing and stylistic improvements. The scientific content and interpretation remain the responsibility of the authors.

