## Supplemental figures for "APOE+ Tumor-Associated Macrophages and CD4-DOCK4 T Cells Reveal Distinct Microenvironmental Features in HER2-Low and HER2-0 Hormone Receptor-Positive Breast Cancer"

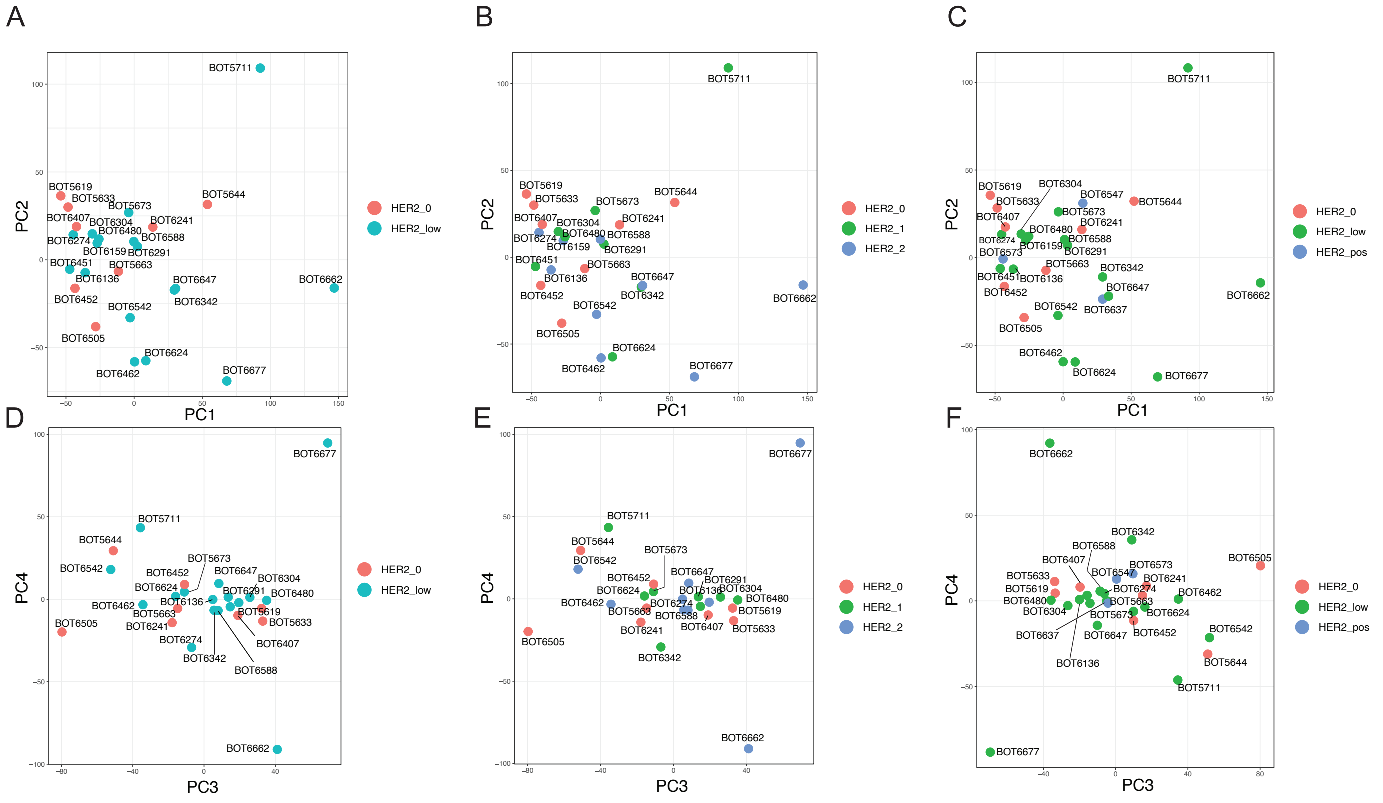


**Supplementary Fig. 1. Principal component analysis (PCA) of RNA-sequencing of bulk tumor from HR+ HER-2-negative breast tumors.** Each dot represents an individual tumor, colored according to HER2 status. (A and D) HER2-0 and HER2-low. (B and E) HER2-0, HER2-1+ and HER2-2+. (C and F) HER2-0, HER2-low, and HER2-positive.

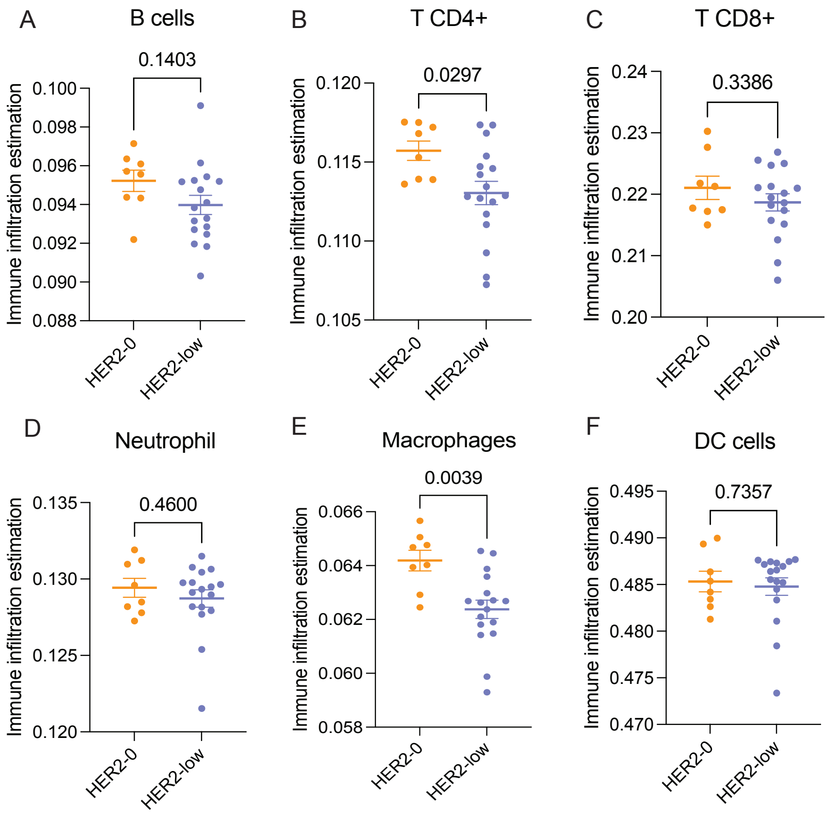


**Supplementary Fig. 2. Immune cell subsets across HR+ HER2-0 and HER2-low breast tumors were analyzed using TIMER.** The proportion of immune cell subsets are show. Each dot represents an individual tumor. Statistical analysis was performed using a two-tailed t-test, with significance set at $p < 0.05$.

A

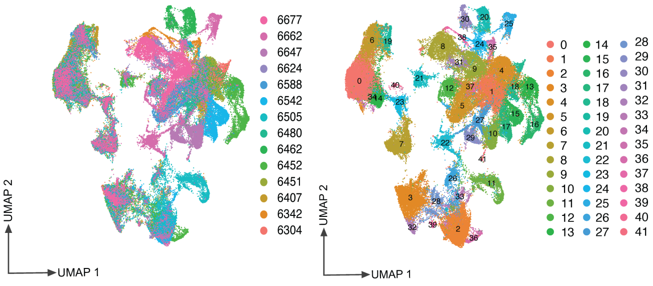


B

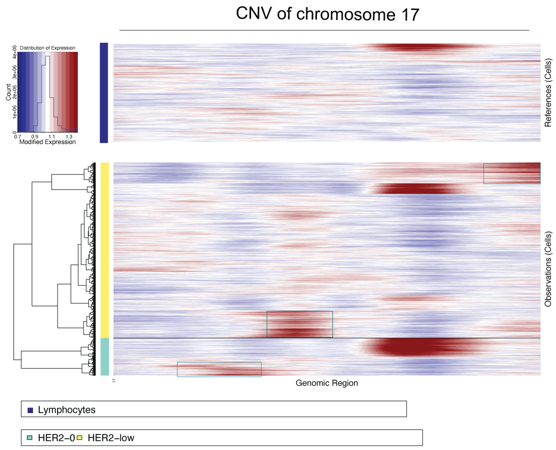


C

| | CNV name | CNV state | CNV event | Chr | Start | End |
|---|---|---|---|---|---|---|
| HER2-0 | *17-region_1* | 1 | Loss | 17 | 212389 | 7561608 |
| | *17-region_3* | 6 | Gain | 17 | 15436015 | 17837002 |
| | *17-region_4* | 6 | Gain | 17 | 17843511 | 31538217 |
| | *17-region_5* | 1 | Loss | 17 | 31860899 | 38729803 |
| | *17-region_7* | 1 | Loss | 17 | 44034635 | 48604912 |
| | *17-region_9* | 6 | Gain | 17 | 50183289 | 50692252 |
| | *17-region_10* | 6 | Gain | 17 | 50693190 | 75825799 |
| | *17-region_11* | 6 | Gain | 17 | 75845699 | 76586956 |
| | *17-region_12* | 6 | Gain | 17 | 76672551 | 83095119 |
| HER2-low | *17-region_13* | 1 | Loss | 17 | 212389 | 8156506 |
| | *17-region_15* | 1 | Loss | 17 | 28614440 | 36543435 |
| | *17-region_17* | 1 | Loss | 17 | 38733897 | 39564907 |
| | *17-region_18* | 6 | Gain | 17 | 39637065 | 40357643 |
| | *17-region_19* | 6 | Gain | 17 | 40388516 | 42676994 |
| | *17-region_20* | 6 | Gain | 17 | 42700275 | 43833170 |
| | *17-region_21* | 6 | Gain | 17 | 44011188 | 44353102 |
| | *17-region_23* | 1 | Loss | 17 | 51153536 | 60078931 |
| | *17-region_25* | 1 | Loss | 17 | 63745250 | 75182983 |
| | *17-region_27* | 6 | Gain | 17 | 76735865 | 80147183 |
| | *17-region_28* | 6 | Gain | 17 | 80169992 | 83095119 |

**Supplementary Fig. 3. Single-Nucleus transcriptomic landscape and chromosome 17 CNV profiling in HER2-0 and HER2-Low breast tumors.** (A) UMAP visualization of 90,248 single nuclei (14 individuals) (left). A UMAP plot showing the distribution of 42 clusters (right). (B) Heatmap showing the inferred copy number variations (CNV) profile of chromosome 17 in HER2-0 and HER2-low epithelial cells. Lymphocytes were used as a reference. (C) Table depicting CNVs in chromosome 17 with their copy number states and precise genomic locations.

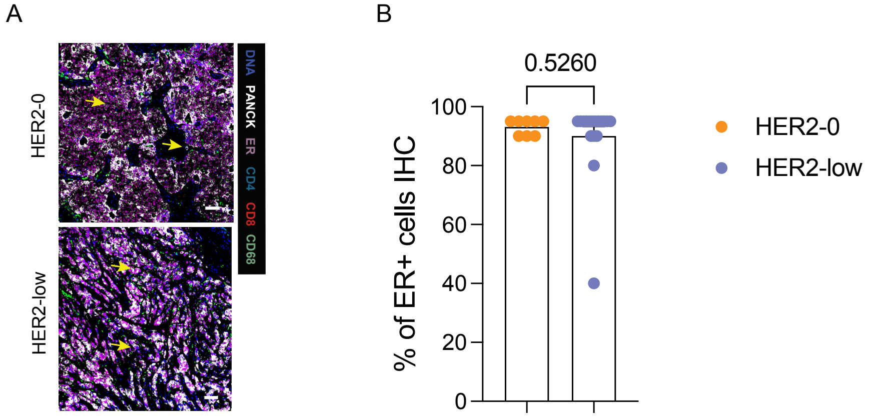


**Supplementary Fig. 4. ER expression in HER2-0 and HER2-low tumors. (A)** Representative images of breast tumors in which cyclic immunofluorescence (CyCIF) was performed to assess ER expression in tumors. DNA (Blue), PanCK (white), ER (purple), CD4 (light blue), CD8 (red), and CD68 (green) staining. Scale bars, 50 µm in the insets, 2 mm in the whole section image. **(B)** The percentage of ER-positive tumor cells in HER2-0 and HER2-low patients are shown. Statistical analysis was performed using a two-tailed t-test, with significance set at $p < 0.05$.

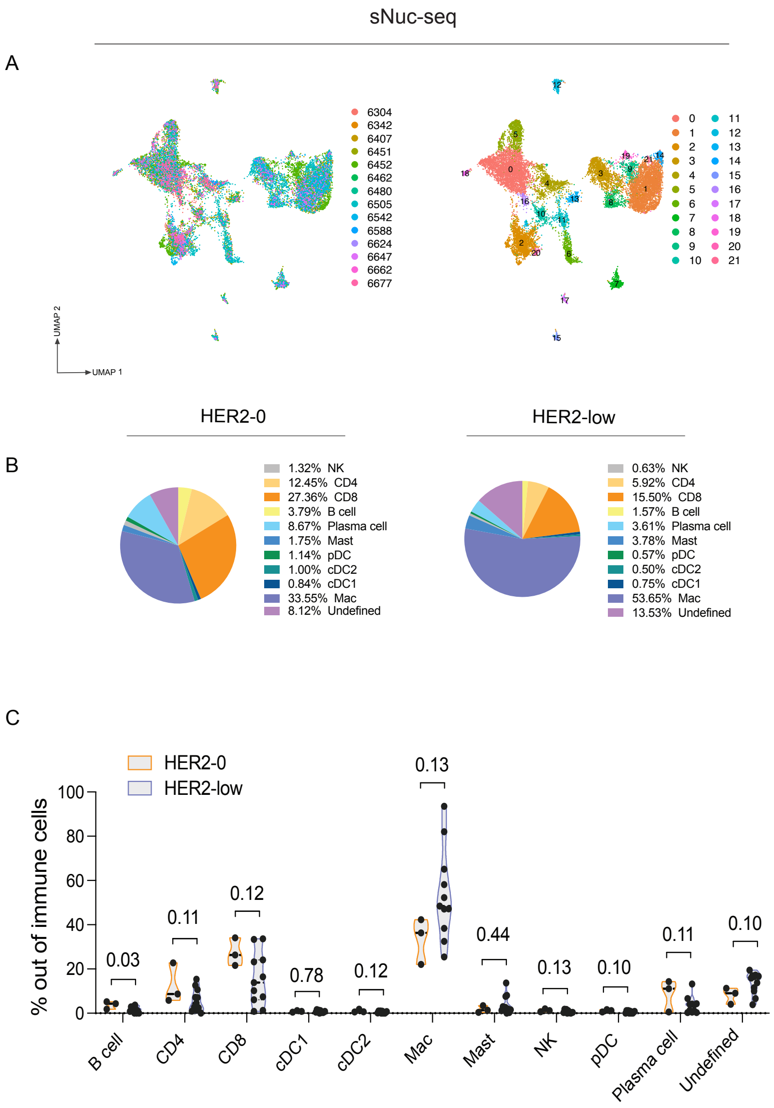


**Supplementary Fig. 5. Immune cell subsets derived from to sNuc-seq.** (A) UMAP plots show immune cell (PTPRC+) subpopulations in subcluster analysis, according to the sample source (left) and clustering distribution (right). (B) The average of immune cell subtype proportion out of immune cells (PTPRC+). (C) The percentage of immune cell subsets out of total cells are shown. Statistical analysis was performed using a two-tailed t-test, with significance set at $p < 0.05$.

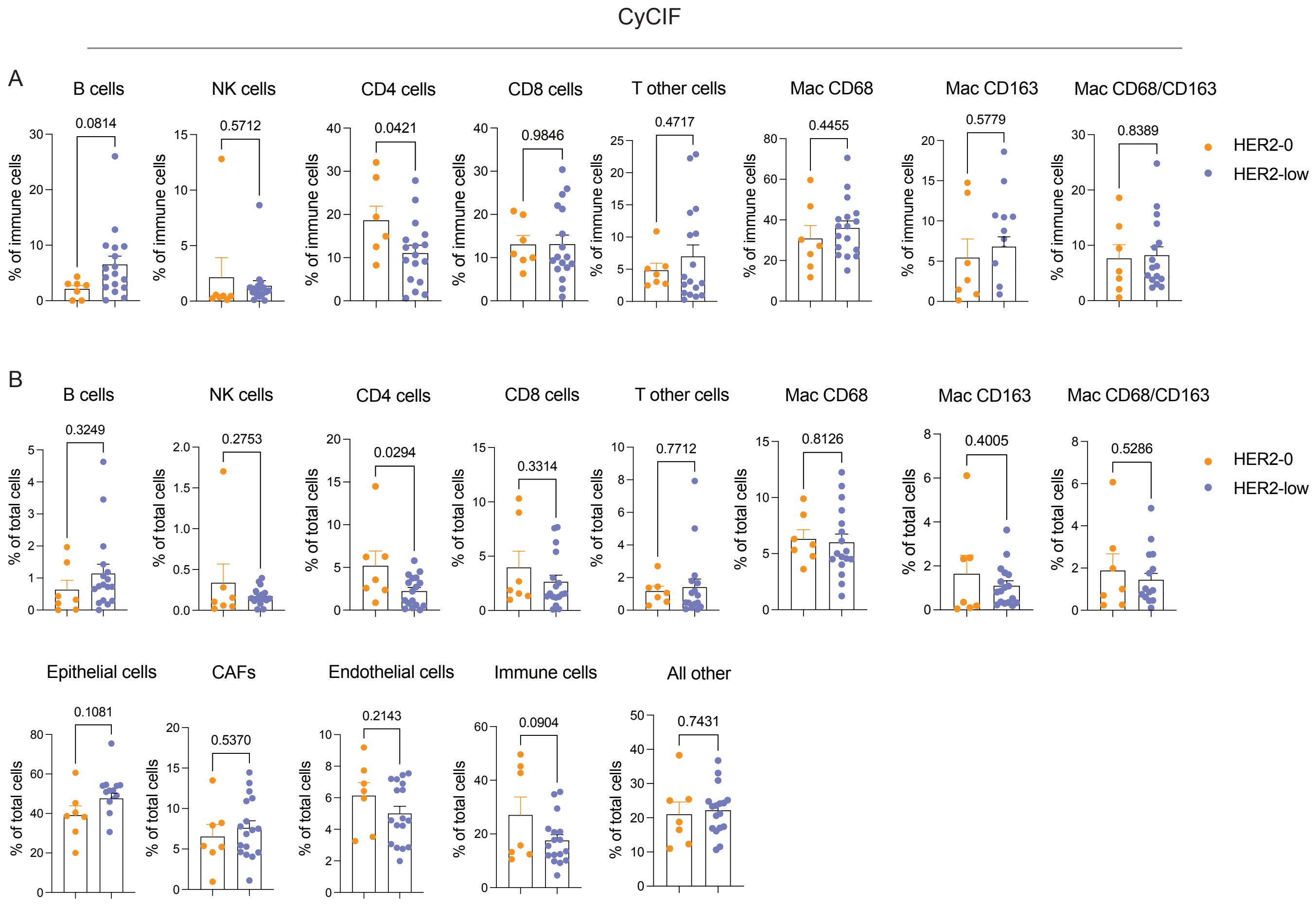


**Supplementary Fig. 6. Immune cell subsets derived from CyCIF.** (A) The frequency of immune cell subsets out of CD45+ cells is shown. (B) Percent of immune cell subsets out of the total of cells. Statistical analysis was performed using a two-tailed t-test, with significance set at $p < 0.05$.

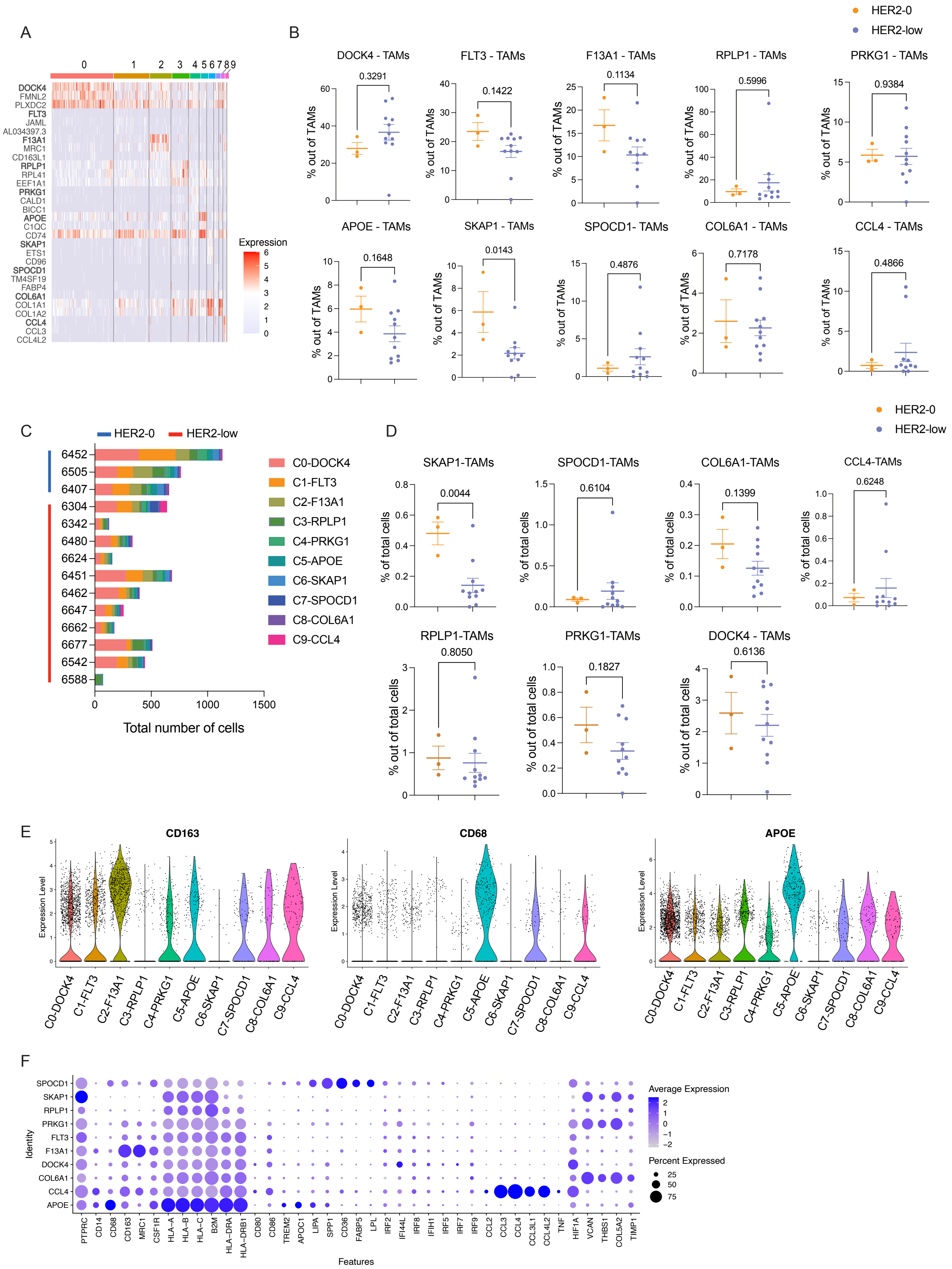
A
B
C
D
E
F
HER2-0
HER2-low
DOCK4 - TAMs
FLT3 - TAMs
F13A1 - TAMs
RPLP1 - TAMs
PRKG1 - TAMs
APOE - TAMs
SKAP1 - TAMs
SPOCD1- TAMs
COL6A1 - TAMs
CCL4 - TAMs
0.3291
0.1422
0.1134
0.5996
0.9384
0.1648
0.0143
0.4876
0.7178
0.4866
% out of TAMs
Expression
DOCK4
FMNL2
PLXDC2
FLT3
JAML
AL034397.3
F13A1
MRC1
CD163L1
RPLP1
RPL41
EEF1A1
PRKG1
CALD1
BICC1
APOE
C1QC
CD74
SKAP1
ETS1
CD96
SPOCD1
TM4SF19
FABP4
COL6A1
COL1A1
COL1A2
CCL4
CCL3
CCL4L2
6452
6505
6407
6304
6342
6480
6624
6451
6462
6647
6662
6677
6542
6588
Total number of cells
C0-DOCK4
C1-FLT3
C2-F13A1
C3-RPLP1
C4-PRKG1
C5-APOE
C6-SKAP1
C7-SPOCD1
C8-COL6A1
C9-CCL4
SKAP1-TAMs
SPOCD1-TAMs
COL6A1-TAMs
CCL4-TAMs
0.0044
0.6104
0.1399
0.6248
% of total cells
RPLP1-TAMs
PRKG1-TAMs
DOCK4 - TAMs
0.8050
0.1827
0.6136
% out of total cells
CD163
CD68
APOE
Expression Level
Identity
Features
Average Expression
Percent Expressed
PTPRC
CD14
CD68
CD163
MRC1
CSF1R
HLA-A
HLA-B
HLA-C
B2M
HLA-DRA
HLA-DRB1
CD80
CD86
TREM2
APOC1
LIPA
SPP1
CD36
FABP5
LPL
IRF2
IFI44L
IRF8
IFIH1
IRF5
IRF7
IRF9
CCL2
CCL3
CCL4
CCL3L1
CCL4L2
TNF
HIF1A
VCAN
THBS1
COL5A2
TIMP1

**Supplementary Fig. 7. Fractions of TAM subsets derived from sNuc-seq.** (A) Heatmap with top 3 marker genes for each cluster of TAMs. (B) The frequency of TAM subsets out of all total amount of TAMs. (C) Total number of TAMs by subtypes shown for each patient. (D) Graph representing the percentage of TAM subsets out of total cells. (E) Violin plots represent *CD163*, *CD68*, and *APOE* expression in TAM clusters. (F) Dot plot showing select TAMs markers gene expression values (log scale) and percentage of nuclei expressing these genes within each cluster. Statistical analysis was performed using a two-tailed t-test, with significance set at $p < 0.05$.

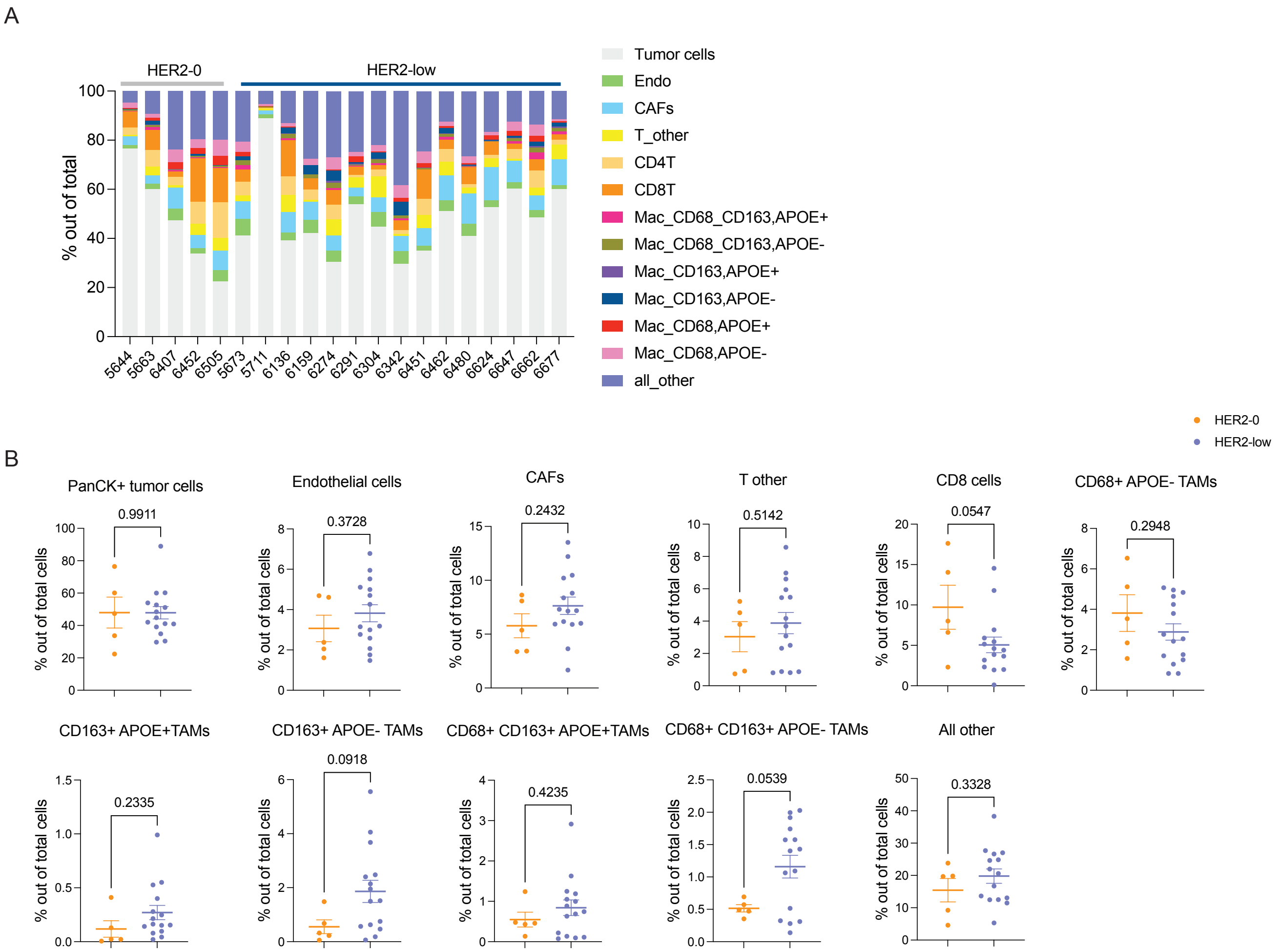


**Supplementary Fig. 8. Proportions of TAMs subsets derived from CyCIF analysis.** (A) The proportion of cells out of the total cells in the tumor are plotted by patient. (B) Graph representing the percentage of tumor cells (PanCK+), endothelial (CD31+), CAFs (αSMA), CD8 T cells and T other (CD3+, CD4- and CD8-) and TAM subsets (CD68+, CD163+, or CD68+/163+, APOE+ or APOE-) and  out of total cells. Statistical analysis was performed using a two-tailed t-test, with significance set at $p < 0.05$.

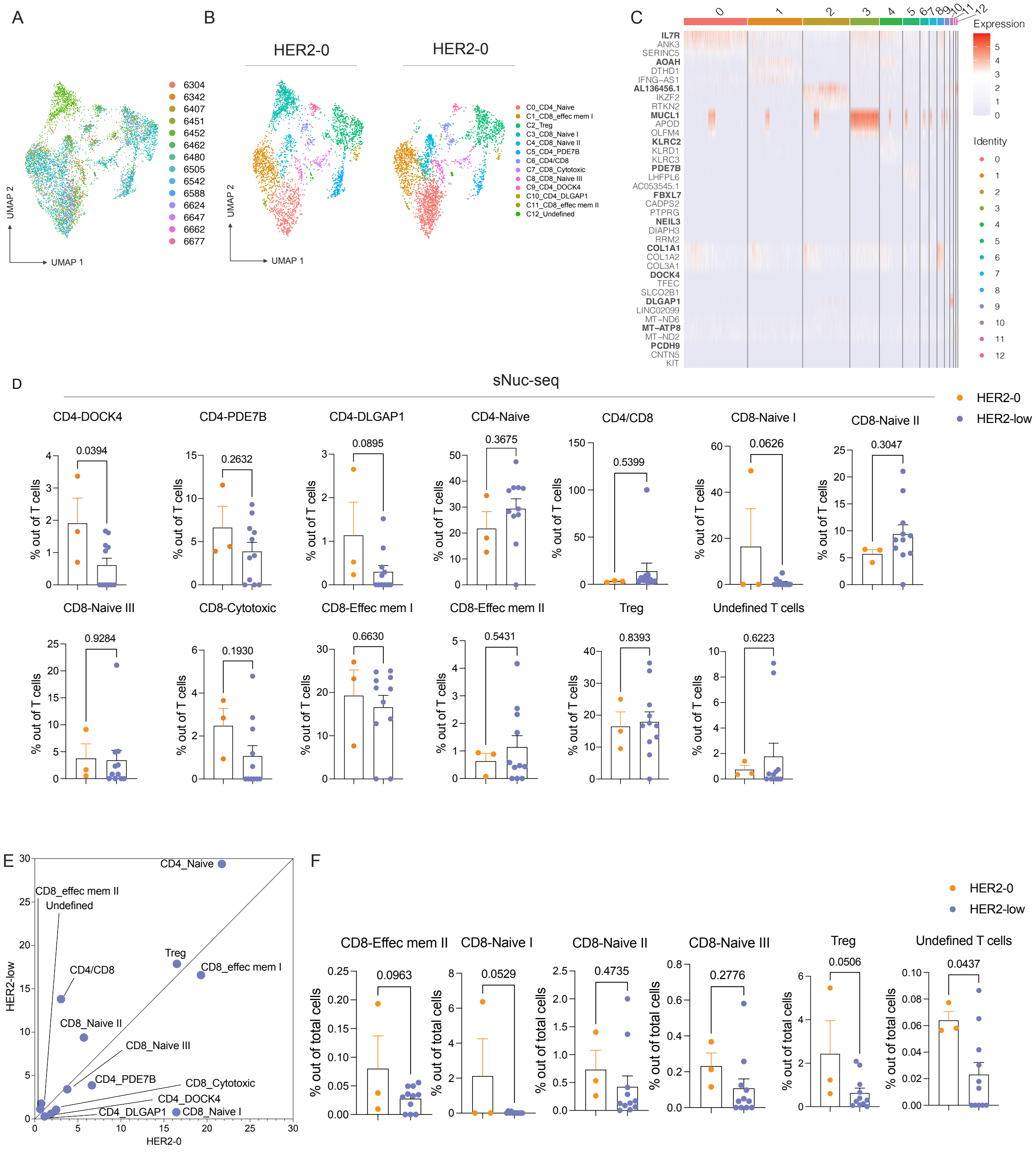


**Supplementary Fig. 9. Frequency of lymphocyte subsets derived from sNuc-seq.** (A) UMAP plots show CD4 and CD8 T cell subsets according to the sample source. (B) UMAP plot showing the distribution of T cells in HER2-0 and HER2-low samples. (C) Heatmap with top 3 marker genes for each CD4 and CD8 lymphocyte cluster. (D) Frequency of T cell subsets out of total CD4 and CD8 cells are shown. (E) Frequency of CD4 and CD8 T cells by HER2-0 (x-axis)

or HER2-low (y-axis) tumors. (F) Frequency of T cell subsets out of the total cells. Statistical analysis was performed using a two-tailed t-test, with significance set at $p < 0.05$.

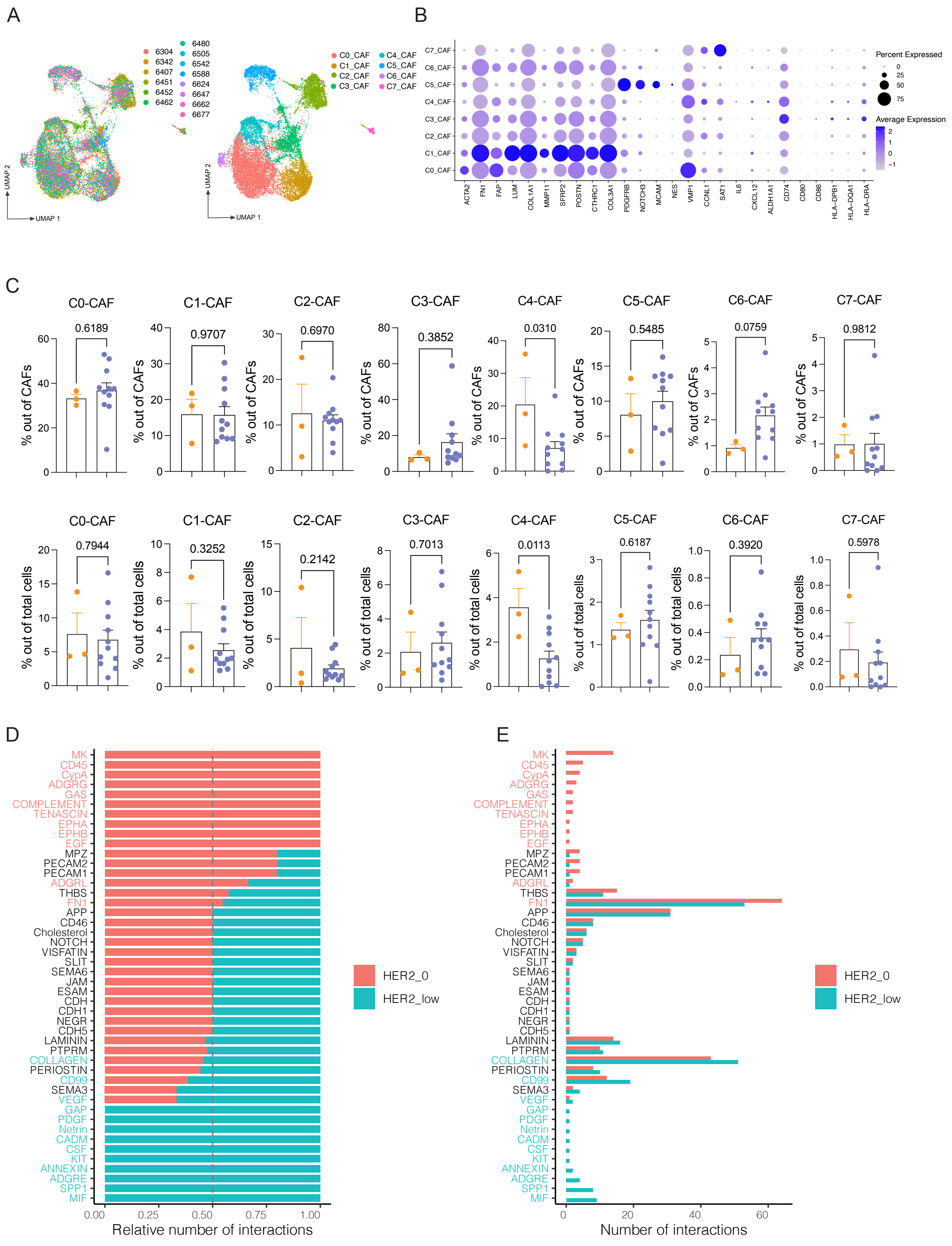

A
B
C
D
E
UMAP 1
UMAP 2
Percent Expressed
Average Expression
C0-CAF
C1-CAF
C2-CAF
C3-CAF
C4-CAF
C5-CAF
C6-CAF
C7-CAF
% out of CAFs
% out of total cells
0.6189
0.9707
0.6970
0.3852
0.0310
0.5485
0.0759
0.9812
0.7944
0.3252
0.2142
0.7013
0.0113
0.6187
0.3920
0.5978
Relative number of interactions
Number of interactions
HER2_0
HER2_low

**Supplementary Fig. 10. Frequency of cancer-associated fibroblast (CAF) subsets derived from sNuc-seq and analyses of communications between cell populations.** (A) UMAP plots show CAF subpopulations shown by patient (left), and UMAP clusters are labeled with inferred CAF types (right). (B) Dot plot showing select CAF markers gene expression values (log scale) and percentage of nuclei expressing these genes within each cluster. (C) The frequency of CAF subsets out of the total number of CAFs (above). The frequency of CAF subsets out of the total cells (below). (D-E) Bar graphs represent the significant signaling pathways, ranked based on differences in the overall information flow within the inferred networks between HER2-0 and HER2-low. The top signaling pathways enriched in HER2-0 (red) and enriched in the HER2-low (blue) are shown. Statistical analysis was performed using a two-tailed t-test, with significance set at $p < 0.05$.
