## Supplemental tables for "APOE+ Tumor-Associated Macrophages and CD4-DOCK4 T Cells Reveal Distinct Microenvironmental Features in HER2-Low and HER2-0 Hormone Receptor-Positive Breast Cancer"

**Supplementary table 1. Patient and tumor characteristics among patients with breast cancer include**

| | Total population (N=25) | Patients with HER-0 tumors tested (N=8) | Patients with HER-low Tumors tested (N=17) | *P*-Value |
|---|---|---|---|---|
| ***Age in years,*** *median (min, max)* | 52.4 (32, 84) | 54.87 (38, 75) | 51.2 (32, 84) | 0.557* |
| ***Sex,*** *n (%)* | - | - | - | >0.999# |
| Female | 24 (96) | 8 (100) | 16 (94.12) | - |
| Male | 1 (4) | - | 1 (5.88) | - |
| ***Ethnicity,*** *n (%)* | - | - | - | >0.999# |
| African American/Black | 1 (4) | - | 1 (5.88) | |
| Asian or Pacific Islander | 1 (4) | - | 1 (5.88) | |
| Caucasian | 23 (92) | 8 (100) | 15 (88.23) | |
| ***Ethnicity,*** *n (%)* | - | - | - | 0.320# |
| Hispanic | 1 (4) | 1 (12.5) | - | - |
| Non-Hispanic | 24 (96) | 8 (87.5) | 17 (100) | - |
| **Stage at initial diagnosis,** *n (%)* | - | - | - | 0.0503# |
| I | 17 (68) | 3 (37.5) | 14 (82.35) | - |
| II | 6 (24) | 4 (50) | 2 (11.76) | - |
| III | 2 (8) | 1 (12.5) | 1 (5.88) | - |
| IV | - | - | - | - |
| ***Hormone receptor status of sample tested,*** *n (%)* | - | - | - | >0.999# |
| Positive | 25 (100) | 8 (100) | 17 (100) | - |
| Negative | - | - | - | - |
| ***Estrogen receptor status of the sample tested,*** *n (%)* | | | | >0.999# |
| Positive | 25 (100) | 8 (100) | 17 (100) | - |
| Negative | - | - | - | - |
| ***Histology at initial diagnosis,*** *n (%)* | - | - | - | 0.030# |

| | | | | |
|---|---|---|---|---|
| Ductal carcinoma in situ (DCIC) | - | - | - | - |
| Invasive ductal carcinoma (IDC) | 18 (72) | 4 (50) | 14 (82.35) | - |
| Invasive lobular carcinoma (ILC) | 5 (20) | 4 (50) | 1 (5.88) | - |
| Mixed (IDC & ILC) | 2 (8) | - | 2 (11.76) | - |
| ***Type of specimen tested,*** *n (%)* | - | - | - | > 0.999# |
| Primary breast | 25 (100) | 8 (100) | 17 (100) | |
| Local recurrence | - | - | - | - |
| Metastasis | - | - | - | - |
| ***Previous therapy,*** *n (%)* | | | | > 0.999# |
| Yes | - | - | - | - |
| no | 25 (100) | 8 (100) | 17 (100) | - |
| **Menopausal status at Initial diagnosis,** *n (%)* | - | - | - | 0.777# |
| Premenopausal | 14 (56) | 4 (50) | 10 (58.82) | - |
| Postmenopausal | 10 (40) | 4 (50) | 6 (35.29) | - |
| Male | 1 (4) | - | 1 (5.88) | - |
| ***Histologic grade,*** *n (%)* | - | - | - | >0.999# |
| I Low grade | 4 (16) | 1 (12.5) | 3 (17.64) | - |
| II Intermediate Grade | 12 (48) | 4 (50) | 8 (47.05) | - |
| III High Grade | 9 (36) | 3 (37.5) | 6 (35.29) | - |

Differences between HER2-0 and HER2-low for categorical variable were investigated using #Fisher's exact test, whereas continuous or nominal variables were assessed using a *t-test. All examinations were conducted utilizing two-sided tests. HER2 - human epidermal growth factor receptor 2.

**Supplementary table 1. Genes differentially expressed in HER2-0 and HER2-low bulk RNA-seq**

| Up regulated in HER2-0 | | |
|---|---|---|
| **Gene** | **logFC** | **Minus log (p value)** |
| PIP5K1B | -1.8044503 | 3.42598512 |
| TRIL | -1.1346097 | 3.33367896 |
| RPAP3-DT | -1.1853875 | 2.86039802 |
| MOXD1 | -1.337598 | 2.79516565 |
| GIMAP5 | -1.373463 | 2.73409649 |
| GRM7 | -1.6937179 | 2.70049023 |
| NRROS | -1.0141049 | 2.65456041 |
| SERPINF2 | -1.5021749 | 2.63990616 |
| ITIH3 | -1.1421137 | 2.62481285 |
| NIBAN1 | -1.1705746 | 2.60330773 |
| ZDBF2 | -1.2348476 | 2.59931678 |
| SULT1B1 | -1.419103 | 2.56395902 |
| ADGRE4P | -1.2875329 | 2.38877654 |
| PDCD1 | -1.5466184 | 2.30507508 |
| XCR1 | -1.187396 | 2.29864898 |
| PTGDS | -1.8654418 | 2.27618767 |
| ZNF582-DT | -1.1032218 | 2.242231 |
| FAM30A | -1.751164 | 2.19889784 |
| FRAS1 | -1.5135289 | 2.18726936 |
| CCDC9B | -1.1268279 | 2.17057532 |
| CACNA1H | -1.9508015 | 2.10756448 |
| WNT10A | -1.3257575 | 2.09863238 |
| IFNG-AS1 | -1.6718008 | 2.07626289 |
| GJB1 | -1.8796113 | 2.0397533 |
| IGKV1-9 | -2.0775106 | 2.03461292 |
| LOC100422687 | -1.0652536 | 2.02684365 |
| MAN1C1 | -1.0036934 | 2.02119488 |
| PLA2G2D | -2.0906634 | 2.01701289 |
| RASGRP2 | -1.0525025 | 2.01504416 |
| TDRD10 | -1.0725196 | 2.0126596 |
| ATP10B | -1.2572418 | 1.99864432 |
| LOC107985216 | -1.0311847 | 1.97534309 |
| LOC105374811 | -1.1034554 | 1.97411897 |
| PPARGC1A | -1.145964 | 1.95113911 |

| | | |
|---|---|---|
| CD40LG | -1.2638413 | 1.93762191 |
| VANGL2 | -1.2501014 | 1.91135508 |
| LINGO4 | -1.2124127 | 1.90916004 |
| NFKBIZ | -1.5345281 | 1.90881265 |
| FCRL1 | -1.603283 | 1.89900687 |
| SDK1 | -1.1465675 | 1.89728414 |
| PRKCQ | -1.1106359 | 1.88030907 |
| IGKV2-28 | -1.9655674 | 1.8775667 |
| TUBA4A | -1.0004639 | 1.86193852 |
| DLGAP2 | -1.3445217 | 1.82141877 |
| IGKJ4 | -2.203455 | 1.82003963 |
| ESPN | -1.140545 | 1.81544463 |
| IGKV1-5 | -2.3235403 | 1.80861162 |
| IGKV1-17 | -2.9350031 | 1.80740279 |
| CD163 | -1.1133222 | 1.80075672 |
| RNU1-155P | -1.7694621 | 1.79744715 |
| CPZ | -1.0071067 | 1.79492349 |
| LCNL1 | -1.3613406 | 1.792244 |
| ATP6V0D2 | -1.388265 | 1.78799345 |
| IGHJ5 | -1.9014976 | 1.78672615 |
| KCNA3 | -1.1706086 | 1.78537024 |
| ASPG | -1.0624788 | 1.78432224 |
| DEPP1 | -1.0202566 | 1.78138681 |
| P2RX5 | -1.461516 | 1.77955247 |
| IGLV7-46 | -2.2472877 | 1.77608568 |
| IGKJ1 | -1.9284655 | 1.76979548 |
| AIM2 | -1.3096241 | 1.76875765 |
| LYNX1 | -1.0686965 | 1.76398951 |
| IGKV1-27 | -2.6000196 | 1.74806507 |
| TNFSF14 | -1.3126251 | 1.74705174 |
| CD52 | -1.3287541 | 1.7422979 |
| ALDH1A1 | -1.0710344 | 1.74153921 |
| NUGGC | -1.2665686 | 1.74041429 |
| P2RY8 | -1.0372124 | 1.70651379 |
| CDHR1 | -1.5742437 | 1.69943799 |
| BNIPL | -1.3313362 | 1.68476936 |
| IGLV2-14 | -2.3413341 | 1.68224228 |
| IGKC | -2.012203 | 1.67150612 |
| LSAMP | -1.2796398 | 1.67013846 |

| | | |
|---|---|---|
| PVALB | -2.9156689 | 1.65958414 |
| LOC105370792 | -1.2472268 | 1.65735802 |
| CD79A | -1.5914227 | 1.65696393 |
| RORB | -1.765301 | 1.64251925 |
| CLEC4E | -1.3936556 | 1.6368457 |
| IGHJ4 | -1.8359857 | 1.63436543 |
| KCND3 | -1.4025061 | 1.62650244 |
| LOC124904627 | -1.313907 | 1.62556756 |
| IGKV3-20 | -2.2038229 | 1.62466411 |
| ALDH3B2 | -1.5528347 | 1.61988879 |
| FOLR2 | -1.2582621 | 1.6169228 |
| MYCN | -1.6840482 | 1.59959605 |
| IGLV2-8 | -1.9440267 | 1.59393073 |
| STUM | -1.849561 | 1.58925982 |
| PDK4 | -1.1058087 | 1.58849238 |
| IGLC3 | -2.0503814 | 1.58559334 |
| FAM177B | -1.008111 | 1.57916654 |
| HECW1 | -1.0180116 | 1.56951448 |
| RIMBP2 | -1.7166902 | 1.55721423 |
| LRRN4CL | -1.1464121 | 1.54552179 |
| CBLN4 | -1.0635444 | 1.52610064 |
| VSNL1 | -1.1225469 | 1.52394094 |
| LY9 | -1.3694241 | 1.52337224 |
| IGHJ3 | -1.6184591 | 1.51349329 |
| IGHM | -1.8562161 | 1.49537435 |
| TPSG1 | -2.230232 | 1.4948524 |
| MSX2 | -1.1501522 | 1.48758391 |
| SIGLEC8 | -1.1903361 | 1.47446166 |
| POU2AF1 | -1.592297 | 1.46269371 |
| CCR2 | -1.2638187 | 1.45453485 |
| IGSF21 | -1.17336 | 1.45213727 |
| IGHV4-39 | -2.1421729 | 1.44761559 |
| CPEB1 | -1.1470145 | 1.44757749 |
| ASS1P1 | -1.1272281 | 1.44414794 |
| IGKV1-39 | -2.001361 | 1.44161767 |
| IGKV1D-16 | -2.0631122 | 1.43801443 |
| ASCL2 | -1.3214721 | 1.43248225 |
| LZTS3 | -1.0779704 | 1.43235409 |
| BMP4 | -1.657917 | 1.4309053 |

| | | |
|---|---|---|
| SDCBP2 | -1.3849934 | 1.42737998 |
| APELA | -1.1889543 | 1.42635769 |
| NKX2-2 | -2.2827343 | 1.42599419 |
| SCNN1B | -1.2528265 | 1.42532988 |
| EDARADD | -1.5435933 | 1.41975067 |
| ANKRD30A | -1.690985 | 1.41224084 |
| IGLV1-40 | -2.1938967 | 1.40823716 |
| GABRD | -1.131667 | 1.40663099 |
| FRMPD3 | -1.0871878 | 1.39571782 |
| MS4A1 | -1.4144852 | 1.39399854 |
| IGHV1-2 | -2.2528693 | 1.39251262 |
| PRF1 | -1.3379289 | 1.38896462 |
| PTGFR | -1.3173188 | 1.38796988 |
| IGKV3-15 | -1.8042929 | 1.38370711 |
| DRAXIN | -1.0931511 | 1.38258819 |
| OSR1 | -1.5027898 | 1.37231333 |
| CD247 | -1.0073492 | 1.36657599 |
| IGKV3-11 | -2.0996085 | 1.36397564 |
| NR2F1 | -1.3153823 | 1.35336371 |
| IGLV2-11 | -1.5296941 | 1.35065157 |
| TMEM150C | -1.5661985 | 1.35043597 |
| FCRL5 | -1.4484627 | 1.34814502 |
| IGHV6-1 | -2.4209793 | 1.34329377 |
| CD38 | -1.1773044 | 1.34160265 |
| LINC01819 | -1.5620124 | 1.34051892 |
| TRGC2 | -1.1368866 | 1.33646281 |
| RPL21P95 | -1.0721863 | 1.33317319 |
| IGKV1D-39 | -1.7521969 | 1.33216479 |
| EBI3 | -1.1686258 | 1.33197322 |
| ATP6V1B1 | -1.1950937 | 1.32669297 |
| IGHV5-51 | -1.9638928 | 1.32635912 |
| PAX5 | -1.3762227 | 1.32488435 |
| RIPOR2 | -1.0959438 | 1.32324108 |
| TMEM119 | -1.1221826 | 1.31879947 |
| RASGRF1 | -1.1048661 | 1.31738133 |
| CLEC12A | -1.2309236 | 1.3172462 |
| MIR5087 | -1.5341054 | 1.31687714 |
| SCML4 | -1.3314341 | 1.31074226 |
| LINC00869;LINC02591 | -1.5948104 | 1.31036342 |

| | | |
|---|---|---|
| LRRC4B | -1.7097208 | 1.30819254 |
| FCRL6 | -1.0646064 | 1.30345885 |
| MTUS2 | -1.1174692 | 1.3000607 |

| Up regulated in HER2-low | | |
|---|---|---|
| **Gene** | **logFC** | **Minus log (p value)** |
| NDUFAF8 | 1.20538305 | 3.23202977 |
| MTND2P26 | 1.41796942 | 2.79890385 |
| ZNG1F | 1.00364325 | 2.69957572 |
| DSCAM-AS1 | 2.97884256 | 2.52665519 |
| ARMC3 | 2.10289961 | 2.44047526 |
| SEMA3B | 1.44359202 | 2.31546041 |
| CFAP91 | 1.18994672 | 2.24116814 |
| LEKR1 | 1.00008465 | 2.21908291 |
| RGPD2 | 1.4087033 | 2.18917053 |
| DPY19L2P1 | 1.371285 | 2.14685375 |
| DNAAF1 | 1.37926329 | 2.14106371 |
| SPAG16 | 1.31143648 | 2.10131044 |
| MTATP6P15 | 1.69784962 | 2.09731546 |
| MTND1P36 | 1.46981547 | 2.07263309 |
| OR52E5 | 1.52766581 | 2.06526957 |
| EPPK1 | 1.00730041 | 2.0434758 |
| RPL13P6 | 1.04846778 | 2.01029988 |
| RGS22 | 1.88718408 | 1.96939443 |
| BMP8A | 1.14687737 | 1.91344452 |
| MTCO2P15 | 1.22533692 | 1.8158831 |
| APLN | 1.23103637 | 1.78146555 |
| TREM1 | 1.11540557 | 1.77108398 |
| MIR181A2HG | 1.04377883 | 1.76911433 |
| LINC01992 | 1.84459526 | 1.75949117 |
| ADGB | 1.4532455 | 1.7285175 |
| KCNJ3 | 2.38531011 | 1.65913276 |
| RNF17 | 1.54914699 | 1.64520236 |
| OR52E7P | 1.28594597 | 1.63870031 |
| AREG | 1.82642551 | 1.63781551 |
| CFAP43 | 1.01231715 | 1.61018519 |
| OR52E6 | 1.52277581 | 1.59935424 |

| | | |
|---|---|---|
| LOC100506071 | 1.0029632 | 1.54542165 |
| MIR4477B | 1.57489188 | 1.53900028 |
| LINC02257 | 1.30035797 | 1.5379332 |
| MIR548T | 1.01259664 | 1.53199678 |
| ROPN1L | 1.12089166 | 1.52056839 |
| MEGF10 | 1.20691271 | 1.48130091 |
| SYT13 | 2.13463349 | 1.46058715 |
| LNCOC1 | 1.39199616 | 1.4471099 |
| B4GALNT3 | 1.23202658 | 1.43769476 |
| COX6C | 1.45998408 | 1.40084622 |
| LOC101928651 | 1.35692021 | 1.4001241 |
| LINC02303 | 1.24689316 | 1.38549032 |
| SLC1A2 | 1.27681648 | 1.373854 |
| GPR83 | 1.24462231 | 1.37271217 |
| KDM5D | 1.39848059 | 1.36850391 |
| AARD | 1.39329492 | 1.33470772 |
| SGCG | 1.14533814 | 1.30767105 |
| SPP1 | 1.16374599 | 1.30251428 |

$p<0.05$ = (minus log10) pvalue > 1.301 and fold change (FC) > 1

### Supplementary table 3. CNV of genes from chromosome 17 in HER2-0 and HER2-low epithelial cells.

| CELL_GROUP | GENE_REGION_NAME | CNV STATE | CNV EVENT | GENE | CHR | START | END |
|---|---|---|---|---|---|---|---|
| **HER2-0.HER2-0_S1** | 17-region_1 | 1 | Loss | RPH3AL | 17 | 212389 | 386254 |
| **HER2-0.HER2-0_S1** | 17-region_1 | 1 | Loss | VPS53 | 17 | 508668 | 721717 |
| **HER2-0.HER2-0_S1** | 17-region_1 | 1 | Loss | GLOD4 | 17 | 757097 | 783390 |
| **HER2-0.HER2-0_S1** | 17-region_1 | 1 | Loss | ABR | 17 | 1003518 | 1229021 |
| **HER2-0.HER2-0_S1** | 17-region_1 | 1 | Loss | YWHAE | 17 | 1344272 | 1400378 |
| **HER2-0.HER2-0_S1** | 17-region_1 | 1 | Loss | CRK | 17 | 1420689 | 1463162 |
| **HER2-0.HER2-0_S1** | 17-region_1 | 1 | Loss | MYO1C | 17 | 1464098 | 1492812 |
| **HER2-0.HER2-0_S1** | 17-region_1 | 1 | Loss | PITPNA | 17 | 1517718 | 1562816 |
| **HER2-0.HER2-0_S1** | 17-region_1 | 1 | Loss | SLC43A2 | 17 | 1569267 | 1628886 |
| **HER2-0.HER2-0_S1** | 17-region_1 | 1 | Loss | PRPF8 | 17 | 1650629 | 1684882 |
| **HER2-0.HER2-0_S1** | 17-region_1 | 1 | Loss | SMYD4 | 17 | 1779485 | 1830634 |
| **HER2-0.HER2-0_S1** | 17-region_1 | 1 | Loss | RPA1 | 17 | 1829702 | 1900082 |
| **HER2-0.HER2-0_S1** | 17-region_1 | 1 | Loss | RTN4RL1 | 17 | 1934677 | 2025345 |
| **HER2-0.HER2-0_S1** | 17-region_1 | 1 | Loss | DPH1 | 17 | 2030110 | 2043430 |
| **HER2-0.HER2-0_S1** | 17-region_1 | 1 | Loss | SMG6 | 17 | 2059839 | 2303771 |
| **HER2-0.HER2-0_S1** | 17-region_1 | 1 | Loss | TSR1 | 17 | 2322503 | 2337507 |
| **HER2-0.HER2-0_S1** | 17-region_1 | 1 | Loss | SGSM2 | 17 | 2337498 | 2381058 |
| **HER2-0.HER2-0_S1** | 17-region_1 | 1 | Loss | MNT | 17 | 2384060 | 2401118 |
| **HER2-0.HER2-0_S1** | 17-region_1 | 1 | Loss | METTL16 | 17 | 2405562 | 2511891 |
| **HER2-0.HER2-0_S1** | 17-region_1 | 1 | Loss | PAFAH1B1 | 17 | 2593210 | 2685615 |
| **HER2-0.HER2-0_S1** | 17-region_1 | 1 | Loss | CLUH | 17 | 2689386 | 2712663 |
| **HER2-0.HER2-0_S1** | 17-region_1 | 1 | Loss | RAP1GAP2 | 17 | 2755705 | 3037739 |
| **HER2-0.HER2-0_S1** | 17-region_1 | 1 | Loss | SHPK | 17 | 3608262 | 3636322 |
| **HER2-0.HER2-0_S1** | 17-region_1 | 1 | Loss | ITGAE | 17 | 3714628 | 3801243 |
| **HER2-0.HER2-0_S1** | 17-region_1 | 1 | Loss | NCBP3 | 17 | 3802165 | 3846251 |
| **HER2-0.HER2-0_S1** | 17-region_1 | 1 | Loss | ATP2A3 | 17 | 3923870 | 3964464 |
| **HER2-0.HER2-0_S1** | 17-region_1 | 1 | Loss | ZZEF1 | 17 | 4004445 | 4143020 |
| **HER2-0.HER2-0_S1** | 17-region_1 | 1 | Loss | ANKFY1 | 17 | 4163907 | 4263977 |
| **HER2-0.HER2-0_S1** | 17-region_1 | 1 | Loss | UBE2G1 | 17 | 4269259 | 4366628 |
| **HER2-0.HER2-0_S1** | 17-region_1 | 1 | Loss | MYBBP1A | 17 | 4538897 | 4555631 |
| **HER2-0.HER2-0_S1** | 17-region_1 | 1 | Loss | PELP1 | 17 | 4669774 | 4704337 |
| **HER2-0.HER2-0_S1** | 17-region_1 | 1 | Loss | PSMB6 | 17 | 4796144 | 4798503 |
| **HER2-0.HER2-0_S1** | 17-region_1 | 1 | Loss | PLD2 | 17 | 4807096 | 4823434 |
| **HER2-0.HER2-0_S1** | 17-region_1 | 1 | Loss | MINK1 | 17 | 4833388 | 4898061 |
| **HER2-0.HER2-0_S1** | 17-region_1 | 1 | Loss | RNF167 | 17 | 4940008 | 4945222 |
| **HER2-0.HER2-0_S1** | 17-region_1 | 1 | Loss | PFN1 | 17 | 4945652 | 4949061 |

| | | | | | | | |
|---|---|---|---|---|---|---|---|
| **HER2-0.HER2-0_S1** | 17-region_1 | 1 | Loss | CAMTA2 | 17 | 4967992 | 4987652 |
| **HER2-0.HER2-0_S1** | 17-region_1 | 1 | Loss | KIF1C | 17 | 4997948 | 5028401 |
| **HER2-0.HER2-0_S1** | 17-region_1 | 1 | Loss | ZNF594 | 17 | 5179536 | 5191883 |
| **HER2-0.HER2-0_S1** | 17-region_1 | 1 | Loss | RABEP1 | 17 | 5282265 | 5385812 |
| **HER2-0.HER2-0_S1** | 17-region_1 | 1 | Loss | NUP88 | 17 | 5360963 | 5420160 |
| **HER2-0.HER2-0_S1** | 17-region_1 | 1 | Loss | RPAIN | 17 | 5419641 | 5432876 |
| **HER2-0.HER2-0_S1** | 17-region_1 | 1 | Loss | C1QBP | 17 | 5432777 | 5448830 |
| **HER2-0.HER2-0_S1** | 17-region_1 | 1 | Loss | DERL2 | 17 | 5471251 | 5486811 |
| **HER2-0.HER2-0_S1** | 17-region_1 | 1 | Loss | KIAA0753 | 17 | 6578148 | 6640927 |
| **HER2-0.HER2-0_S1** | 17-region_1 | 1 | Loss | XAF1 | 17 | 6755447 | 6775647 |
| **HER2-0.HER2-0_S1** | 17-region_1 | 1 | Loss | ALOX12-AS1 | 17 | 6876635 | 7012349 |
| **HER2-0.HER2-0_S1** | 17-region_1 | 1 | Loss | RNASEK | 17 | 7012417 | 7014532 |
| **HER2-0.HER2-0_S1** | 17-region_1 | 1 | Loss | ACADVL | 17 | 7217125 | 7225273 |
| **HER2-0.HER2-0_S1** | 17-region_1 | 1 | Loss | GABARAP | 17 | 7240014 | 7242770 |
| **HER2-0.HER2-0_S1** | 17-region_1 | 1 | Loss | CTDNEP1 | 17 | 7243591 | 7252491 |
| **HER2-0.HER2-0_S1** | 17-region_1 | 1 | Loss | CLDN7 | 17 | 7259903 | 7263983 |
| **HER2-0.HER2-0_S1** | 17-region_1 | 1 | Loss | EIF5A | 17 | 7306999 | 7312463 |
| **HER2-0.HER2-0_S1** | 17-region_1 | 1 | Loss | GPS2 | 17 | 7311324 | 7315564 |
| **HER2-0.HER2-0_S1** | 17-region_1 | 1 | Loss | ZBTB4 | 17 | 7459366 | 7484263 |
| **HER2-0.HER2-0_S1** | 17-region_1 | 1 | Loss | POLR2A | 17 | 7484366 | 7514616 |
| **HER2-0.HER2-0_S1** | 17-region_1 | 1 | Loss | TNFSF13 | 17 | 7558292 | 7561608 |
| **HER2-0.HER2-0_S1** | 17-region_3 | 6 | Gain | CDRT4 | 17 | 15436015 | 15503608 |
| **HER2-0.HER2-0_S1** | 17-region_3 | 6 | Gain | TVP23C | 17 | 15502264 | 15563595 |
| **HER2-0.HER2-0_S1** | 17-region_3 | 6 | Gain | TRIM16 | 17 | 15627960 | 15684311 |
| **HER2-0.HER2-0_S1** | 17-region_3 | 6 | Gain | ZSWIM7 | 17 | 15976560 | 15999717 |
| **HER2-0.HER2-0_S1** | 17-region_3 | 6 | Gain | TTC19 | 17 | 15999380 | 16045015 |
| **HER2-0.HER2-0_S1** | 17-region_3 | 6 | Gain | NCOR1 | 17 | 16029157 | 16218185 |
| **HER2-0.HER2-0_S1** | 17-region_3 | 6 | Gain | PIGL | 17 | 16217191 | 16351797 |
| **HER2-0.HER2-0_S1** | 17-region_3 | 6 | Gain | UBB | 17 | 16380798 | 16382745 |
| **HER2-0.HER2-0_S1** | 17-region_3 | 6 | Gain | CCDC144A | 17 | 16689537 | 16777881 |
| **HER2-0.HER2-0_S1** | 17-region_3 | 6 | Gain | MPRIP | 17 | 17042545 | 17217679 |
| **HER2-0.HER2-0_S1** | 17-region_3 | 6 | Gain | FLCN | 17 | 17212212 | 17237188 |
| **HER2-0.HER2-0_S1** | 17-region_3 | 6 | Gain | COPS3 | 17 | 17246820 | 17281293 |
| **HER2-0.HER2-0_S1** | 17-region_3 | 6 | Gain | RAI1 | 17 | 17681473 | 17811453 |
| **HER2-0.HER2-0_S1** | 17-region_3 | 6 | Gain | SREBF1 | 17 | 17810399 | 17837002 |
| **HER2-0.HER2-0_S1** | 17-region_4 | 6 | Gain | TOM1L2 | 17 | 17843511 | 17972422 |
| **HER2-0.HER2-0_S1** | 17-region_4 | 6 | Gain | DRC3 | 17 | 17972813 | 18016889 |
| **HER2-0.HER2-0_S1** | 17-region_4 | 6 | Gain | ALKBH5 | 17 | 18183078 | 18209954 |
| **HER2-0.HER2-0_S1** | 17-region_4 | 6 | Gain | FLII | 17 | 18244836 | 18258916 |
| **HER2-0.HER2-0_S1** | 17-region_4 | 6 | Gain | TOP3A | 17 | 18271428 | 18315007 |

| | | | | | | | |
|---|---|---|---|---|---|---|---|
| **HER2-0.HER2-0_S1** | 17-region_4 | 6 | Gain | SHMT1 | 17 | 18327860 | 18363563 |
| **HER2-0.HER2-0_S1** | 17-region_4 | 6 | Gain | TVP23B | 17 | 18780995 | 18806714 |
| **HER2-0.HER2-0_S1** | 17-region_4 | 6 | Gain | PRPSAP2 | 17 | 18840085 | 18931287 |
| **HER2-0.HER2-0_S1** | 17-region_4 | 6 | Gain | EPN2 | 17 | 19215615 | 19336715 |
| **HER2-0.HER2-0_S1** | 17-region_4 | 6 | Gain | B9D1 | 17 | 19337554 | 19378182 |
| **HER2-0.HER2-0_S1** | 17-region_4 | 6 | Gain | ALDH3A2 | 17 | 19648136 | 19677598 |
| **HER2-0.HER2-0_S1** | 17-region_4 | 6 | Gain | ULK2 | 17 | 19770829 | 19867936 |
| **HER2-0.HER2-0_S1** | 17-region_4 | 6 | Gain | AKAP10 | 17 | 19904302 | 19978343 |
| **HER2-0.HER2-0_S1** | 17-region_4 | 6 | Gain | SPECC1 | 17 | 20009344 | 20319026 |
| **HER2-0.HER2-0_S1** | 17-region_4 | 6 | Gain | USP22 | 17 | 20999593 | 21043760 |
| **HER2-0.HER2-0_S1** | 17-region_4 | 6 | Gain | DHRS7B | 17 | 21123364 | 21193265 |
| **HER2-0.HER2-0_S1** | 17-region_4 | 6 | Gain | MAP2K3 | 17 | 21284672 | 21315240 |
| **HER2-0.HER2-0_S1** | 17-region_4 | 6 | Gain | WSB1 | 17 | 27294076 | 27315926 |
| **HER2-0.HER2-0_S1** | 17-region_4 | 6 | Gain | KSR1 | 17 | 27456714 | 27626438 |
| **HER2-0.HER2-0_S1** | 17-region_4 | 6 | Gain | NLK | 17 | 28041737 | 28196381 |
| **HER2-0.HER2-0_S1** | 17-region_4 | 6 | Gain | IFT20 | 17 | 28328325 | 28335489 |
| **HER2-0.HER2-0_S1** | 17-region_4 | 6 | Gain | TNFAIP1 | 17 | 28335602 | 28347009 |
| **HER2-0.HER2-0_S1** | 17-region_4 | 6 | Gain | POLDIP2 | 17 | 28347177 | 28357522 |
| **HER2-0.HER2-0_S1** | 17-region_4 | 6 | Gain | TMEM199 | 17 | 28357581 | 28363683 |
| **HER2-0.HER2-0_S1** | 17-region_4 | 6 | Gain | SLC46A1 | 17 | 28394756 | 28407197 |
| **HER2-0.HER2-0_S1** | 17-region_4 | 6 | Gain | UNC119 | 17 | 28546707 | 28552668 |
| **HER2-0.HER2-0_S1** | 17-region_4 | 6 | Gain | PIGS | 17 | 28553383 | 28571872 |
| **HER2-0.HER2-0_S1** | 17-region_4 | 6 | Gain | KIAA0100 | 17 | 28614440 | 28645454 |
| **HER2-0.HER2-0_S1** | 17-region_4 | 6 | Gain | SDF2 | 17 | 28648356 | 28662189 |
| **HER2-0.HER2-0_S1** | 17-region_4 | 6 | Gain | SUPT6H | 17 | 28662091 | 28702684 |
| **HER2-0.HER2-0_S1** | 17-region_4 | 6 | Gain | RPL23A | 17 | 28719393 | 28724359 |
| **HER2-0.HER2-0_S1** | 17-region_4 | 6 | Gain | TRAF4 | 17 | 28743984 | 28750958 |
| **HER2-0.HER2-0_S1** | 17-region_4 | 6 | Gain | FAM222B | 17 | 28755978 | 28855232 |
| **HER2-0.HER2-0_S1** | 17-region_4 | 6 | Gain | ERAL1 | 17 | 28854938 | 28861067 |
| **HER2-0.HER2-0_S1** | 17-region_4 | 6 | Gain | FLOT2 | 17 | 28879335 | 28897679 |
| **HER2-0.HER2-0_S1** | 17-region_4 | 6 | Gain | PHF12 | 17 | 28905250 | 28951771 |
| **HER2-0.HER2-0_S1** | 17-region_4 | 6 | Gain | MYO18A | 17 | 29071124 | 29180412 |
| **HER2-0.HER2-0_S1** | 17-region_4 | 6 | Gain | NUFIP2 | 17 | 29255836 | 29294118 |
| **HER2-0.HER2-0_S1** | 17-region_4 | 6 | Gain | TAOK1 | 17 | 29390464 | 29551904 |
| **HER2-0.HER2-0_S1** | 17-region_4 | 6 | Gain | TP53I13 | 17 | 29566052 | 29573157 |
| **HER2-0.HER2-0_S1** | 17-region_4 | 6 | Gain | GIT1 | 17 | 29573469 | 29594054 |
| **HER2-0.HER2-0_S1** | 17-region_4 | 6 | Gain | CORO6 | 17 | 29614756 | 29622907 |
| **HER2-0.HER2-0_S1** | 17-region_4 | 6 | Gain | SSH2 | 17 | 29625938 | 29930276 |
| **HER2-0.HER2-0_S1** | 17-region_4 | 6 | Gain | NSRP1 | 17 | 30115521 | 30186475 |
| **HER2-0.HER2-0_S1** | 17-region_4 | 6 | Gain | BLMH | 17 | 30248195 | 30292056 |

| | | | | | | | |
|---|---|---|---|---|---|---|---|
| **HER2-0.HER2-0_S1** | 17-region_4 | 6 | Gain | CPD | 17 | 30378905 | 30469989 |
| **HER2-0.HER2-0_S1** | 17-region_4 | 6 | Gain | GOSR1 | 17 | 30477362 | 30527592 |
| **HER2-0.HER2-0_S1** | 17-region_4 | 6 | Gain | AC005562.1 | 17 | 30576464 | 30672789 |
| **HER2-0.HER2-0_S1** | 17-region_4 | 6 | Gain | CRLF3 | 17 | 30769388 | 30824776 |
| **HER2-0.HER2-0_S1** | 17-region_4 | 6 | Gain | ATAD5 | 17 | 30831970 | 30895869 |
| **HER2-0.HER2-0_S1** | 17-region_4 | 6 | Gain | TEFM | 17 | 30897336 | 30906820 |
| **HER2-0.HER2-0_S1** | 17-region_4 | 6 | Gain | RNF135 | 17 | 30968785 | 30999911 |
| **HER2-0.HER2-0_S1** | 17-region_4 | 6 | Gain | NF1 | 17 | 31094927 | 31382116 |
| **HER2-0.HER2-0_S1** | 17-region_4 | 6 | Gain | RAB11FIP4 | 17 | 31391624 | 31538217 |
| **HER2-0.HER2-0_S1** | 17-region_5 | 1 | Loss | UTP6 | 17 | 31860899 | 31901765 |
| **HER2-0.HER2-0_S1** | 17-region_5 | 1 | Loss | SUZ12 | 17 | 31937018 | 32001045 |
| **HER2-0.HER2-0_S1** | 17-region_5 | 1 | Loss | RHOT1 | 17 | 32142454 | 32253374 |
| **HER2-0.HER2-0_S1** | 17-region_5 | 1 | Loss | C17orf75 | 17 | 32324565 | 32350023 |
| **HER2-0.HER2-0_S1** | 17-region_5 | 1 | Loss | ZNF207 | 17 | 32350109 | 32381886 |
| **HER2-0.HER2-0_S1** | 17-region_5 | 1 | Loss | PSMD11 | 17 | 32444261 | 32483318 |
| **HER2-0.HER2-0_S1** | 17-region_5 | 1 | Loss | MYO1D | 17 | 32492522 | 32877177 |
| **HER2-0.HER2-0_S1** | 17-region_5 | 1 | Loss | ASIC2 | 17 | 33013087 | 34174964 |
| **HER2-0.HER2-0_S1** | 17-region_5 | 1 | Loss | ZNF830 | 17 | 34961530 | 34963775 |
| **HER2-0.HER2-0_S1** | 17-region_5 | 1 | Loss | LIG3 | 17 | 34980494 | 35009743 |
| **HER2-0.HER2-0_S1** | 17-region_5 | 1 | Loss | RFFL | 17 | 35005990 | 35089319 |
| **HER2-0.HER2-0_S1** | 17-region_5 | 1 | Loss | SLFN5 | 17 | 35243036 | 35273655 |
| **HER2-0.HER2-0_S1** | 17-region_5 | 1 | Loss | AP2B1 | 17 | 35578046 | 35726409 |
| **HER2-0.HER2-0_S1** | 17-region_5 | 1 | Loss | TAF15 | 17 | 35713791 | 35864615 |
| **HER2-0.HER2-0_S1** | 17-region_5 | 1 | Loss | MYO19 | 17 | 36495633 | 36543435 |
| **HER2-0.HER2-0_S1** | 17-region_5 | 1 | Loss | GGNBP2 | 17 | 36544888 | 36589848 |
| **HER2-0.HER2-0_S1** | 17-region_5 | 1 | Loss | AATF | 17 | 36948875 | 37056871 |
| **HER2-0.HER2-0_S1** | 17-region_5 | 1 | Loss | ACACA | 17 | 37084988 | 37406818 |
| **HER2-0.HER2-0_S1** | 17-region_5 | 1 | Loss | TADA2A | 17 | 37406874 | 37479730 |
| **HER2-0.HER2-0_S1** | 17-region_5 | 1 | Loss | SYNRG | 17 | 37514797 | 37609496 |
| **HER2-0.HER2-0_S1** | 17-region_5 | 1 | Loss | DDX52 | 17 | 37609739 | 37643464 |
| **HER2-0.HER2-0_S1** | 17-region_5 | 1 | Loss | MRPL45 | 17 | 38297023 | 38323218 |
| **HER2-0.HER2-0_S1** | 17-region_5 | 1 | Loss | SOCS7 | 17 | 38352228 | 38405593 |
| **HER2-0.HER2-0_S1** | 17-region_5 | 1 | Loss | SRCIN1 | 17 | 38530016 | 38605930 |
| **HER2-0.HER2-0_S1** | 17-region_5 | 1 | Loss | MLLT6 | 17 | 38705542 | 38729803 |
| **HER2-0.HER2-0_S1** | 17-region_7 | 1 | Loss | LSM12 | 17 | 44034635 | 44067619 |
| **HER2-0.HER2-0_S1** | 17-region_7 | 1 | Loss | G6PC3 | 17 | 44070730 | 44076344 |
| **HER2-0.HER2-0_S1** | 17-region_7 | 1 | Loss | HDAC5 | 17 | 44076746 | 44123702 |
| **HER2-0.HER2-0_S1** | 17-region_7 | 1 | Loss | ASB16-AS1 | 17 | 44175973 | 44186717 |
| **HER2-0.HER2-0_S1** | 17-region_7 | 1 | Loss | TMUB2 | 17 | 44186970 | 44191731 |
| **HER2-0.HER2-0_S1** | 17-region_7 | 1 | Loss | ATXN7L3 | 17 | 44191805 | 44200113 |

| | | | | | | | |
|---|---|---|---|---|---|---|---|
| **HER2-0.HER2-0_S1** | 17-region_7 | 1 | Loss | UBTF | 17 | 44205033 | 44221626 |
| **HER2-0.HER2-0_S1** | 17-region_7 | 1 | Loss | SLC25A39 | 17 | 44319625 | 44324870 |
| **HER2-0.HER2-0_S1** | 17-region_7 | 1 | Loss | GRN | 17 | 44345086 | 44353102 |
| **HER2-0.HER2-0_S1** | 17-region_7 | 1 | Loss | GPATCH8 | 17 | 44395284 | 44503430 |
| **HER2-0.HER2-0_S1** | 17-region_7 | 1 | Loss | EFTUD2 | 17 | 44849943 | 44899662 |
| **HER2-0.HER2-0_S1** | 17-region_7 | 1 | Loss | DCAKD | 17 | 45023340 | 45061109 |
| **HER2-0.HER2-0_S1** | 17-region_7 | 1 | Loss | NMT1 | 17 | 45051610 | 45109016 |
| **HER2-0.HER2-0_S1** | 17-region_7 | 1 | Loss | MAP3K14 | 17 | 45263121 | 45317040 |
| **HER2-0.HER2-0_S1** | 17-region_7 | 1 | Loss | ARHGAP27 | 17 | 45393902 | 45434421 |
| **HER2-0.HER2-0_S1** | 17-region_7 | 1 | Loss | PLEKHM1 | 17 | 45435900 | 45490749 |
| **HER2-0.HER2-0_S1** | 17-region_7 | 1 | Loss | MAPT-AS1 | 17 | 45799390 | 45895630 |
| **HER2-0.HER2-0_S1** | 17-region_7 | 1 | Loss | MAPT | 17 | 45894382 | 46028334 |
| **HER2-0.HER2-0_S1** | 17-region_7 | 1 | Loss | KANSL1 | 17 | 46029916 | 46225374 |
| **HER2-0.HER2-0_S1** | 17-region_7 | 1 | Loss | ARL17B | 17 | 46274784 | 46361797 |
| **HER2-0.HER2-0_S1** | 17-region_7 | 1 | Loss | NSF | 17 | 46590669 | 46757464 |
| **HER2-0.HER2-0_S1** | 17-region_7 | 1 | Loss | GOSR2 | 17 | 46923117 | 46967019 |
| **HER2-0.HER2-0_S1** | 17-region_7 | 1 | Loss | CDC27 | 17 | 47117703 | 47189422 |
| **HER2-0.HER2-0_S1** | 17-region_7 | 1 | Loss | EFCAB13 | 17 | 47323290 | 47441312 |
| **HER2-0.HER2-0_S1** | 17-region_7 | 1 | Loss | NPEPPS | 17 | 47522942 | 47623276 |
| **HER2-0.HER2-0_S1** | 17-region_7 | 1 | Loss | KPNB1 | 17 | 47649476 | 47685505 |
| **HER2-0.HER2-0_S1** | 17-region_7 | 1 | Loss | SCRN2 | 17 | 47837692 | 47841333 |
| **HER2-0.HER2-0_S1** | 17-region_7 | 1 | Loss | SP2 | 17 | 47896150 | 47928957 |
| **HER2-0.HER2-0_S1** | 17-region_7 | 1 | Loss | CDK5RAP3 | 17 | 47967810 | 47981774 |
| **HER2-0.HER2-0_S1** | 17-region_7 | 1 | Loss | NFE2L1 | 17 | 48048329 | 48061487 |
| **HER2-0.HER2-0_S1** | 17-region_7 | 1 | Loss | CBX1 | 17 | 48070052 | 48101521 |
| **HER2-0.HER2-0_S1** | 17-region_7 | 1 | Loss | SKAP1 | 17 | 48133440 | 48430275 |
| **HER2-0.HER2-0_S1** | 17-region_7 | 1 | Loss | HOXB2 | 17 | 48540894 | 48544989 |
| **HER2-0.HER2-0_S1** | 17-region_7 | 1 | Loss | HOXB3 | 17 | 48548870 | 48604912 |
| **HER2-0.HER2-0_S1** | 17-region_9 | 6 | Gain | COL1A1 | 17 | 50183289 | 50201632 |
| **HER2-0.HER2-0_S1** | 17-region_9 | 6 | Gain | XYLT2 | 17 | 50346092 | 50363138 |
| **HER2-0.HER2-0_S1** | 17-region_9 | 6 | Gain | MRPL27 | 17 | 50367857 | 50373214 |
| **HER2-0.HER2-0_S1** | 17-region_9 | 6 | Gain | LRRC59 | 17 | 50375059 | 50397553 |
| **HER2-0.HER2-0_S1** | 17-region_9 | 6 | Gain | ACSF2 | 17 | 50426158 | 50474845 |
| **HER2-0.HER2-0_S1** | 17-region_9 | 6 | Gain | RSAD1 | 17 | 50478800 | 50485975 |
| **HER2-0.HER2-0_S1** | 17-region_9 | 6 | Gain | SPATA20 | 17 | 50543058 | 50555852 |
| **HER2-0.HER2-0_S1** | 17-region_9 | 6 | Gain | ABCC3 | 17 | 50634777 | 50692252 |
| **HER2-0.HER2-0_S1** | 17-region_10 | 6 | Gain | ANKRD40 | 17 | 50693190 | 50707924 |
| **HER2-0.HER2-0_S1** | 17-region_10 | 6 | Gain | LUC7L3 | 17 | 50719544 | 50756213 |
| **HER2-0.HER2-0_S1** | 17-region_10 | 6 | Gain | TOB1 | 17 | 50862223 | 50867978 |
| **HER2-0.HER2-0_S1** | 17-region_10 | 6 | Gain | SPAG9 | 17 | 50962174 | 51120865 |

| | | | | | | | |
|---|---|---|---|---|---|---|---|
| **HER2-0.HER2-0_S1** | 17-region_10 | 6 | Gain | NME1 | 17 | 51153536 | 51162428 |
| **HER2-0.HER2-0_S1** | 17-region_10 | 6 | Gain | NME2 | 17 | 51165435 | 51171747 |
| **HER2-0.HER2-0_S1** | 17-region_10 | 6 | Gain | MBTD1 | 17 | 51177425 | 51260163 |
| **HER2-0.HER2-0_S1** | 17-region_10 | 6 | Gain | UTP18 | 17 | 51260528 | 51297936 |
| **HER2-0.HER2-0_S1** | 17-region_10 | 6 | Gain | TOM1L1 | 17 | 54899387 | 54961956 |
| **HER2-0.HER2-0_S1** | 17-region_10 | 6 | Gain | COX11 | 17 | 54951902 | 54968785 |
| **HER2-0.HER2-0_S1** | 17-region_10 | 6 | Gain | STXBP4 | 17 | 54968727 | 55173632 |
| **HER2-0.HER2-0_S1** | 17-region_10 | 6 | Gain | ANKFN1 | 17 | 55882301 | 56511659 |
| **HER2-0.HER2-0_S1** | 17-region_10 | 6 | Gain | C17orf67 | 17 | 56791913 | 56838773 |
| **HER2-0.HER2-0_S1** | 17-region_10 | 6 | Gain | DGKE | 17 | 56834099 | 56869567 |
| **HER2-0.HER2-0_S1** | 17-region_10 | 6 | Gain | TRIM25 | 17 | 56887909 | 56914038 |
| **HER2-0.HER2-0_S1** | 17-region_10 | 6 | Gain | COIL | 17 | 56938187 | 56961054 |
| **HER2-0.HER2-0_S1** | 17-region_10 | 6 | Gain | SCPEP1 | 17 | 56978105 | 57006768 |
| **HER2-0.HER2-0_S1** | 17-region_10 | 6 | Gain | AKAP1 | 17 | 57085092 | 57121349 |
| **HER2-0.HER2-0_S1** | 17-region_10 | 6 | Gain | MSI2 | 17 | 57255851 | 57684685 |
| **HER2-0.HER2-0_S1** | 17-region_10 | 6 | Gain | MRPS23 | 17 | 57834781 | 57850056 |
| **HER2-0.HER2-0_S1** | 17-region_10 | 6 | Gain | CUEDC1 | 17 | 57861243 | 57955323 |
| **HER2-0.HER2-0_S1** | 17-region_10 | 6 | Gain | VEZF1 | 17 | 57971547 | 57988259 |
| **HER2-0.HER2-0_S1** | 17-region_10 | 6 | Gain | SRSF1 | 17 | 58003360 | 58007346 |
| **HER2-0.HER2-0_S1** | 17-region_10 | 6 | Gain | DYNLL2 | 17 | 58083415 | 58095536 |
| **HER2-0.HER2-0_S1** | 17-region_10 | 6 | Gain | MKS1 | 17 | 58205436 | 58219605 |
| **HER2-0.HER2-0_S1** | 17-region_10 | 6 | Gain | TSPOAP1 | 17 | 58301228 | 58328760 |
| **HER2-0.HER2-0_S1** | 17-region_10 | 6 | Gain | TSPOAP1-AS1 | 17 | 58325450 | 58415766 |
| **HER2-0.HER2-0_S1** | 17-region_10 | 6 | Gain | SUPT4H1 | 17 | 58345175 | 58353093 |
| **HER2-0.HER2-0_S1** | 17-region_10 | 6 | Gain | RNF43 | 17 | 58352500 | 58417595 |
| **HER2-0.HER2-0_S1** | 17-region_10 | 6 | Gain | MTMR4 | 17 | 58489529 | 58517905 |
| **HER2-0.HER2-0_S1** | 17-region_10 | 6 | Gain | TEX14 | 17 | 58556678 | 58692055 |
| **HER2-0.HER2-0_S1** | 17-region_10 | 6 | Gain | RAD51C | 17 | 58692573 | 58735611 |
| **HER2-0.HER2-0_S1** | 17-region_10 | 6 | Gain | PPM1E | 17 | 58755869 | 58985176 |
| **HER2-0.HER2-0_S1** | 17-region_10 | 6 | Gain | TRIM37 | 17 | 58982638 | 59106921 |
| **HER2-0.HER2-0_S1** | 17-region_10 | 6 | Gain | SKA2 | 17 | 59109951 | 59155269 |
| **HER2-0.HER2-0_S1** | 17-region_10 | 6 | Gain | GDPD1 | 17 | 59220467 | 59275967 |
| **HER2-0.HER2-0_S1** | 17-region_10 | 6 | Gain | YPEL2 | 17 | 59331689 | 59401729 |
| **HER2-0.HER2-0_S1** | 17-region_10 | 6 | Gain | DHX40 | 17 | 59565525 | 59608345 |
| **HER2-0.HER2-0_S1** | 17-region_10 | 6 | Gain | CLTC | 17 | 59619689 | 59696956 |
| **HER2-0.HER2-0_S1** | 17-region_10 | 6 | Gain | PTRH2 | 17 | 59674636 | 59707626 |
| **HER2-0.HER2-0_S1** | 17-region_10 | 6 | Gain | VMP1 | 17 | 59707192 | 59842255 |
| **HER2-0.HER2-0_S1** | 17-region_10 | 6 | Gain | TUBD1 | 17 | 59859482 | 59892945 |
| **HER2-0.HER2-0_S1** | 17-region_10 | 6 | Gain | RPS6KB1 | 17 | 59893046 | 59950564 |
| **HER2-0.HER2-0_S1** | 17-region_10 | 6 | Gain | RNFT1 | 17 | 59952240 | 59964761 |

| | | | | | | | |
|---|---|---|---|---|---|---|---|
| **HER2-0.HER2-0_S1** | 17-region_10 | 6 | Gain | HEATR6 | 17 | 60043194 | 60078931 |
| **HER2-0.HER2-0_S1** | 17-region_10 | 6 | Gain | USP32 | 17 | 60179094 | 60422470 |
| **HER2-0.HER2-0_S1** | 17-region_10 | 6 | Gain | APPBP2 | 17 | 60443149 | 60526219 |
| **HER2-0.HER2-0_S1** | 17-region_10 | 6 | Gain | PPM1D | 17 | 60600183 | 60666280 |
| **HER2-0.HER2-0_S1** | 17-region_10 | 6 | Gain | BCAS3 | 17 | 60677453 | 61392838 |
| **HER2-0.HER2-0_S1** | 17-region_10 | 6 | Gain | BRIP1 | 17 | 61679186 | 61863559 |
| **HER2-0.HER2-0_S1** | 17-region_10 | 6 | Gain | INTS2 | 17 | 61865367 | 61928016 |
| **HER2-0.HER2-0_S1** | 17-region_10 | 6 | Gain | MED13 | 17 | 61942605 | 62065282 |
| **HER2-0.HER2-0_S1** | 17-region_10 | 6 | Gain | METTL2A | 17 | 62423867 | 62450822 |
| **HER2-0.HER2-0_S1** | 17-region_10 | 6 | Gain | TLK2 | 17 | 62458658 | 62615481 |
| **HER2-0.HER2-0_S1** | 17-region_10 | 6 | Gain | TANC2 | 17 | 63009556 | 63427699 |
| **HER2-0.HER2-0_S1** | 17-region_10 | 6 | Gain | AC015923.1 | 17 | 63381231 | 63414312 |
| **HER2-0.HER2-0_S1** | 17-region_10 | 6 | Gain | CYB561 | 17 | 63432304 | 63446378 |
| **HER2-0.HER2-0_S1** | 17-region_10 | 6 | Gain | DCAF7 | 17 | 63550461 | 63594266 |
| **HER2-0.HER2-0_S1** | 17-region_10 | 6 | Gain | MAP3K3 | 17 | 63622415 | 63696303 |
| **HER2-0.HER2-0_S1** | 17-region_10 | 6 | Gain | LIMD2 | 17 | 63695902 | 63701172 |
| **HER2-0.HER2-0_S1** | 17-region_10 | 6 | Gain | STRADA | 17 | 63702832 | 63741970 |
| **HER2-0.HER2-0_S1** | 17-region_10 | 6 | Gain | CCDC47 | 17 | 63745250 | 63776351 |
| **HER2-0.HER2-0_S1** | 17-region_10 | 6 | Gain | DDX42 | 17 | 63773603 | 63819317 |
| **HER2-0.HER2-0_S1** | 17-region_10 | 6 | Gain | FTSJ3 | 17 | 63819433 | 63830012 |
| **HER2-0.HER2-0_S1** | 17-region_10 | 6 | Gain | PSMC5 | 17 | 63827152 | 63832026 |
| **HER2-0.HER2-0_S1** | 17-region_10 | 6 | Gain | SMARCD2 | 17 | 63832081 | 63843065 |
| **HER2-0.HER2-0_S1** | 17-region_10 | 6 | Gain | ERN1 | 17 | 64039142 | 64130819 |
| **HER2-0.HER2-0_S1** | 17-region_10 | 6 | Gain | SNHG25 | 17 | 64145970 | 64146476 |
| **HER2-0.HER2-0_S1** | 17-region_10 | 6 | Gain | TEX2 | 17 | 64147227 | 64263301 |
| **HER2-0.HER2-0_S1** | 17-region_10 | 6 | Gain | PECAM1 | 17 | 64319415 | 64413776 |
| **HER2-0.HER2-0_S1** | 17-region_10 | 6 | Gain | POLG2 | 17 | 64477785 | 64497036 |
| **HER2-0.HER2-0_S1** | 17-region_10 | 6 | Gain | DDX5 | 17 | 64499616 | 64508199 |
| **HER2-0.HER2-0_S1** | 17-region_10 | 6 | Gain | CEP95 | 17 | 64506588 | 64542461 |
| **HER2-0.HER2-0_S1** | 17-region_10 | 6 | Gain | SMURF2 | 17 | 64542295 | 64662068 |
| **HER2-0.HER2-0_S1** | 17-region_10 | 6 | Gain | LRRC37A3 | 17 | 64854312 | 64919480 |
| **HER2-0.HER2-0_S1** | 17-region_10 | 6 | Gain | GNA13 | 17 | 65010715 | 65056839 |
| **HER2-0.HER2-0_S1** | 17-region_10 | 6 | Gain | CEP112 | 17 | 65635538 | 66192084 |
| **HER2-0.HER2-0_S1** | 17-region_10 | 6 | Gain | PRKCA | 17 | 66302636 | 66810743 |
| **HER2-0.HER2-0_S1** | 17-region_10 | 6 | Gain | CACNG4 | 17 | 66964910 | 67033398 |
| **HER2-0.HER2-0_S1** | 17-region_10 | 6 | Gain | HELZ | 17 | 67070438 | 67245989 |
| **HER2-0.HER2-0_S1** | 17-region_10 | 6 | Gain | PSMD12 | 17 | 67337916 | 67366627 |
| **HER2-0.HER2-0_S1** | 17-region_10 | 6 | Gain | PITPNC1 | 17 | 67377281 | 67697261 |
| **HER2-0.HER2-0_S1** | 17-region_10 | 6 | Gain | NOL11 | 17 | 67717833 | 67744531 |
| **HER2-0.HER2-0_S1** | 17-region_10 | 6 | Gain | BPTF | 17 | 67825524 | 67984378 |

| | | | | | | | |
|---|---|---|---|---|---|---|---|
| **HER2-0.HER2-0_S1** | 17-region_10 | 6 | Gain | C17orf58 | 17 | 67991101 | 67996431 |
| **HER2-0.HER2-0_S1** | 17-region_10 | 6 | Gain | KPNA2 | 17 | 68035519 | 68046842 |
| **HER2-0.HER2-0_S1** | 17-region_10 | 6 | Gain | AMZ2 | 17 | 68247574 | 68257164 |
| **HER2-0.HER2-0_S1** | 17-region_10 | 6 | Gain | ARSG | 17 | 68259182 | 68422731 |
| **HER2-0.HER2-0_S1** | 17-region_10 | 6 | Gain | SLC16A6 | 17 | 68267026 | 68291267 |
| **HER2-0.HER2-0_S1** | 17-region_10 | 6 | Gain | WIPI1 | 17 | 68420948 | 68457513 |
| **HER2-0.HER2-0_S1** | 17-region_10 | 6 | Gain | PRKAR1A | 17 | 68511780 | 68551319 |
| **HER2-0.HER2-0_S1** | 17-region_10 | 6 | Gain | ABCA5 | 17 | 69244311 | 69327244 |
| **HER2-0.HER2-0_S1** | 17-region_10 | 6 | Gain | MAP2K6 | 17 | 69414698 | 69543331 |
| **HER2-0.HER2-0_S1** | 17-region_10 | 6 | Gain | SOX9 | 17 | 72121020 | 72126420 |
| **HER2-0.HER2-0_S1** | 17-region_10 | 6 | Gain | LINC00511 | 17 | 72323123 | 72640472 |
| **HER2-0.HER2-0_S1** | 17-region_10 | 6 | Gain | SLC39A11 | 17 | 72645949 | 73092712 |
| **HER2-0.HER2-0_S1** | 17-region_10 | 6 | Gain | COG1 | 17 | 73192632 | 73208507 |
| **HER2-0.HER2-0_S1** | 17-region_10 | 6 | Gain | FAM104A | 17 | 73207353 | 73236753 |
| **HER2-0.HER2-0_S1** | 17-region_10 | 6 | Gain | C17orf80 | 17 | 73232233 | 73248947 |
| **HER2-0.HER2-0_S1** | 17-region_10 | 6 | Gain | CDC42EP4 | 17 | 73283624 | 73312175 |
| **HER2-0.HER2-0_S1** | 17-region_10 | 6 | Gain | SDK2 | 17 | 73334384 | 73644089 |
| **HER2-0.HER2-0_S1** | 17-region_10 | 6 | Gain | RPL38 | 17 | 74203582 | 74210655 |
| **HER2-0.HER2-0_S1** | 17-region_10 | 6 | Gain | GPRC5C | 17 | 74424851 | 74451653 |
| **HER2-0.HER2-0_S1** | 17-region_10 | 6 | Gain | SLC9A3R1 | 17 | 74748613 | 74769353 |
| **HER2-0.HER2-0_S1** | 17-region_10 | 6 | Gain | NAT9 | 17 | 74770547 | 74776367 |
| **HER2-0.HER2-0_S1** | 17-region_10 | 6 | Gain | TMEM104 | 17 | 74776483 | 74839779 |
| **HER2-0.HER2-0_S1** | 17-region_10 | 6 | Gain | HID1 | 17 | 74950743 | 74973166 |
| **HER2-0.HER2-0_S1** | 17-region_10 | 6 | Gain | NT5C | 17 | 75130225 | 75131795 |
| **HER2-0.HER2-0_S1** | 17-region_10 | 6 | Gain | SUMO2 | 17 | 75165586 | 75182983 |
| **HER2-0.HER2-0_S1** | 17-region_10 | 6 | Gain | NUP85 | 17 | 75205659 | 75235758 |
| **HER2-0.HER2-0_S1** | 17-region_10 | 6 | Gain | GGA3 | 17 | 75236599 | 75262363 |
| **HER2-0.HER2-0_S1** | 17-region_10 | 6 | Gain | MRPS7 | 17 | 75261674 | 75266373 |
| **HER2-0.HER2-0_S1** | 17-region_10 | 6 | Gain | GRB2 | 17 | 75318076 | 75405709 |
| **HER2-0.HER2-0_S1** | 17-region_10 | 6 | Gain | TMEM94 | 17 | 75441159 | 75500090 |
| **HER2-0.HER2-0_S1** | 17-region_10 | 6 | Gain | TSEN54 | 17 | 75516060 | 75524739 |
| **HER2-0.HER2-0_S1** | 17-region_10 | 6 | Gain | LLGL2 | 17 | 75525080 | 75575208 |
| **HER2-0.HER2-0_S1** | 17-region_10 | 6 | Gain | MYO15B | 17 | 75588058 | 75626501 |
| **HER2-0.HER2-0_S1** | 17-region_10 | 6 | Gain | RECQL5 | 17 | 75626845 | 75667189 |
| **HER2-0.HER2-0_S1** | 17-region_10 | 6 | Gain | SAP30BP | 17 | 75667116 | 75708062 |
| **HER2-0.HER2-0_S1** | 17-region_10 | 6 | Gain | ITGB4 | 17 | 75721328 | 75757818 |
| **HER2-0.HER2-0_S1** | 17-region_10 | 6 | Gain | H3F3B | 17 | 75776434 | 75785893 |
| **HER2-0.HER2-0_S1** | 17-region_10 | 6 | Gain | UNK | 17 | 75784771 | 75825799 |
| **HER2-0.HER2-0_S1** | 17-region_11 | 6 | Gain | WBP2 | 17 | 75845699 | 75856507 |
| **HER2-0.HER2-0_S1** | 17-region_11 | 6 | Gain | TRIM65 | 17 | 75880335 | 75897003 |

| | | | | | | | |
|---|---|---|---|---|---|---|---|
| **HER2-0.HER2-0_S1** | 17-region_11 | 6 | Gain | MRPL38 | 17 | 75898643 | 75905413 |
| **HER2-0.HER2-0_S1** | 17-region_11 | 6 | Gain | FBF1 | 17 | 75909574 | 75941140 |
| **HER2-0.HER2-0_S1** | 17-region_11 | 6 | Gain | ACOX1 | 17 | 75941507 | 75979363 |
| **HER2-0.HER2-0_S1** | 17-region_11 | 6 | Gain | EVPL | 17 | 76004502 | 76027452 |
| **HER2-0.HER2-0_S1** | 17-region_11 | 6 | Gain | SRP68 | 17 | 76038775 | 76072653 |
| **HER2-0.HER2-0_S1** | 17-region_11 | 6 | Gain | EXOC7 | 17 | 76081017 | 76121576 |
| **HER2-0.HER2-0_S1** | 17-region_11 | 6 | Gain | UBALD2 | 17 | 76265202 | 76271299 |
| **HER2-0.HER2-0_S1** | 17-region_11 | 6 | Gain | PRPSAP1 | 17 | 76309486 | 76384521 |
| **HER2-0.HER2-0_S1** | 17-region_11 | 6 | Gain | UBE2O | 17 | 76389451 | 76453206 |
| **HER2-0.HER2-0_S1** | 17-region_11 | 6 | Gain | RHBDF2 | 17 | 76470891 | 76501790 |
| **HER2-0.HER2-0_S1** | 17-region_11 | 6 | Gain | SNHG16 | 17 | 76557766 | 76565348 |
| **HER2-0.HER2-0_S1** | 17-region_11 | 6 | Gain | ST6GALNAC2 | 17 | 76565379 | 76586956 |
| **HER2-0.HER2-0_S1** | 17-region_12 | 6 | Gain | MXRA7 | 17 | 76672551 | 76711016 |
| **HER2-0.HER2-0_S1** | 17-region_12 | 6 | Gain | METTL23 | 17 | 76726830 | 76733936 |
| **HER2-0.HER2-0_S1** | 17-region_12 | 6 | Gain | SRSF2 | 17 | 76734115 | 76737374 |
| **HER2-0.HER2-0_S1** | 17-region_12 | 6 | Gain | MFSD11 | 17 | 76735865 | 76781449 |
| **HER2-0.HER2-0_S1** | 17-region_12 | 6 | Gain | SEC14L1 | 17 | 77086716 | 77217101 |
| **HER2-0.HER2-0_S1** | 17-region_12 | 6 | Gain | TNRC6C | 17 | 77959240 | 78108835 |
| **HER2-0.HER2-0_S1** | 17-region_12 | 6 | Gain | TMC6 | 17 | 78110458 | 78132407 |
| **HER2-0.HER2-0_S1** | 17-region_12 | 6 | Gain | SYNGR2 | 17 | 78168558 | 78173527 |
| **HER2-0.HER2-0_S1** | 17-region_12 | 6 | Gain | AFMID | 17 | 78187317 | 78207701 |
| **HER2-0.HER2-0_S1** | 17-region_12 | 6 | Gain | PGS1 | 17 | 78378640 | 78425114 |
| **HER2-0.HER2-0_S1** | 17-region_12 | 6 | Gain | CYTH1 | 17 | 78674048 | 78782297 |
| **HER2-0.HER2-0_S1** | 17-region_12 | 6 | Gain | USP36 | 17 | 78787381 | 78841441 |
| **HER2-0.HER2-0_S1** | 17-region_12 | 6 | Gain | TIMP2 | 17 | 78852977 | 78925387 |
| **HER2-0.HER2-0_S1** | 17-region_12 | 6 | Gain | LGALS3BP | 17 | 78971238 | 78980109 |
| **HER2-0.HER2-0_S1** | 17-region_12 | 6 | Gain | CANT1 | 17 | 78991717 | 79009867 |
| **HER2-0.HER2-0_S1** | 17-region_12 | 6 | Gain | ENGASE | 17 | 79074939 | 79088599 |
| **HER2-0.HER2-0_S1** | 17-region_12 | 6 | Gain | CBX4 | 17 | 79833156 | 79839429 |
| **HER2-0.HER2-0_S1** | 17-region_12 | 6 | Gain | TBC1D16 | 17 | 79932343 | 80035848 |
| **HER2-0.HER2-0_S1** | 17-region_12 | 6 | Gain | CCDC40 | 17 | 80036632 | 80100613 |
| **HER2-0.HER2-0_S1** | 17-region_12 | 6 | Gain | GAA | 17 | 80101556 | 80119879 |
| **HER2-0.HER2-0_S1** | 17-region_12 | 6 | Gain | EIF4A3 | 17 | 80135214 | 80147183 |
| **HER2-0.HER2-0_S1** | 17-region_12 | 6 | Gain | CARD14 | 17 | 80169992 | 80209331 |
| **HER2-0.HER2-0_S1** | 17-region_12 | 6 | Gain | SGSH | 17 | 80206716 | 80220923 |
| **HER2-0.HER2-0_S1** | 17-region_12 | 6 | Gain | SLC26A11 | 17 | 80219699 | 80253500 |
| **HER2-0.HER2-0_S1** | 17-region_12 | 6 | Gain | RNF213 | 17 | 80260866 | 80398786 |
| **HER2-0.HER2-0_S1** | 17-region_12 | 6 | Gain | ENDOV | 17 | 80415165 | 80438086 |
| **HER2-0.HER2-0_S1** | 17-region_12 | 6 | Gain | RPTOR | 17 | 80544819 | 80966371 |
| **HER2-0.HER2-0_S1** | 17-region_12 | 6 | Gain | BAIAP2 | 17 | 81035122 | 81117432 |

| | | | | | | | |
|---|---|---|---|---|---|---|---|
| **HER2-0.HER2-0_S1** | 17-region_12 | 6 | Gain | SLC38A10 | 17 | 81245000 | 81295547 |
| **HER2-0.HER2-0_S1** | 17-region_12 | 6 | Gain | ACTG1 | 17 | 81509971 | 81523847 |
| **HER2-0.HER2-0_S1** | 17-region_12 | 6 | Gain | NPLOC4 | 17 | 81556887 | 81648465 |
| **HER2-0.HER2-0_S1** | 17-region_12 | 6 | Gain | OXLD1 | 17 | 81665036 | 81666635 |
| **HER2-0.HER2-0_S1** | 17-region_12 | 6 | Gain | CCDC137 | 17 | 81666364 | 81673904 |
| **HER2-0.HER2-0_S1** | 17-region_12 | 6 | Gain | ARL16 | 17 | 81681174 | 81683924 |
| **HER2-0.HER2-0_S1** | 17-region_12 | 6 | Gain | HGS | 17 | 81683326 | 81703138 |
| **HER2-0.HER2-0_S1** | 17-region_12 | 6 | Gain | MCRIP1 | 17 | 81822361 | 81833302 |
| **HER2-0.HER2-0_S1** | 17-region_12 | 6 | Gain | P4HB | 17 | 81843159 | 81860694 |
| **HER2-0.HER2-0_S1** | 17-region_12 | 6 | Gain | ARHGDIA | 17 | 81867721 | 81871406 |
| **HER2-0.HER2-0_S1** | 17-region_12 | 6 | Gain | ALYREF | 17 | 81887844 | 81891586 |
| **HER2-0.HER2-0_S1** | 17-region_12 | 6 | Gain | ANAPC11 | 17 | 81890790 | 81900991 |
| **HER2-0.HER2-0_S1** | 17-region_12 | 6 | Gain | SIRT7 | 17 | 81911939 | 81921323 |
| **HER2-0.HER2-0_S1** | 17-region_12 | 6 | Gain | PYCR1 | 17 | 81932384 | 81942412 |
| **HER2-0.HER2-0_S1** | 17-region_12 | 6 | Gain | ASPSCR1 | 17 | 81976807 | 82017406 |
| **HER2-0.HER2-0_S1** | 17-region_12 | 6 | Gain | DCXR | 17 | 82035136 | 82037732 |
| **HER2-0.HER2-0_S1** | 17-region_12 | 6 | Gain | RFNG | 17 | 82047902 | 82051831 |
| **HER2-0.HER2-0_S1** | 17-region_12 | 6 | Gain | GPS1 | 17 | 82050691 | 82057470 |
| **HER2-0.HER2-0_S1** | 17-region_12 | 6 | Gain | DUS1L | 17 | 82057506 | 82065887 |
| **HER2-0.HER2-0_S1** | 17-region_12 | 6 | Gain | FASN | 17 | 82078333 | 82098332 |
| **HER2-0.HER2-0_S1** | 17-region_12 | 6 | Gain | CCDC57 | 17 | 82101460 | 82212830 |
| **HER2-0.HER2-0_S1** | 17-region_12 | 6 | Gain | SLC16A3 | 17 | 82228397 | 82261129 |
| **HER2-0.HER2-0_S1** | 17-region_12 | 6 | Gain | CSNK1D | 17 | 82239023 | 82273731 |
| **HER2-0.HER2-0_S1** | 17-region_12 | 6 | Gain | OGFOD3 | 17 | 82389223 | 82418637 |
| **HER2-0.HER2-0_S1** | 17-region_12 | 6 | Gain | NARF | 17 | 82458180 | 82490537 |
| **HER2-0.HER2-0_S1** | 17-region_12 | 6 | Gain | FOXK2 | 17 | 82519713 | 82644662 |
| **HER2-0.HER2-0_S1** | 17-region_12 | 6 | Gain | WDR45B | 17 | 82614562 | 82648553 |
| **HER2-0.HER2-0_S1** | 17-region_12 | 6 | Gain | FN3KRP | 17 | 82716683 | 82730328 |
| **HER2-0.HER2-0_S1** | 17-region_12 | 6 | Gain | FN3K | 17 | 82735575 | 82751197 |
| **HER2-0.HER2-0_S1** | 17-region_12 | 6 | Gain | TBCD | 17 | 82752064 | 82945922 |
| **HER2-0.HER2-0_S1** | 17-region_12 | 6 | Gain | B3GNTL1 | 17 | 82942155 | 83051810 |
| **HER2-0.HER2-0_S1** | 17-region_12 | 6 | Gain | METRNL | 17 | 83079691 | 83095119 |
| **HER2-LOW.HER2-LOW_S1** | 17-region_13 | 1 | Loss | RPH3AL | 17 | 212389 | 386254 |
| **HER2-LOW.HER2-LOW_S1** | 17-region_13 | 1 | Loss | VPS53 | 17 | 508668 | 721717 |
| **HER2-LOW.HER2-LOW_S1** | 17-region_13 | 1 | Loss | GLOD4 | 17 | 757097 | 783390 |
| **HER2-LOW.HER2-LOW_S1** | 17-region_13 | 1 | Loss | ABR | 17 | 1003518 | 1229021 |
| **HER2-LOW.HER2-LOW_S1** | 17-region_13 | 1 | Loss | YWHAE | 17 | 1344272 | 1400378 |
| **HER2-LOW.HER2-LOW_S1** | 17-region_13 | 1 | Loss | CRK | 17 | 1420689 | 1463162 |
| **HER2-LOW.HER2-LOW_S1** | 17-region_13 | 1 | Loss | MYO1C | 17 | 1464098 | 1492812 |
| **HER2-LOW.HER2-LOW_S1** | 17-region_13 | 1 | Loss | PITPNA | 17 | 1517718 | 1562816 |

| | | | | | | | |
|---|---|---|---|---|---|---|---|
| **HER2-LOW.HER2-LOW_S1** | 17-region_13 | 1 | Loss | SLC43A2 | 17 | 1569267 | 1628886 |
| **HER2-LOW.HER2-LOW_S1** | 17-region_13 | 1 | Loss | PRPF8 | 17 | 1650629 | 1684882 |
| **HER2-LOW.HER2-LOW_S1** | 17-region_13 | 1 | Loss | SMYD4 | 17 | 1779485 | 1830634 |
| **HER2-LOW.HER2-LOW_S1** | 17-region_13 | 1 | Loss | RPA1 | 17 | 1829702 | 1900082 |
| **HER2-LOW.HER2-LOW_S1** | 17-region_13 | 1 | Loss | RTN4RL1 | 17 | 1934677 | 2025345 |
| **HER2-LOW.HER2-LOW_S1** | 17-region_13 | 1 | Loss | DPH1 | 17 | 2030110 | 2043430 |
| **HER2-LOW.HER2-LOW_S1** | 17-region_13 | 1 | Loss | SMG6 | 17 | 2059839 | 2303771 |
| **HER2-LOW.HER2-LOW_S1** | 17-region_13 | 1 | Loss | TSR1 | 17 | 2322503 | 2337507 |
| **HER2-LOW.HER2-LOW_S1** | 17-region_13 | 1 | Loss | SGSM2 | 17 | 2337498 | 2381058 |
| **HER2-LOW.HER2-LOW_S1** | 17-region_13 | 1 | Loss | MNT | 17 | 2384060 | 2401118 |
| **HER2-LOW.HER2-LOW_S1** | 17-region_13 | 1 | Loss | METTL16 | 17 | 2405562 | 2511891 |
| **HER2-LOW.HER2-LOW_S1** | 17-region_13 | 1 | Loss | PAFAH1B1 | 17 | 2593210 | 2685615 |
| **HER2-LOW.HER2-LOW_S1** | 17-region_13 | 1 | Loss | CLUH | 17 | 2689386 | 2712663 |
| **HER2-LOW.HER2-LOW_S1** | 17-region_13 | 1 | Loss | RAP1GAP2 | 17 | 2755705 | 3037739 |
| **HER2-LOW.HER2-LOW_S1** | 17-region_13 | 1 | Loss | SHPK | 17 | 3608262 | 3636322 |
| **HER2-LOW.HER2-LOW_S1** | 17-region_13 | 1 | Loss | ITGAE | 17 | 3714628 | 3801243 |
| **HER2-LOW.HER2-LOW_S1** | 17-region_13 | 1 | Loss | NCBP3 | 17 | 3802165 | 3846251 |
| **HER2-LOW.HER2-LOW_S1** | 17-region_13 | 1 | Loss | ATP2A3 | 17 | 3923870 | 3964464 |
| **HER2-LOW.HER2-LOW_S1** | 17-region_13 | 1 | Loss | ZZEF1 | 17 | 4004445 | 4143020 |
| **HER2-LOW.HER2-LOW_S1** | 17-region_13 | 1 | Loss | ANKFY1 | 17 | 4163907 | 4263977 |
| **HER2-LOW.HER2-LOW_S1** | 17-region_13 | 1 | Loss | UBE2G1 | 17 | 4269259 | 4366628 |
| **HER2-LOW.HER2-LOW_S1** | 17-region_13 | 1 | Loss | MYBBP1A | 17 | 4538897 | 4555631 |
| **HER2-LOW.HER2-LOW_S1** | 17-region_13 | 1 | Loss | PELP1 | 17 | 4669774 | 4704337 |
| **HER2-LOW.HER2-LOW_S1** | 17-region_13 | 1 | Loss | PSMB6 | 17 | 4796144 | 4798503 |
| **HER2-LOW.HER2-LOW_S1** | 17-region_13 | 1 | Loss | PLD2 | 17 | 4807096 | 4823434 |
| **HER2-LOW.HER2-LOW_S1** | 17-region_13 | 1 | Loss | MINK1 | 17 | 4833388 | 4898061 |
| **HER2-LOW.HER2-LOW_S1** | 17-region_13 | 1 | Loss | RNF167 | 17 | 4940008 | 4945222 |
| **HER2-LOW.HER2-LOW_S1** | 17-region_13 | 1 | Loss | PFN1 | 17 | 4945652 | 4949061 |
| **HER2-LOW.HER2-LOW_S1** | 17-region_13 | 1 | Loss | CAMTA2 | 17 | 4967992 | 4987652 |
| **HER2-LOW.HER2-LOW_S1** | 17-region_13 | 1 | Loss | KIF1C | 17 | 4997948 | 5028401 |
| **HER2-LOW.HER2-LOW_S1** | 17-region_13 | 1 | Loss | ZNF594 | 17 | 5179536 | 5191883 |
| **HER2-LOW.HER2-LOW_S1** | 17-region_13 | 1 | Loss | RABEP1 | 17 | 5282265 | 5385812 |
| **HER2-LOW.HER2-LOW_S1** | 17-region_13 | 1 | Loss | NUP88 | 17 | 5360963 | 5420160 |
| **HER2-LOW.HER2-LOW_S1** | 17-region_13 | 1 | Loss | RPAIN | 17 | 5419641 | 5432876 |
| **HER2-LOW.HER2-LOW_S1** | 17-region_13 | 1 | Loss | C1QBP | 17 | 5432777 | 5448830 |
| **HER2-LOW.HER2-LOW_S1** | 17-region_13 | 1 | Loss | DERL2 | 17 | 5471251 | 5486811 |
| **HER2-LOW.HER2-LOW_S1** | 17-region_13 | 1 | Loss | KIAA0753 | 17 | 6578148 | 6640927 |
| **HER2-LOW.HER2-LOW_S1** | 17-region_13 | 1 | Loss | XAF1 | 17 | 6755447 | 6775647 |
| **HER2-LOW.HER2-LOW_S1** | 17-region_13 | 1 | Loss | ALOX12-AS1 | 17 | 6876635 | 7012349 |
| **HER2-LOW.HER2-LOW_S1** | 17-region_13 | 1 | Loss | RNASEK | 17 | 7012417 | 7014532 |

| | | | | | | | |
|---|---|---|---|---|---|---|---|
| **HER2-LOW.HER2-LOW_S1** | 17-region_13 | 1 | Loss | ACADVL | 17 | 7217125 | 7225273 |
| **HER2-LOW.HER2-LOW_S1** | 17-region_13 | 1 | Loss | GABARAP | 17 | 7240014 | 7242770 |
| **HER2-LOW.HER2-LOW_S1** | 17-region_13 | 1 | Loss | CTDNEP1 | 17 | 7243591 | 7252491 |
| **HER2-LOW.HER2-LOW_S1** | 17-region_13 | 1 | Loss | CLDN7 | 17 | 7259903 | 7263983 |
| **HER2-LOW.HER2-LOW_S1** | 17-region_13 | 1 | Loss | EIF5A | 17 | 7306999 | 7312463 |
| **HER2-LOW.HER2-LOW_S1** | 17-region_13 | 1 | Loss | GPS2 | 17 | 7311324 | 7315564 |
| **HER2-LOW.HER2-LOW_S1** | 17-region_13 | 1 | Loss | ZBTB4 | 17 | 7459366 | 7484263 |
| **HER2-LOW.HER2-LOW_S1** | 17-region_13 | 1 | Loss | POLR2A | 17 | 7484366 | 7514616 |
| **HER2-LOW.HER2-LOW_S1** | 17-region_13 | 1 | Loss | TNFSF13 | 17 | 7558292 | 7561608 |
| **HER2-LOW.HER2-LOW_S1** | 17-region_13 | 1 | Loss | EIF4A1 | 17 | 7572706 | 7579005 |
| **HER2-LOW.HER2-LOW_S1** | 17-region_13 | 1 | Loss | MPDU1 | 17 | 7583529 | 7592789 |
| **HER2-LOW.HER2-LOW_S1** | 17-region_13 | 1 | Loss | SAT2 | 17 | 7626234 | 7627876 |
| **HER2-LOW.HER2-LOW_S1** | 17-region_13 | 1 | Loss | KDM6B | 17 | 7839904 | 7854796 |
| **HER2-LOW.HER2-LOW_S1** | 17-region_13 | 1 | Loss | NAA38 | 17 | 7856685 | 7885238 |
| **HER2-LOW.HER2-LOW_S1** | 17-region_13 | 1 | Loss | CHD3 | 17 | 7884806 | 7912760 |
| **HER2-LOW.HER2-LOW_S1** | 17-region_13 | 1 | Loss | TRAPPC1 | 17 | 7930345 | 7932123 |
| **HER2-LOW.HER2-LOW_S1** | 17-region_13 | 1 | Loss | PER1 | 17 | 8140472 | 8156506 |
| **HER2-LOW.HER2-LOW_S1** | 17-region_15 | 1 | Loss | KIAA0100 | 17 | 28614440 | 28645454 |
| **HER2-LOW.HER2-LOW_S1** | 17-region_15 | 1 | Loss | SDF2 | 17 | 28648356 | 28662189 |
| **HER2-LOW.HER2-LOW_S1** | 17-region_15 | 1 | Loss | SUPT6H | 17 | 28662091 | 28702684 |
| **HER2-LOW.HER2-LOW_S1** | 17-region_15 | 1 | Loss | RPL23A | 17 | 28719393 | 28724359 |
| **HER2-LOW.HER2-LOW_S1** | 17-region_15 | 1 | Loss | TRAF4 | 17 | 28743984 | 28750958 |
| **HER2-LOW.HER2-LOW_S1** | 17-region_15 | 1 | Loss | FAM222B | 17 | 28755978 | 28855232 |
| **HER2-LOW.HER2-LOW_S1** | 17-region_15 | 1 | Loss | ERAL1 | 17 | 28854938 | 28861067 |
| **HER2-LOW.HER2-LOW_S1** | 17-region_15 | 1 | Loss | FLOT2 | 17 | 28879335 | 28897679 |
| **HER2-LOW.HER2-LOW_S1** | 17-region_15 | 1 | Loss | PHF12 | 17 | 28905250 | 28951771 |
| **HER2-LOW.HER2-LOW_S1** | 17-region_15 | 1 | Loss | MYO18A | 17 | 29071124 | 29180412 |
| **HER2-LOW.HER2-LOW_S1** | 17-region_15 | 1 | Loss | NUFIP2 | 17 | 29255836 | 29294118 |
| **HER2-LOW.HER2-LOW_S1** | 17-region_15 | 1 | Loss | TAOK1 | 17 | 29390464 | 29551904 |
| **HER2-LOW.HER2-LOW_S1** | 17-region_15 | 1 | Loss | TP53I13 | 17 | 29566052 | 29573157 |
| **HER2-LOW.HER2-LOW_S1** | 17-region_15 | 1 | Loss | GIT1 | 17 | 29573469 | 29594054 |
| **HER2-LOW.HER2-LOW_S1** | 17-region_15 | 1 | Loss | CORO6 | 17 | 29614756 | 29622907 |
| **HER2-LOW.HER2-LOW_S1** | 17-region_15 | 1 | Loss | SSH2 | 17 | 29625938 | 29930276 |
| **HER2-LOW.HER2-LOW_S1** | 17-region_15 | 1 | Loss | NSRP1 | 17 | 30115521 | 30186475 |
| **HER2-LOW.HER2-LOW_S1** | 17-region_15 | 1 | Loss | BLMH | 17 | 30248195 | 30292056 |
| **HER2-LOW.HER2-LOW_S1** | 17-region_15 | 1 | Loss | CPD | 17 | 30378905 | 30469989 |
| **HER2-LOW.HER2-LOW_S1** | 17-region_15 | 1 | Loss | GOSR1 | 17 | 30477362 | 30527592 |
| **HER2-LOW.HER2-LOW_S1** | 17-region_15 | 1 | Loss | AC005562.1 | 17 | 30576464 | 30672789 |
| **HER2-LOW.HER2-LOW_S1** | 17-region_15 | 1 | Loss | CRLF3 | 17 | 30769388 | 30824776 |
| **HER2-LOW.HER2-LOW_S1** | 17-region_15 | 1 | Loss | ATAD5 | 17 | 30831970 | 30895869 |

| | | | | | | | |
|---|---|---|---|---|---|---|---|
| **HER2-LOW.HER2-LOW_S1** | 17-region_15 | 1 | Loss | TEFM | 17 | 30897336 | 30906820 |
| **HER2-LOW.HER2-LOW_S1** | 17-region_15 | 1 | Loss | RNF135 | 17 | 30968785 | 30999911 |
| **HER2-LOW.HER2-LOW_S1** | 17-region_15 | 1 | Loss | NF1 | 17 | 31094927 | 31382116 |
| **HER2-LOW.HER2-LOW_S1** | 17-region_15 | 1 | Loss | RAB11FIP4 | 17 | 31391624 | 31538217 |
| **HER2-LOW.HER2-LOW_S1** | 17-region_15 | 1 | Loss | UTP6 | 17 | 31860899 | 31901765 |
| **HER2-LOW.HER2-LOW_S1** | 17-region_15 | 1 | Loss | SUZ12 | 17 | 31937018 | 32001045 |
| **HER2-LOW.HER2-LOW_S1** | 17-region_15 | 1 | Loss | RHOT1 | 17 | 32142454 | 32253374 |
| **HER2-LOW.HER2-LOW_S1** | 17-region_15 | 1 | Loss | C17orf75 | 17 | 32324565 | 32350023 |
| **HER2-LOW.HER2-LOW_S1** | 17-region_15 | 1 | Loss | ZNF207 | 17 | 32350109 | 32381886 |
| **HER2-LOW.HER2-LOW_S1** | 17-region_15 | 1 | Loss | PSMD11 | 17 | 32444261 | 32483318 |
| **HER2-LOW.HER2-LOW_S1** | 17-region_15 | 1 | Loss | MYO1D | 17 | 32492522 | 32877177 |
| **HER2-LOW.HER2-LOW_S1** | 17-region_15 | 1 | Loss | ASIC2 | 17 | 33013087 | 34174964 |
| **HER2-LOW.HER2-LOW_S1** | 17-region_15 | 1 | Loss | ZNF830 | 17 | 34961530 | 34963775 |
| **HER2-LOW.HER2-LOW_S1** | 17-region_15 | 1 | Loss | LIG3 | 17 | 34980494 | 35009743 |
| **HER2-LOW.HER2-LOW_S1** | 17-region_15 | 1 | Loss | RFFL | 17 | 35005990 | 35089319 |
| **HER2-LOW.HER2-LOW_S1** | 17-region_15 | 1 | Loss | SLFN5 | 17 | 35243036 | 35273655 |
| **HER2-LOW.HER2-LOW_S1** | 17-region_15 | 1 | Loss | AP2B1 | 17 | 35578046 | 35726409 |
| **HER2-LOW.HER2-LOW_S1** | 17-region_15 | 1 | Loss | TAF15 | 17 | 35713791 | 35864615 |
| **HER2-LOW.HER2-LOW_S1** | 17-region_15 | 1 | Loss | MYO19 | 17 | 36495633 | 36543435 |
| **HER2-LOW.HER2-LOW_S1** | 17-region_17 | 1 | Loss | PCGF2 | 17 | 38733897 | 38749817 |
| **HER2-LOW.HER2-LOW_S1** | 17-region_17 | 1 | Loss | PSMB3 | 17 | 38752736 | 38764231 |
| **HER2-LOW.HER2-LOW_S1** | 17-region_17 | 1 | Loss | PIP4K2B | 17 | 38765689 | 38800126 |
| **HER2-LOW.HER2-LOW_S1** | 17-region_17 | 1 | Loss | CWC25 | 17 | 38800434 | 38825481 |
| **HER2-LOW.HER2-LOW_S1** | 17-region_17 | 1 | Loss | RPL23 | 17 | 38847865 | 38853843 |
| **HER2-LOW.HER2-LOW_S1** | 17-region_17 | 1 | Loss | LASP1 | 17 | 38869859 | 38921770 |
| **HER2-LOW.HER2-LOW_S1** | 17-region_17 | 1 | Loss | RPL19 | 17 | 39200283 | 39204727 |
| **HER2-LOW.HER2-LOW_S1** | 17-region_17 | 1 | Loss | FBXL20 | 17 | 39252644 | 39402523 |
| **HER2-LOW.HER2-LOW_S1** | 17-region_17 | 1 | Loss | MED1 | 17 | 39404285 | 39451286 |
| **HER2-LOW.HER2-LOW_S1** | 17-region_17 | 1 | Loss | CDK12 | 17 | 39461511 | 39564907 |
| **HER2-LOW.HER2-LOW_S1** | 17-region_18 | 6 | Gain | STARD3 | 17 | 39637065 | 39663484 |
| **HER2-LOW.HER2-LOW_S1** | 17-region_18 | 6 | Gain | PGAP3 | 17 | 39671122 | 39696797 |
| **HER2-LOW.HER2-LOW_S1** | 17-region_18 | 6 | Gain | ERBB2 | 17 | 39687914 | 39730426 |
| **HER2-LOW.HER2-LOW_S1** | 17-region_18 | 6 | Gain | MIEN1 | 17 | 39728496 | 39730787 |
| **HER2-LOW.HER2-LOW_S1** | 17-region_18 | 6 | Gain | GRB7 | 17 | 39737927 | 39747291 |
| **HER2-LOW.HER2-LOW_S1** | 17-region_18 | 6 | Gain | IKZF3 | 17 | 39757715 | 39864188 |
| **HER2-LOW.HER2-LOW_S1** | 17-region_18 | 6 | Gain | ORMDL3 | 17 | 39921041 | 39927601 |
| **HER2-LOW.HER2-LOW_S1** | 17-region_18 | 6 | Gain | PSMD3 | 17 | 39980797 | 39997960 |
| **HER2-LOW.HER2-LOW_S1** | 17-region_18 | 6 | Gain | MED24 | 17 | 40019097 | 40061215 |
| **HER2-LOW.HER2-LOW_S1** | 17-region_18 | 6 | Gain | THRA | 17 | 40058290 | 40093867 |
| **HER2-LOW.HER2-LOW_S1** | 17-region_18 | 6 | Gain | MSL1 | 17 | 40122298 | 40136916 |

| | | | | | | | |
|---|---|---|---|---|---|---|---|
| **HER2-LOW.HER2-LOW_S1** | 17-region_18 | 6 | Gain | CASC3 | 17 | 40140318 | 40172183 |
| **HER2-LOW.HER2-LOW_S1** | 17-region_18 | 6 | Gain | WIPF2 | 17 | 40219304 | 40284136 |
| **HER2-LOW.HER2-LOW_S1** | 17-region_18 | 6 | Gain | RARA | 17 | 40309192 | 40357643 |
| **HER2-LOW.HER2-LOW_S1** | 17-region_19 | 6 | Gain | TOP2A | 17 | 40388516 | 40417950 |
| **HER2-LOW.HER2-LOW_S1** | 17-region_19 | 6 | Gain | IGFBP4 | 17 | 40443461 | 40457731 |
| **HER2-LOW.HER2-LOW_S1** | 17-region_19 | 6 | Gain | TNS4 | 17 | 40475828 | 40501597 |
| **HER2-LOW.HER2-LOW_S1** | 17-region_19 | 6 | Gain | SMARCE1 | 17 | 40624962 | 40648508 |
| **HER2-LOW.HER2-LOW_S1** | 17-region_19 | 6 | Gain | KRT10 | 17 | 40818117 | 40822595 |
| **HER2-LOW.HER2-LOW_S1** | 17-region_19 | 6 | Gain | KRT23 | 17 | 40922696 | 40937634 |
| **HER2-LOW.HER2-LOW_S1** | 17-region_19 | 6 | Gain | KRT15 | 17 | 41513743 | 41522529 |
| **HER2-LOW.HER2-LOW_S1** | 17-region_19 | 6 | Gain | KRT19 | 17 | 41523617 | 41528308 |
| **HER2-LOW.HER2-LOW_S1** | 17-region_19 | 6 | Gain | EIF1 | 17 | 41688885 | 41692668 |
| **HER2-LOW.HER2-LOW_S1** | 17-region_19 | 6 | Gain | JUP | 17 | 41754604 | 41786931 |
| **HER2-LOW.HER2-LOW_S1** | 17-region_19 | 6 | Gain | P3H4 | 17 | 41801947 | 41812604 |
| **HER2-LOW.HER2-LOW_S1** | 17-region_19 | 6 | Gain | FKBP10 | 17 | 41812680 | 41823217 |
| **HER2-LOW.HER2-LOW_S1** | 17-region_19 | 6 | Gain | NT5C3B | 17 | 41825181 | 41836263 |
| **HER2-LOW.HER2-LOW_S1** | 17-region_19 | 6 | Gain | ACLY | 17 | 41866908 | 41930542 |
| **HER2-LOW.HER2-LOW_S1** | 17-region_19 | 6 | Gain | DNAJC7 | 17 | 41976433 | 42021376 |
| **HER2-LOW.HER2-LOW_S1** | 17-region_19 | 6 | Gain | KAT2A | 17 | 42113108 | 42121358 |
| **HER2-LOW.HER2-LOW_S1** | 17-region_19 | 6 | Gain | RAB5C | 17 | 42124976 | 42155044 |
| **HER2-LOW.HER2-LOW_S1** | 17-region_19 | 6 | Gain | STAT5B | 17 | 42199168 | 42276707 |
| **HER2-LOW.HER2-LOW_S1** | 17-region_19 | 6 | Gain | STAT3 | 17 | 42313324 | 42388568 |
| **HER2-LOW.HER2-LOW_S1** | 17-region_19 | 6 | Gain | ATP6V0A1 | 17 | 42458844 | 42522611 |
| **HER2-LOW.HER2-LOW_S1** | 17-region_19 | 6 | Gain | NAGLU | 17 | 42536172 | 42544449 |
| **HER2-LOW.HER2-LOW_S1** | 17-region_19 | 6 | Gain | HSD17B1 | 17 | 42549214 | 42555213 |
| **HER2-LOW.HER2-LOW_S1** | 17-region_19 | 6 | Gain | COASY | 17 | 42561467 | 42566277 |
| **HER2-LOW.HER2-LOW_S1** | 17-region_19 | 6 | Gain | MLX | 17 | 42567068 | 42573239 |
| **HER2-LOW.HER2-LOW_S1** | 17-region_19 | 6 | Gain | TUBG2 | 17 | 42659305 | 42667006 |
| **HER2-LOW.HER2-LOW_S1** | 17-region_19 | 6 | Gain | PLEKHH3 | 17 | 42667914 | 42676994 |
| **HER2-LOW.HER2-LOW_S1** | 17-region_20 | 6 | Gain | EZH1 | 17 | 42700275 | 42745049 |
| **HER2-LOW.HER2-LOW_S1** | 17-region_20 | 6 | Gain | VPS25 | 17 | 42773436 | 42779599 |
| **HER2-LOW.HER2-LOW_S1** | 17-region_20 | 6 | Gain | WNK4 | 17 | 42780678 | 42796936 |
| **HER2-LOW.HER2-LOW_S1** | 17-region_20 | 6 | Gain | COA3 | 17 | 42795147 | 42798704 |
| **HER2-LOW.HER2-LOW_S1** | 17-region_20 | 6 | Gain | BECN1 | 17 | 42810134 | 42833350 |
| **HER2-LOW.HER2-LOW_S1** | 17-region_20 | 6 | Gain | PSME3 | 17 | 42824385 | 42843758 |
| **HER2-LOW.HER2-LOW_S1** | 17-region_20 | 6 | Gain | AARSD1 | 17 | 42950526 | 42964498 |
| **HER2-LOW.HER2-LOW_S1** | 17-region_20 | 6 | Gain | RUNDC1 | 17 | 42980565 | 42993690 |
| **HER2-LOW.HER2-LOW_S1** | 17-region_20 | 6 | Gain | RPL27 | 17 | 42998273 | 43002959 |
| **HER2-LOW.HER2-LOW_S1** | 17-region_20 | 6 | Gain | IFI35 | 17 | 43006725 | 43014456 |
| **HER2-LOW.HER2-LOW_S1** | 17-region_20 | 6 | Gain | VAT1 | 17 | 43014605 | 43025123 |

| | | | | | | | |
|---|---|---|---|---|---|---|---|
| **HER2-LOW.HER2-LOW_S1** | 17-region_20 | 6 | Gain | BRCA1 | 17 | 43044295 | 43170245 |
| **HER2-LOW.HER2-LOW_S1** | 17-region_20 | 6 | Gain | NBR2 | 17 | 43125610 | 43153671 |
| **HER2-LOW.HER2-LOW_S1** | 17-region_20 | 6 | Gain | NBR1 | 17 | 43170481 | 43211689 |
| **HER2-LOW.HER2-LOW_S1** | 17-region_20 | 6 | Gain | DHX8 | 17 | 43483865 | 43544463 |
| **HER2-LOW.HER2-LOW_S1** | 17-region_20 | 6 | Gain | DUSP3 | 17 | 43766121 | 43778988 |
| **HER2-LOW.HER2-LOW_S1** | 17-region_20 | 6 | Gain | MPP3 | 17 | 43800799 | 43833170 |
| **HER2-LOW.HER2-LOW_S1** | 17-region_21 | 6 | Gain | TMEM101 | 17 | 44011188 | 44023946 |
| **HER2-LOW.HER2-LOW_S1** | 17-region_21 | 6 | Gain | LSM12 | 17 | 44034635 | 44067619 |
| **HER2-LOW.HER2-LOW_S1** | 17-region_21 | 6 | Gain | G6PC3 | 17 | 44070730 | 44076344 |
| **HER2-LOW.HER2-LOW_S1** | 17-region_21 | 6 | Gain | HDAC5 | 17 | 44076746 | 44123702 |
| **HER2-LOW.HER2-LOW_S1** | 17-region_21 | 6 | Gain | ASB16-AS1 | 17 | 44175973 | 44186717 |
| **HER2-LOW.HER2-LOW_S1** | 17-region_21 | 6 | Gain | TMUB2 | 17 | 44186970 | 44191731 |
| **HER2-LOW.HER2-LOW_S1** | 17-region_21 | 6 | Gain | ATXN7L3 | 17 | 44191805 | 44200113 |
| **HER2-LOW.HER2-LOW_S1** | 17-region_21 | 6 | Gain | UBTF | 17 | 44205033 | 44221626 |
| **HER2-LOW.HER2-LOW_S1** | 17-region_21 | 6 | Gain | SLC25A39 | 17 | 44319625 | 44324870 |
| **HER2-LOW.HER2-LOW_S1** | 17-region_21 | 6 | Gain | GRN | 17 | 44345086 | 44353102 |
| **HER2-LOW.HER2-LOW_S1** | 17-region_23 | 1 | Loss | NME1 | 17 | 51153536 | 51162428 |
| **HER2-LOW.HER2-LOW_S1** | 17-region_23 | 1 | Loss | NME2 | 17 | 51165435 | 51171747 |
| **HER2-LOW.HER2-LOW_S1** | 17-region_23 | 1 | Loss | MBTD1 | 17 | 51177425 | 51260163 |
| **HER2-LOW.HER2-LOW_S1** | 17-region_23 | 1 | Loss | UTP18 | 17 | 51260528 | 51297936 |
| **HER2-LOW.HER2-LOW_S1** | 17-region_23 | 1 | Loss | TOM1L1 | 17 | 54899387 | 54961956 |
| **HER2-LOW.HER2-LOW_S1** | 17-region_23 | 1 | Loss | COX11 | 17 | 54951902 | 54968785 |
| **HER2-LOW.HER2-LOW_S1** | 17-region_23 | 1 | Loss | STXBP4 | 17 | 54968727 | 55173632 |
| **HER2-LOW.HER2-LOW_S1** | 17-region_23 | 1 | Loss | ANKFN1 | 17 | 55882301 | 56511659 |
| **HER2-LOW.HER2-LOW_S1** | 17-region_23 | 1 | Loss | C17orf67 | 17 | 56791913 | 56838773 |
| **HER2-LOW.HER2-LOW_S1** | 17-region_23 | 1 | Loss | DGKE | 17 | 56834099 | 56869567 |
| **HER2-LOW.HER2-LOW_S1** | 17-region_23 | 1 | Loss | TRIM25 | 17 | 56887909 | 56914038 |
| **HER2-LOW.HER2-LOW_S1** | 17-region_23 | 1 | Loss | COIL | 17 | 56938187 | 56961054 |
| **HER2-LOW.HER2-LOW_S1** | 17-region_23 | 1 | Loss | SCPEP1 | 17 | 56978105 | 57006768 |
| **HER2-LOW.HER2-LOW_S1** | 17-region_23 | 1 | Loss | AKAP1 | 17 | 57085092 | 57121349 |
| **HER2-LOW.HER2-LOW_S1** | 17-region_23 | 1 | Loss | MSI2 | 17 | 57255851 | 57684685 |
| **HER2-LOW.HER2-LOW_S1** | 17-region_23 | 1 | Loss | MRPS23 | 17 | 57834781 | 57850056 |
| **HER2-LOW.HER2-LOW_S1** | 17-region_23 | 1 | Loss | CUEDC1 | 17 | 57861243 | 57955323 |
| **HER2-LOW.HER2-LOW_S1** | 17-region_23 | 1 | Loss | VEZF1 | 17 | 57971547 | 57988259 |
| **HER2-LOW.HER2-LOW_S1** | 17-region_23 | 1 | Loss | SRSF1 | 17 | 58003360 | 58007346 |
| **HER2-LOW.HER2-LOW_S1** | 17-region_23 | 1 | Loss | DYNLL2 | 17 | 58083415 | 58095536 |
| **HER2-LOW.HER2-LOW_S1** | 17-region_23 | 1 | Loss | MKS1 | 17 | 58205436 | 58219605 |
| **HER2-LOW.HER2-LOW_S1** | 17-region_23 | 1 | Loss | TSPOAP1 | 17 | 58301228 | 58328760 |
| **HER2-LOW.HER2-LOW_S1** | 17-region_23 | 1 | Loss | TSPOAP1-AS1 | 17 | 58325450 | 58415766 |
| **HER2-LOW.HER2-LOW_S1** | 17-region_23 | 1 | Loss | SUPT4H1 | 17 | 58345175 | 58353093 |

| | | | | | | | |
|---|---|---|---|---|---|---|---|
| **HER2-LOW.HER2-LOW_S1** | 17-region_23 | 1 | Loss | RNF43 | 17 | 58352500 | 58417595 |
| **HER2-LOW.HER2-LOW_S1** | 17-region_23 | 1 | Loss | MTMR4 | 17 | 58489529 | 58517905 |
| **HER2-LOW.HER2-LOW_S1** | 17-region_23 | 1 | Loss | TEX14 | 17 | 58556678 | 58692055 |
| **HER2-LOW.HER2-LOW_S1** | 17-region_23 | 1 | Loss | RAD51C | 17 | 58692573 | 58735611 |
| **HER2-LOW.HER2-LOW_S1** | 17-region_23 | 1 | Loss | PPM1E | 17 | 58755869 | 58985176 |
| **HER2-LOW.HER2-LOW_S1** | 17-region_23 | 1 | Loss | TRIM37 | 17 | 58982638 | 59106921 |
| **HER2-LOW.HER2-LOW_S1** | 17-region_23 | 1 | Loss | SKA2 | 17 | 59109951 | 59155269 |
| **HER2-LOW.HER2-LOW_S1** | 17-region_23 | 1 | Loss | GDPD1 | 17 | 59220467 | 59275967 |
| **HER2-LOW.HER2-LOW_S1** | 17-region_23 | 1 | Loss | YPEL2 | 17 | 59331689 | 59401729 |
| **HER2-LOW.HER2-LOW_S1** | 17-region_23 | 1 | Loss | DHX40 | 17 | 59565525 | 59608345 |
| **HER2-LOW.HER2-LOW_S1** | 17-region_23 | 1 | Loss | CLTC | 17 | 59619689 | 59696956 |
| **HER2-LOW.HER2-LOW_S1** | 17-region_23 | 1 | Loss | PTRH2 | 17 | 59674636 | 59707626 |
| **HER2-LOW.HER2-LOW_S1** | 17-region_23 | 1 | Loss | VMP1 | 17 | 59707192 | 59842255 |
| **HER2-LOW.HER2-LOW_S1** | 17-region_23 | 1 | Loss | TUBD1 | 17 | 59859482 | 59892945 |
| **HER2-LOW.HER2-LOW_S1** | 17-region_23 | 1 | Loss | RPS6KB1 | 17 | 59893046 | 59950564 |
| **HER2-LOW.HER2-LOW_S1** | 17-region_23 | 1 | Loss | RNFT1 | 17 | 59952240 | 59964761 |
| **HER2-LOW.HER2-LOW_S1** | 17-region_23 | 1 | Loss | HEATR6 | 17 | 60043194 | 60078931 |
| **HER2-LOW.HER2-LOW_S1** | 17-region_25 | 1 | Loss | CCDC47 | 17 | 63745250 | 63776351 |
| **HER2-LOW.HER2-LOW_S1** | 17-region_25 | 1 | Loss | DDX42 | 17 | 63773603 | 63819317 |
| **HER2-LOW.HER2-LOW_S1** | 17-region_25 | 1 | Loss | FTSJ3 | 17 | 63819433 | 63830012 |
| **HER2-LOW.HER2-LOW_S1** | 17-region_25 | 1 | Loss | PSMC5 | 17 | 63827152 | 63832026 |
| **HER2-LOW.HER2-LOW_S1** | 17-region_25 | 1 | Loss | SMARCD2 | 17 | 63832081 | 63843065 |
| **HER2-LOW.HER2-LOW_S1** | 17-region_25 | 1 | Loss | ERN1 | 17 | 64039142 | 64130819 |
| **HER2-LOW.HER2-LOW_S1** | 17-region_25 | 1 | Loss | SNHG25 | 17 | 64145970 | 64146476 |
| **HER2-LOW.HER2-LOW_S1** | 17-region_25 | 1 | Loss | TEX2 | 17 | 64147227 | 64263301 |
| **HER2-LOW.HER2-LOW_S1** | 17-region_25 | 1 | Loss | PECAM1 | 17 | 64319415 | 64413776 |
| **HER2-LOW.HER2-LOW_S1** | 17-region_25 | 1 | Loss | POLG2 | 17 | 64477785 | 64497036 |
| **HER2-LOW.HER2-LOW_S1** | 17-region_25 | 1 | Loss | DDX5 | 17 | 64499616 | 64508199 |
| **HER2-LOW.HER2-LOW_S1** | 17-region_25 | 1 | Loss | CEP95 | 17 | 64506588 | 64542461 |
| **HER2-LOW.HER2-LOW_S1** | 17-region_25 | 1 | Loss | SMURF2 | 17 | 64542295 | 64662068 |
| **HER2-LOW.HER2-LOW_S1** | 17-region_25 | 1 | Loss | LRRC37A3 | 17 | 64854312 | 64919480 |
| **HER2-LOW.HER2-LOW_S1** | 17-region_25 | 1 | Loss | GNA13 | 17 | 65010715 | 65056839 |
| **HER2-LOW.HER2-LOW_S1** | 17-region_25 | 1 | Loss | CEP112 | 17 | 65635538 | 66192084 |
| **HER2-LOW.HER2-LOW_S1** | 17-region_25 | 1 | Loss | PRKCA | 17 | 66302636 | 66810743 |
| **HER2-LOW.HER2-LOW_S1** | 17-region_25 | 1 | Loss | CACNG4 | 17 | 66964910 | 67033398 |
| **HER2-LOW.HER2-LOW_S1** | 17-region_25 | 1 | Loss | HELZ | 17 | 67070438 | 67245989 |
| **HER2-LOW.HER2-LOW_S1** | 17-region_25 | 1 | Loss | PSMD12 | 17 | 67337916 | 67366627 |
| **HER2-LOW.HER2-LOW_S1** | 17-region_25 | 1 | Loss | PITPNC1 | 17 | 67377281 | 67697261 |
| **HER2-LOW.HER2-LOW_S1** | 17-region_25 | 1 | Loss | NOL11 | 17 | 67717833 | 67744531 |
| **HER2-LOW.HER2-LOW_S1** | 17-region_25 | 1 | Loss | BPTF | 17 | 67825524 | 67984378 |

| | | | | | | | |
|---|---|---|---|---|---|---|---|
| **HER2-LOW.HER2-LOW_S1** | 17-region_25 | 1 | Loss | C17orf58 | 17 | 67991101 | 67996431 |
| **HER2-LOW.HER2-LOW_S1** | 17-region_25 | 1 | Loss | KPNA2 | 17 | 68035519 | 68046842 |
| **HER2-LOW.HER2-LOW_S1** | 17-region_25 | 1 | Loss | AMZ2 | 17 | 68247574 | 68257164 |
| **HER2-LOW.HER2-LOW_S1** | 17-region_25 | 1 | Loss | ARSG | 17 | 68259182 | 68422731 |
| **HER2-LOW.HER2-LOW_S1** | 17-region_25 | 1 | Loss | SLC16A6 | 17 | 68267026 | 68291267 |
| **HER2-LOW.HER2-LOW_S1** | 17-region_25 | 1 | Loss | WIPI1 | 17 | 68420948 | 68457513 |
| **HER2-LOW.HER2-LOW_S1** | 17-region_25 | 1 | Loss | PRKAR1A | 17 | 68511780 | 68551319 |
| **HER2-LOW.HER2-LOW_S1** | 17-region_25 | 1 | Loss | ABCA5 | 17 | 69244311 | 69327244 |
| **HER2-LOW.HER2-LOW_S1** | 17-region_25 | 1 | Loss | MAP2K6 | 17 | 69414698 | 69543331 |
| **HER2-LOW.HER2-LOW_S1** | 17-region_25 | 1 | Loss | SOX9 | 17 | 72121020 | 72126420 |
| **HER2-LOW.HER2-LOW_S1** | 17-region_25 | 1 | Loss | LINC00511 | 17 | 72323123 | 72640472 |
| **HER2-LOW.HER2-LOW_S1** | 17-region_25 | 1 | Loss | SLC39A11 | 17 | 72645949 | 73092712 |
| **HER2-LOW.HER2-LOW_S1** | 17-region_25 | 1 | Loss | COG1 | 17 | 73192632 | 73208507 |
| **HER2-LOW.HER2-LOW_S1** | 17-region_25 | 1 | Loss | FAM104A | 17 | 73207353 | 73236753 |
| **HER2-LOW.HER2-LOW_S1** | 17-region_25 | 1 | Loss | C17orf80 | 17 | 73232233 | 73248947 |
| **HER2-LOW.HER2-LOW_S1** | 17-region_25 | 1 | Loss | CDC42EP4 | 17 | 73283624 | 73312175 |
| **HER2-LOW.HER2-LOW_S1** | 17-region_25 | 1 | Loss | SDK2 | 17 | 73334384 | 73644089 |
| **HER2-LOW.HER2-LOW_S1** | 17-region_25 | 1 | Loss | RPL38 | 17 | 74203582 | 74210655 |
| **HER2-LOW.HER2-LOW_S1** | 17-region_25 | 1 | Loss | GPRC5C | 17 | 74424851 | 74451653 |
| **HER2-LOW.HER2-LOW_S1** | 17-region_25 | 1 | Loss | SLC9A3R1 | 17 | 74748613 | 74769353 |
| **HER2-LOW.HER2-LOW_S1** | 17-region_25 | 1 | Loss | NAT9 | 17 | 74770547 | 74776367 |
| **HER2-LOW.HER2-LOW_S1** | 17-region_25 | 1 | Loss | TMEM104 | 17 | 74776483 | 74839779 |
| **HER2-LOW.HER2-LOW_S1** | 17-region_25 | 1 | Loss | HID1 | 17 | 74950743 | 74973166 |
| **HER2-LOW.HER2-LOW_S1** | 17-region_25 | 1 | Loss | NT5C | 17 | 75130225 | 75131795 |
| **HER2-LOW.HER2-LOW_S1** | 17-region_25 | 1 | Loss | SUMO2 | 17 | 75165586 | 75182983 |
| **HER2-LOW.HER2-LOW_S1** | 17-region_27 | 6 | Gain | MFSD11 | 17 | 76735865 | 76781449 |
| **HER2-LOW.HER2-LOW_S1** | 17-region_27 | 6 | Gain | SEC14L1 | 17 | 77086716 | 77217101 |
| **HER2-LOW.HER2-LOW_S1** | 17-region_27 | 6 | Gain | TNRC6C | 17 | 77959240 | 78108835 |
| **HER2-LOW.HER2-LOW_S1** | 17-region_27 | 6 | Gain | TMC6 | 17 | 78110458 | 78132407 |
| **HER2-LOW.HER2-LOW_S1** | 17-region_27 | 6 | Gain | SYNGR2 | 17 | 78168558 | 78173527 |
| **HER2-LOW.HER2-LOW_S1** | 17-region_27 | 6 | Gain | AFMID | 17 | 78187317 | 78207701 |
| **HER2-LOW.HER2-LOW_S1** | 17-region_27 | 6 | Gain | PGS1 | 17 | 78378640 | 78425114 |
| **HER2-LOW.HER2-LOW_S1** | 17-region_27 | 6 | Gain | CYTH1 | 17 | 78674048 | 78782297 |
| **HER2-LOW.HER2-LOW_S1** | 17-region_27 | 6 | Gain | USP36 | 17 | 78787381 | 78841441 |
| **HER2-LOW.HER2-LOW_S1** | 17-region_27 | 6 | Gain | TIMP2 | 17 | 78852977 | 78925387 |
| **HER2-LOW.HER2-LOW_S1** | 17-region_27 | 6 | Gain | LGALS3BP | 17 | 78971238 | 78980109 |
| **HER2-LOW.HER2-LOW_S1** | 17-region_27 | 6 | Gain | CANT1 | 17 | 78991717 | 79009867 |
| **HER2-LOW.HER2-LOW_S1** | 17-region_27 | 6 | Gain | ENGASE | 17 | 79074939 | 79088599 |
| **HER2-LOW.HER2-LOW_S1** | 17-region_27 | 6 | Gain | CBX4 | 17 | 79833156 | 79839429 |
| **HER2-LOW.HER2-LOW_S1** | 17-region_27 | 6 | Gain | TBC1D16 | 17 | 79932343 | 80035848 |

| | | | | | | | |
|---|---|---|---|---|---|---|---|
| **HER2-LOW.HER2-LOW_S1** | 17-region_27 | 6 | Gain | CCDC40 | 17 | 80036632 | 80100613 |
| **HER2-LOW.HER2-LOW_S1** | 17-region_27 | 6 | Gain | GAA | 17 | 80101556 | 80119879 |
| **HER2-LOW.HER2-LOW_S1** | 17-region_27 | 6 | Gain | EIF4A3 | 17 | 80135214 | 80147183 |
| **HER2-LOW.HER2-LOW_S1** | 17-region_28 | 6 | Gain | CARD14 | 17 | 80169992 | 80209331 |
| **HER2-LOW.HER2-LOW_S1** | 17-region_28 | 6 | Gain | SGSH | 17 | 80206716 | 80220923 |
| **HER2-LOW.HER2-LOW_S1** | 17-region_28 | 6 | Gain | SLC26A11 | 17 | 80219699 | 80253500 |
| **HER2-LOW.HER2-LOW_S1** | 17-region_28 | 6 | Gain | RNF213 | 17 | 80260866 | 80398786 |
| **HER2-LOW.HER2-LOW_S1** | 17-region_28 | 6 | Gain | ENDOV | 17 | 80415165 | 80438086 |
| **HER2-LOW.HER2-LOW_S1** | 17-region_28 | 6 | Gain | RPTOR | 17 | 80544819 | 80966371 |
| **HER2-LOW.HER2-LOW_S1** | 17-region_28 | 6 | Gain | BAIAP2 | 17 | 81035122 | 81117432 |
| **HER2-LOW.HER2-LOW_S1** | 17-region_28 | 6 | Gain | SLC38A10 | 17 | 81245000 | 81295547 |
| **HER2-LOW.HER2-LOW_S1** | 17-region_28 | 6 | Gain | ACTG1 | 17 | 81509971 | 81523847 |
| **HER2-LOW.HER2-LOW_S1** | 17-region_28 | 6 | Gain | NPLOC4 | 17 | 81556887 | 81648465 |
| **HER2-LOW.HER2-LOW_S1** | 17-region_28 | 6 | Gain | OXLD1 | 17 | 81665036 | 81666635 |
| **HER2-LOW.HER2-LOW_S1** | 17-region_28 | 6 | Gain | CCDC137 | 17 | 81666364 | 81673904 |
| **HER2-LOW.HER2-LOW_S1** | 17-region_28 | 6 | Gain | ARL16 | 17 | 81681174 | 81683924 |
| **HER2-LOW.HER2-LOW_S1** | 17-region_28 | 6 | Gain | HGS | 17 | 81683326 | 81703138 |
| **HER2-LOW.HER2-LOW_S1** | 17-region_28 | 6 | Gain | MCRIP1 | 17 | 81822361 | 81833302 |
| **HER2-LOW.HER2-LOW_S1** | 17-region_28 | 6 | Gain | P4HB | 17 | 81843159 | 81860694 |
| **HER2-LOW.HER2-LOW_S1** | 17-region_28 | 6 | Gain | ARHGDIA | 17 | 81867721 | 81871406 |
| **HER2-LOW.HER2-LOW_S1** | 17-region_28 | 6 | Gain | ALYREF | 17 | 81887844 | 81891586 |
| **HER2-LOW.HER2-LOW_S1** | 17-region_28 | 6 | Gain | ANAPC11 | 17 | 81890790 | 81900991 |
| **HER2-LOW.HER2-LOW_S1** | 17-region_28 | 6 | Gain | SIRT7 | 17 | 81911939 | 81921323 |
| **HER2-LOW.HER2-LOW_S1** | 17-region_28 | 6 | Gain | PYCR1 | 17 | 81932384 | 81942412 |
| **HER2-LOW.HER2-LOW_S1** | 17-region_28 | 6 | Gain | ASPSCR1 | 17 | 81976807 | 82017406 |
| **HER2-LOW.HER2-LOW_S1** | 17-region_28 | 6 | Gain | DCXR | 17 | 82035136 | 82037732 |
| **HER2-LOW.HER2-LOW_S1** | 17-region_28 | 6 | Gain | RFNG | 17 | 82047902 | 82051831 |
| **HER2-LOW.HER2-LOW_S1** | 17-region_28 | 6 | Gain | GPS1 | 17 | 82050691 | 82057470 |
| **HER2-LOW.HER2-LOW_S1** | 17-region_28 | 6 | Gain | DUS1L | 17 | 82057506 | 82065887 |
| **HER2-LOW.HER2-LOW_S1** | 17-region_28 | 6 | Gain | FASN | 17 | 82078333 | 82098332 |
| **HER2-LOW.HER2-LOW_S1** | 17-region_28 | 6 | Gain | CCDC57 | 17 | 82101460 | 82212830 |
| **HER2-LOW.HER2-LOW_S1** | 17-region_28 | 6 | Gain | SLC16A3 | 17 | 82228397 | 82261129 |
| **HER2-LOW.HER2-LOW_S1** | 17-region_28 | 6 | Gain | CSNK1D | 17 | 82239023 | 82273731 |
| **HER2-LOW.HER2-LOW_S1** | 17-region_28 | 6 | Gain | OGFOD3 | 17 | 82389223 | 82418637 |
| **HER2-LOW.HER2-LOW_S1** | 17-region_28 | 6 | Gain | NARF | 17 | 82458180 | 82490537 |
| **HER2-LOW.HER2-LOW_S1** | 17-region_28 | 6 | Gain | FOXK2 | 17 | 82519713 | 82644662 |
| **HER2-LOW.HER2-LOW_S1** | 17-region_28 | 6 | Gain | WDR45B | 17 | 82614562 | 82648553 |
| **HER2-LOW.HER2-LOW_S1** | 17-region_28 | 6 | Gain | FN3KRP | 17 | 82716683 | 82730328 |
| **HER2-LOW.HER2-LOW_S1** | 17-region_28 | 6 | Gain | FN3K | 17 | 82735575 | 82751197 |
| **HER2-LOW.HER2-LOW_S1** | 17-region_28 | 6 | Gain | TBCD | 17 | 82752064 | 82945922 |

| | | | | | | | |
|---|---|---|---|---|---|---|---|
| **HER2-LOW.HER2-LOW_S1** | 17-region_28 | 6 | Gain | B3GNTL1 | 17 | 82942155 | 83051810 |
| **HER2-LOW.HER2-LOW_S1** | 17-region_28 | 6 | Gain | METRNL | 17 | 83079691 | 83095119 |

**Supplementary table 4. Top 100 differentially expressed genes in tumor-associated macrophage (TAM) clusters.**

| | p_val | avg_log2FC | pct.1 | pct.2 | p_val_adj | cluster | gene |
|---|---|---|---|---|---|---|---|
| **1** | 0 | 1.39867614 | 0.919 | 0.585 | 0 | 0 | DOCK4 |
| **2** | 4.67E-262 | 1.29971706 | 0.872 | 0.52 | 1.25E-257 | 0 | FMNL2 |
| **3** | 1.08E-226 | 1.00984423 | 0.94 | 0.706 | 2.88E-222 | 0 | PLXDC2 |
| **4** | 2.33E-184 | 1.19782748 | 0.687 | 0.341 | 6.21E-180 | 0 | DLEU1 |
| **5** | 2.05E-179 | 1.07030627 | 0.809 | 0.472 | 5.46E-175 | 0 | FRMD4A |
| **6** | 9.62E-145 | 1.05889932 | 0.678 | 0.37 | 2.57E-140 | 0 | MSR1 |
| **7** | 6.16E-143 | 1.11662697 | 0.682 | 0.399 | 1.64E-138 | 0 | TANC2 |
| **8** | 1.70E-106 | 1.12698223 | 0.368 | 0.135 | 4.54E-102 | 0 | OLR1 |
| **9** | 4.39E-99 | 1.01711828 | 0.407 | 0.169 | 1.17E-94 | 0 | CADM1 |
| **10** | 3.22E-97 | 1.02357435 | 0.589 | 0.347 | 8.59E-93 | 0 | ELL2 |
| **11** | 2.06E-91 | 1.312168 | 0.27 | 0.084 | 5.50E-87 | 0 | AC011586.2 |
| **12** | 1.00E-87 | 1.02508046 | 0.401 | 0.181 | 2.67E-83 | 0 | TPRG1 |
| **13** | 1.57E-82 | 1.79794102 | 0.199 | 0.049 | 4.19E-78 | 0 | AC093895.1 |
| **14** | 1.30E-79 | 1.07458477 | 0.298 | 0.113 | 3.47E-75 | 0 | MACC1 |
| **15** | 4.94E-72 | 1.2296076 | 0.379 | 0.186 | 1.32E-67 | 0 | ZNF331 |
| **16** | 1.94E-58 | 1.03914348 | 0.259 | 0.108 | 5.18E-54 | 0 | KCNQ3 |
| **17** | 2.73E-92 | 1.18027318 | 0.16 | 0.022 | 7.28E-88 | 1 | FLT3 |
| **18** | 5.88E-85 | 1.3039153 | 0.303 | 0.101 | 1.57E-80 | 1 | JAML |
| **19** | 1.12E-73 | 1.22027145 | 0.23 | 0.066 | 2.98E-69 | 1 | AL034397.3 |
| **20** | 1.47E-64 | 1.41614491 | 0.405 | 0.208 | 3.93E-60 | 1 | CCSER1 |
| **21** | 2.29E-59 | 1.37870038 | 0.522 | 0.338 | 6.12E-55 | 1 | HDAC9 |
| **22** | 6.52E-50 | 1.00849062 | 0.163 | 0.047 | 1.74E-45 | 1 | P2RY14 |
| **23** | 9.07E-46 | 1.44455944 | 0.374 | 0.207 | 2.42E-41 | 1 | AFF3 |
| **24** | 4.71E-43 | 1.08208831 | 0.444 | 0.284 | 1.26E-38 | 1 | STK17B |
| **25** | 6.85E-38 | 1.07330613 | 0.466 | 0.324 | 1.83E-33 | 1 | RTN1 |
| **26** | 3.63E-17 | 1.05026183 | 0.281 | 0.186 | 9.69E-13 | 1 | MUCL1 |
| **27** | 0 | 4.13734174 | 0.828 | 0.144 | 0 | 2 | F13A1 |
| **28** | 0 | 3.03756467 | 0.804 | 0.188 | 0 | 2 | MRC1 |
| **29** | 2.35E-297 | 2.91332803 | 0.571 | 0.093 | 6.28E-293 | 2 | CD163L1 |
| **30** | 8.79E-221 | 1.9908288 | 0.831 | 0.35 | 2.35E-216 | 2 | CD163 |
| **31** | 7.45E-204 | 1.96314647 | 0.682 | 0.216 | 1.99E-199 | 2 | MS4A4A |
| **32** | 9.57E-203 | 1.73959599 | 0.332 | 0.036 | 2.55E-198 | 2 | LILRB5 |
| **33** | 5.31E-197 | 2.08173123 | 0.698 | 0.242 | 1.42E-192 | 2 | RGL1 |
| **34** | 1.51E-184 | 2.01469011 | 0.755 | 0.329 | 4.04E-180 | 2 | PDGFC |
| **35** | 5.80E-176 | 1.91453025 | 0.194 | 0.008 | 1.55E-171 | 2 | LYVE1 |

| | | | | | | | |
|---|---|---|---|---|---|---|---|
| **36** | 1.24E-175 | 1.76120018 | 0.799 | 0.38 | 3.30E-171 | 2 | SLC9A9 |
| **37** | 1.90E-172 | 1.90632396 | 0.682 | 0.248 | 5.08E-168 | 2 | MTSS1 |
| **38** | 4.58E-171 | 1.87658906 | 0.266 | 0.026 | 1.22E-166 | 2 | MPPED2 |
| **39** | 1.43E-161 | 1.69019388 | 0.628 | 0.211 | 3.81E-157 | 2 | STAB1 |
| **40** | 1.79E-155 | 1.66203211 | 0.712 | 0.303 | 4.79E-151 | 2 | ITSN1 |
| **41** | 3.25E-143 | 2.01776359 | 0.38 | 0.079 | 8.69E-139 | 2 | COLEC12 |
| **42** | 2.24E-137 | 1.54469926 | 0.622 | 0.241 | 5.98E-133 | 2 | AP2A2 |
| **43** | 6.75E-137 | 1.76367891 | 0.553 | 0.18 | 1.80E-132 | 2 | CSGALNACT1 |
| **44** | 5.20E-132 | 1.41073432 | 0.782 | 0.418 | 1.39E-127 | 2 | SLCO2B1 |
| **45** | 1.00E-131 | 2.24305872 | 0.699 | 0.343 | 2.68E-127 | 2 | RBPJ |
| **46** | 3.54E-131 | 1.62905973 | 0.701 | 0.331 | 9.45E-127 | 2 | STARD13 |
| **47** | 1.70E-128 | 1.45519831 | 0.821 | 0.474 | 4.55E-124 | 2 | FMN1 |
| **48** | 5.54E-125 | 1.62306464 | 0.51 | 0.162 | 1.48E-120 | 2 | SELENOP |
| **49** | 1.21E-124 | 1.80960128 | 0.703 | 0.356 | 3.23E-120 | 2 | MAN1A1 |
| **50** | 1.80E-121 | 1.53458434 | 0.193 | 0.019 | 4.82E-117 | 2 | MAMDC2 |
| **51** | 2.30E-121 | 1.31441601 | 0.79 | 0.44 | 6.15E-117 | 2 | FRMD4B |
| **52** | 1.24E-114 | 1.41074465 | 0.617 | 0.265 | 3.31E-110 | 2 | TBC1D14 |
| **53** | 3.63E-114 | 1.63313075 | 0.4 | 0.109 | 9.69E-110 | 2 | PLTP |
| **54** | 1.75E-110 | 1.56308122 | 0.307 | 0.064 | 4.68E-106 | 2 | AL162414.1 |
| **55** | 1.38E-107 | 1.77927892 | 0.552 | 0.211 | 3.70E-103 | 2 | PDK4 |
| **56** | 4.47E-105 | 1.22668956 | 0.768 | 0.418 | 1.19E-100 | 2 | MERTK |
| **57** | 5.44E-102 | 1.27167609 | 0.812 | 0.526 | 1.45E-97 | 2 | FCHSD2 |
| **58** | 2.56E-99 | 1.27714396 | 0.278 | 0.057 | 6.83E-95 | 2 | WLS |
| **59** | 1.62E-97 | 1.34170642 | 0.619 | 0.292 | 4.33E-93 | 2 | PEAK1 |
| **60** | 1.81E-95 | 1.27471064 | 0.65 | 0.315 | 4.85E-91 | 2 | SGMS1 |
| **61** | 9.36E-95 | 1.54118353 | 0.247 | 0.047 | 2.50E-90 | 2 | IL2RA |
| **62** | 8.19E-94 | 1.39897814 | 0.269 | 0.056 | 2.19E-89 | 2 | EDA |
| **63** | 1.97E-92 | 1.12968399 | 0.145 | 0.014 | 5.26E-88 | 2 | SCN9A |
| **64** | 5.77E-92 | 1.29733091 | 0.596 | 0.265 | 1.54E-87 | 2 | NRP1 |
| **65** | 2.85E-90 | 1.08478474 | 0.787 | 0.457 | 7.60E-86 | 2 | MS4A6A |
| **66** | 2.83E-89 | 1.54930788 | 0.292 | 0.071 | 7.55E-85 | 2 | AC100849.1 |
| **67** | 1.25E-88 | 1.0241996 | 0.203 | 0.033 | 3.33E-84 | 2 | FOLR2 |
| **68** | 5.63E-87 | 1.24059125 | 0.619 | 0.314 | 1.50E-82 | 2 | ANKS1A |
| **69** | 2.59E-85 | 1.26435705 | 0.428 | 0.15 | 6.90E-81 | 2 | ME1 |
| **70** | 3.91E-84 | 1.11131058 | 0.673 | 0.349 | 1.04E-79 | 2 | MS4A7 |
| **71** | 2.24E-83 | 1.20414919 | 0.521 | 0.223 | 5.99E-79 | 2 | DAB2 |
| **72** | 3.80E-83 | 1.04951603 | 0.789 | 0.534 | 1.01E-78 | 2 | ITPR2 |
| **73** | 7.50E-82 | 1.14182263 | 0.217 | 0.041 | 2.00E-77 | 2 | CR1 |
| **74** | 1.07E-80 | 1.19928574 | 0.322 | 0.092 | 2.85E-76 | 2 | SLC22A23 |

| | | | | | | | |
|---|---|---|---|---|---|---|---|
| **75** | 1.63E-79 | 1.38068418 | 0.513 | 0.227 | 4.34E-75 | 2 | SH3BP5 |
| **76** | 3.23E-78 | 1.6971662 | 0.221 | 0.046 | 8.62E-74 | 2 | TTN |
| **77** | 2.51E-77 | 1.1305176 | 0.274 | 0.069 | 6.71E-73 | 2 | IGF1 |
| **78** | 8.76E-77 | 1.0826823 | 0.586 | 0.275 | 2.34E-72 | 2 | MS4A4E |
| **79** | 2.06E-76 | 1.48506213 | 0.336 | 0.106 | 5.51E-72 | 2 | NAV2 |
| **80** | 7.06E-72 | 1.07937637 | 0.684 | 0.386 | 1.89E-67 | 2 | CPM |
| **81** | 2.15E-71 | 1.40776961 | 0.172 | 0.03 | 5.74E-67 | 2 | CCDC141 |
| **82** | 2.35E-67 | 1.05320971 | 0.722 | 0.462 | 6.28E-63 | 2 | ARHGAP18 |
| **83** | 7.47E-65 | 1.06833122 | 0.28 | 0.083 | 2.00E-60 | 2 | VSIG4 |
| **84** | 3.21E-63 | 1.40349913 | 0.332 | 0.117 | 8.57E-59 | 2 | PID1 |
| **85** | 1.22E-62 | 1.19967517 | 0.456 | 0.21 | 3.25E-58 | 2 | WWP1 |
| **86** | 1.84E-59 | 1.66896108 | 0.187 | 0.043 | 4.91E-55 | 2 | FGF13 |
| **87** | 5.54E-58 | 1.04682649 | 0.686 | 0.447 | 1.48E-53 | 2 | MGAT5 |
| **88** | 1.35E-54 | 1.08024117 | 0.524 | 0.28 | 3.61E-50 | 2 | SNX9 |
| **89** | 2.26E-53 | 1.15538245 | 0.273 | 0.093 | 6.05E-49 | 2 | RCAN1 |
| **90** | 2.47E-51 | 1.01876417 | 0.414 | 0.189 | 6.60E-47 | 2 | TNFRSF21 |
| **91** | 9.37E-51 | 1.19957021 | 0.215 | 0.062 | 2.50E-46 | 2 | RNF150 |
| **92** | 9.07E-42 | 1.0398293 | 0.333 | 0.149 | 2.42E-37 | 2 | FAM20A |
| **93** | 4.04E-35 | 1.08951347 | 0.247 | 0.1 | 1.08E-30 | 2 | PAPSS2 |
| **94** | 2.21E-24 | 1.02208827 | 0.337 | 0.192 | 5.91E-20 | 2 | TFRC |
| **95** | 4.73E-104 | 1.90597601 | 0.742 | 0.509 | 1.26E-99 | 3 | RPLP1 |
| **96** | 1.38E-92 | 2.19897764 | 0.735 | 0.525 | 3.69E-88 | 3 | RPL41 |
| **97** | 2.37E-75 | 1.53033692 | 0.715 | 0.568 | 6.33E-71 | 3 | EEF1A1 |
| **98** | 7.00E-72 | 1.86794092 | 0.583 | 0.349 | 1.87E-67 | 3 | RPL13 |
| **99** | 4.80E-70 | 2.25497078 | 0.488 | 0.237 | 1.28E-65 | 3 | RPL32 |
| **100** | 3.24E-62 | 2.16957845 | 0.57 | 0.365 | 8.66E-58 | 3 | RPL28 |
| **101** | 8.71E-62 | 1.80680614 | 0.532 | 0.308 | 2.33E-57 | 3 | RPLP0 |
| **102** | 1.70E-59 | 1.6111104 | 0.669 | 0.542 | 4.55E-55 | 3 | RPL10 |
| **103** | 4.96E-59 | 1.58905972 | 0.537 | 0.318 | 1.33E-54 | 3 | RPL8 |
| **104** | 1.66E-58 | 2.90967957 | 0.397 | 0.174 | 4.42E-54 | 3 | RPL39 |
| **105** | 2.37E-58 | 1.63507953 | 0.575 | 0.368 | 6.32E-54 | 3 | RPS8 |
| **106** | 7.98E-58 | 1.82007001 | 0.493 | 0.268 | 2.13E-53 | 3 | RPS15 |
| **107** | 1.87E-56 | 1.67612953 | 0.592 | 0.412 | 4.99E-52 | 3 | RPS18 |
| **108** | 2.21E-55 | 1.77961256 | 0.509 | 0.289 | 5.90E-51 | 3 | RPL18A |
| **109** | 9.95E-53 | 1.67833998 | 0.499 | 0.284 | 2.66E-48 | 3 | RPL19 |
| **110** | 7.03E-52 | 1.85171449 | 0.512 | 0.313 | 1.88E-47 | 3 | RPL30 |
| **111** | 9.17E-52 | 1.00838289 | 0.827 | 0.82 | 2.45E-47 | 3 | B2M |
| **112** | 2.37E-51 | 1.45333647 | 0.523 | 0.323 | 6.33E-47 | 3 | RPS4X |
| **113** | 2.76E-51 | 1.9622932 | 0.416 | 0.203 | 7.37E-47 | 3 | RPL36 |

| | | | | | | | |
|---|---|---|---|---|---|---|---|
| **114** | 5.12E-51 | 2.04477223 | 0.472 | 0.271 | 1.37E-46 | 3 | RPS12 |
| **115** | 3.09E-49 | 1.82193365 | 0.507 | 0.311 | 8.25E-45 | 3 | RPS19 |
| **116** | 4.24E-49 | 2.3516049 | 0.441 | 0.24 | 1.13E-44 | 3 | RPS27 |
| **117** | 6.52E-47 | 2.09654747 | 0.367 | 0.172 | 1.74E-42 | 3 | RPL34 |
| **118** | 1.69E-46 | 1.16435181 | 0.776 | 0.694 | 4.52E-42 | 3 | MT-CO2 |
| **119** | 2.00E-45 | 1.30711371 | 0.583 | 0.434 | 5.33E-41 | 3 | ACTG1 |
| **120** | 4.12E-45 | 1.87295638 | 0.431 | 0.241 | 1.10E-40 | 3 | RPL29 |
| **121** | 4.61E-45 | 1.10197115 | 0.709 | 0.665 | 1.23E-40 | 3 | ACTB |
| **122** | 2.17E-44 | 1.6915327 | 0.403 | 0.208 | 5.80E-40 | 3 | RPS5 |
| **123** | 3.33E-44 | 1.60820075 | 0.476 | 0.289 | 8.88E-40 | 3 | RPL18 |
| **124** | 5.13E-44 | 1.81926285 | 0.43 | 0.235 | 1.37E-39 | 3 | RPS13 |
| **125** | 1.30E-43 | 2.08551091 | 0.435 | 0.246 | 3.47E-39 | 3 | RPS15A |
| **126** | 1.60E-43 | 1.57666092 | 0.504 | 0.32 | 4.28E-39 | 3 | RPS3 |
| **127** | 2.86E-43 | 1.51622058 | 0.51 | 0.339 | 7.64E-39 | 3 | RPL11 |
| **128** | 3.95E-43 | 2.08845849 | 0.661 | 0.556 | 1.05E-38 | 3 | MTRNR2L12 |
| **129** | 2.61E-42 | 1.91811791 | 0.335 | 0.154 | 6.96E-38 | 3 | RPL35A |
| **130** | 1.74E-40 | 2.0076246 | 0.411 | 0.229 | 4.66E-36 | 3 | RPS21 |
| **131** | 1.80E-39 | 1.42621463 | 0.461 | 0.285 | 4.80E-35 | 3 | RPS3A |
| **132** | 2.41E-39 | 2.16462406 | 0.444 | 0.276 | 6.43E-35 | 3 | RPS28 |
| **133** | 3.58E-39 | 1.39362001 | 0.51 | 0.346 | 9.55E-35 | 3 | RPS2 |
| **134** | 1.28E-38 | 1.48106384 | 0.474 | 0.304 | 3.41E-34 | 3 | GAPDH |
| **135** | 4.32E-38 | 1.57431216 | 0.476 | 0.31 | 1.15E-33 | 3 | RPS24 |
| **136** | 1.89E-36 | 1.67862623 | 0.438 | 0.276 | 5.04E-32 | 3 | RPL12 |
| **137** | 2.71E-36 | 1.6739604 | 0.342 | 0.172 | 7.25E-32 | 3 | RPLP2 |
| **138** | 4.66E-36 | 1.66405212 | 0.359 | 0.187 | 1.24E-31 | 3 | FAU |
| **139** | 1.59E-35 | 1.85517182 | 0.402 | 0.237 | 4.26E-31 | 3 | RPS14 |
| **140** | 7.10E-35 | 1.37324481 | 0.52 | 0.393 | 1.90E-30 | 3 | TMSB4X |
| **141** | 6.61E-34 | 1.66075317 | 0.394 | 0.234 | 1.76E-29 | 3 | RPS23 |
| **142** | 9.00E-34 | 1.65644934 | 0.324 | 0.165 | 2.40E-29 | 3 | PPDPF |
| **143** | 2.86E-32 | 1.44441858 | 0.37 | 0.216 | 7.63E-28 | 3 | HSPB1 |
| **144** | 2.76E-30 | 1.34773685 | 0.476 | 0.345 | 7.36E-26 | 3 | RPL7A |
| **145** | 5.93E-30 | 2.51167078 | 0.2 | 0.076 | 1.58E-25 | 3 | RPS29 |
| **146** | 6.45E-30 | 1.2318046 | 0.616 | 0.53 | 1.72E-25 | 3 | FTL |
| **147** | 7.51E-30 | 1.29032466 | 0.476 | 0.339 | 2.01E-25 | 3 | RPL3 |
| **148** | 1.11E-29 | 1.80699833 | 0.321 | 0.173 | 2.96E-25 | 3 | RPS26 |
| **149** | 1.15E-29 | 1.34193699 | 0.357 | 0.204 | 3.06E-25 | 3 | RPL13A |
| **150** | 2.09E-29 | 1.43727793 | 0.367 | 0.217 | 5.57E-25 | 3 | RPL14 |
| **151** | 2.64E-28 | 1.45828911 | 0.312 | 0.165 | 7.06E-24 | 3 | RPL37A |
| **152** | 8.69E-28 | 1.52963888 | 0.345 | 0.201 | 2.32E-23 | 3 | RPL26 |

| | | | | | | | |
|---|---|---|---|---|---|---|---|
| **153** | 3.98E-27 | 1.39992504 | 0.365 | 0.223 | 1.06E-22 | 3 | RPL15 |
| **154** | 4.51E-27 | 1.39338732 | 0.359 | 0.217 | 1.20E-22 | 3 | RPS16 |
| **155** | 5.45E-27 | 1.00433161 | 0.6 | 0.533 | 1.46E-22 | 3 | MT-ND4L |
| **156** | 5.71E-27 | 1.41273816 | 0.306 | 0.164 | 1.52E-22 | 3 | RPL23A |
| **157** | 4.04E-25 | 1.68170431 | 0.485 | 0.394 | 1.08E-20 | 3 | TMSB10 |
| **158** | 4.83E-25 | 1.84908653 | 0.312 | 0.177 | 1.29E-20 | 3 | COX6C |
| **159** | 5.53E-25 | 1.76415227 | 0.345 | 0.212 | 1.48E-20 | 3 | RPL37 |
| **160** | 5.89E-25 | 1.42974682 | 0.34 | 0.202 | 1.57E-20 | 3 | MIF |
| **161** | 1.61E-24 | 1.4486535 | 0.419 | 0.294 | 4.29E-20 | 3 | KRT19 |
| **162** | 2.95E-24 | 1.26908938 | 0.263 | 0.132 | 7.88E-20 | 3 | S100A6 |
| **163** | 4.77E-24 | 1.36796149 | 0.502 | 0.397 | 1.27E-19 | 3 | MT-ND4 |
| **164** | 3.53E-23 | 1.55925775 | 0.23 | 0.104 | 9.43E-19 | 3 | MTRNR2L8 |
| **165** | 6.71E-23 | 1.44934268 | 0.211 | 0.096 | 1.79E-18 | 3 | ATP5F1E |
| **166** | 8.04E-23 | 1.23842694 | 0.381 | 0.255 | 2.15E-18 | 3 | RPS27A |
| **167** | 8.10E-23 | 1.38398859 | 0.282 | 0.156 | 2.16E-18 | 3 | RPSA |
| **168** | 1.68E-22 | 1.43312906 | 0.175 | 0.072 | 4.50E-18 | 3 | RPL38 |
| **169** | 2.09E-22 | 1.16702207 | 0.406 | 0.294 | 5.57E-18 | 3 | PTMA |
| **170** | 2.17E-22 | 1.35989167 | 0.265 | 0.142 | 5.79E-18 | 3 | COX6A1 |
| **171** | 4.13E-22 | 1.20782897 | 0.449 | 0.353 | 1.10E-17 | 3 | TPT1 |
| **172** | 6.94E-22 | 1.19008244 | 0.403 | 0.288 | 1.85E-17 | 3 | EEF2 |
| **173** | 9.12E-22 | 1.28617197 | 0.263 | 0.141 | 2.43E-17 | 3 | RPL36A |
| **174** | 2.24E-20 | 1.26036162 | 0.268 | 0.149 | 5.97E-16 | 3 | RPL24 |
| **175** | 3.10E-20 | 1.38046055 | 0.291 | 0.173 | 8.28E-16 | 3 | NDUFC2 |
| **176** | 8.72E-20 | 1.25226132 | 0.198 | 0.093 | 2.33E-15 | 3 | RPL27 |
| **177** | 2.05E-19 | 1.10987713 | 0.22 | 0.111 | 5.47E-15 | 3 | RPL7 |
| **178** | 4.42E-18 | 1.28693334 | 0.23 | 0.122 | 1.18E-13 | 3 | LGALS1 |
| **179** | 1.18E-17 | 1.21882998 | 0.169 | 0.076 | 3.16E-13 | 3 | COX7C |
| **180** | 3.64E-17 | 1.09042746 | 0.617 | 0.6 | 9.73E-13 | 3 | MGP |
| **181** | 4.82E-17 | 1.16505371 | 0.15 | 0.064 | 1.29E-12 | 3 | COX8A |
| **182** | 9.73E-17 | 1.18815682 | 0.224 | 0.122 | 2.60E-12 | 3 | NME2 |
| **183** | 1.03E-16 | 1.01848601 | 0.466 | 0.4 | 2.76E-12 | 3 | XBP1 |
| **184** | 1.81E-16 | 1.25531249 | 0.208 | 0.11 | 4.84E-12 | 3 | RPS11 |
| **185** | 2.20E-16 | 1.1661318 | 0.362 | 0.266 | 5.88E-12 | 3 | S100A11 |
| **186** | 2.75E-16 | 1.38474549 | 0.191 | 0.097 | 7.33E-12 | 3 | UBA52 |
| **187** | 5.46E-16 | 1.30912374 | 0.244 | 0.143 | 1.46E-11 | 3 | IGFBP5 |
| **188** | 3.22E-15 | 1.10993963 | 0.288 | 0.186 | 8.59E-11 | 3 | KRT8 |
| **189** | 3.49E-15 | 1.08948764 | 0.198 | 0.106 | 9.31E-11 | 3 | PRDX1 |
| **190** | 4.56E-15 | 1.12743311 | 0.339 | 0.25 | 1.22E-10 | 3 | PFN1 |
| **191** | 8.38E-15 | 1.14228302 | 0.34 | 0.251 | 2.24E-10 | 3 | RPS6 |

| | | | | | | | |
|---|---|---|---|---|---|---|---|
| **192** | 1.79E-14 | 1.0934645 | 0.269 | 0.172 | 4.79E-10 | 3 | EEF1G |
| **193** | 3.72E-14 | 1.14957526 | 0.392 | 0.313 | 9.94E-10 | 3 | MT-ND2 |
| **194** | 7.22E-14 | 1.3582224 | 0.189 | 0.102 | 1.93E-09 | 3 | THSD4 |
| **195** | 0 | 3.15037904 | 0.674 | 0.045 | 0 | 4 | PRKG1 |
| **196** | 0 | 2.81347276 | 0.761 | 0.094 | 0 | 4 | CALD1 |
| **197** | 0 | 2.45225482 | 0.557 | 0.029 | 0 | 4 | BICC1 |
| **198** | 0 | 2.30626005 | 0.507 | 0.021 | 0 | 4 | ZFPM2 |
| **199** | 0 | 2.26088084 | 0.552 | 0.029 | 0 | 4 | DLC1 |
| **200** | 0 | 2.17356364 | 0.44 | 0.016 | 0 | 4 | NOX4 |
| **201** | 0 | 2.12464752 | 0.592 | 0.046 | 0 | 4 | FBXL7 |
| **202** | 0 | 2.11521507 | 0.496 | 0.024 | 0 | 4 | CACNA1C |
| **203** | 0 | 2.08157442 | 0.401 | 0.013 | 0 | 4 | SGIP1 |
| **204** | 0 | 2.0447952 | 0.462 | 0.022 | 0 | 4 | SGCD |
| **205** | 0 | 2.0319275 | 0.602 | 0.043 | 0 | 4 | PTPRG |
| **206** | 0 | 2.01951156 | 0.446 | 0.022 | 0 | 4 | PCDH7 |
| **207** | 0 | 1.94616158 | 0.504 | 0.029 | 0 | 4 | PRICKLE2 |
| **208** | 0 | 1.94579877 | 0.469 | 0.023 | 0 | 4 | PDZRN3 |
| **209** | 0 | 1.75283418 | 0.459 | 0.013 | 0 | 4 | MEG3 |
| **210** | 0 | 1.71175564 | 0.414 | 0.016 | 0 | 4 | ZNF521 |
| **211** | 0 | 1.70470046 | 0.422 | 0.013 | 0 | 4 | PRKD1 |
| **212** | 7.13E-304 | 1.94210065 | 0.462 | 0.025 | 1.90E-299 | 4 | SLIT2 |
| **213** | 1.05E-297 | 2.58105 | 0.443 | 0.023 | 2.79E-293 | 4 | GPC6 |
| **214** | 1.27E-295 | 2.0132525 | 0.52 | 0.036 | 3.40E-291 | 4 | CASC15 |
| **215** | 4.23E-293 | 2.02377605 | 0.427 | 0.022 | 1.13E-288 | 4 | SUGCT |
| **216** | 1.02E-290 | 1.30983964 | 0.337 | 0.01 | 2.73E-286 | 4 | MEIS2 |
| **217** | 5.59E-288 | 2.04567981 | 0.544 | 0.042 | 1.49E-283 | 4 | TSHZ2 |
| **218** | 7.59E-287 | 2.12896092 | 0.363 | 0.013 | 2.03E-282 | 4 | KCND2 |
| **219** | 2.77E-282 | 2.7366515 | 0.52 | 0.041 | 7.40E-278 | 4 | LAMA2 |
| **220** | 1.27E-279 | 1.78602054 | 0.472 | 0.03 | 3.38E-275 | 4 | PRRX1 |
| **221** | 4.71E-279 | 1.34603371 | 0.366 | 0.014 | 1.26E-274 | 4 | CACNA2D1 |
| **222** | 2.81E-278 | 1.31579293 | 0.305 | 0.008 | 7.51E-274 | 4 | MEG8 |
| **223** | 8.47E-278 | 1.82714692 | 0.554 | 0.046 | 2.26E-273 | 4 | FAP |
| **224** | 1.62E-276 | 1.78445695 | 0.427 | 0.023 | 4.32E-272 | 4 | ADAMTS12 |
| **225** | 1.50E-271 | 1.80859446 | 0.403 | 0.02 | 4.01E-267 | 4 | LDB2 |
| **226** | 2.57E-271 | 1.95079161 | 0.459 | 0.03 | 6.87E-267 | 4 | LMCD1 |
| **227** | 9.26E-267 | 1.72963944 | 0.393 | 0.02 | 2.47E-262 | 4 | EBF1 |
| **228** | 1.70E-266 | 1.04734012 | 0.294 | 0.007 | 4.55E-262 | 4 | NR2F1-AS1 |
| **229** | 3.93E-254 | 1.36014552 | 0.385 | 0.02 | 1.05E-249 | 4 | MIR100HG |
| **230** | 3.36E-250 | 2.25302232 | 0.581 | 0.062 | 8.97E-246 | 4 | CDH11 |

| | | | | | | | |
|---|---|---|---|---|---|---|---|
| **231** | 1.10E-248 | 1.23658895 | 0.329 | 0.013 | 2.95E-244 | 4 | ZFHX4 |
| **232** | 6.43E-243 | 1.54638806 | 0.509 | 0.045 | 1.72E-238 | 4 | PARD3 |
| **233** | 6.30E-242 | 1.93638468 | 0.504 | 0.045 | 1.68E-237 | 4 | GEM |
| **234** | 2.41E-240 | 2.44349649 | 0.727 | 0.115 | 6.43E-236 | 4 | RORA |
| **235** | 5.01E-240 | 1.92489844 | 0.369 | 0.02 | 1.34E-235 | 4 | DGKI |
| **236** | 6.71E-239 | 1.33434787 | 0.369 | 0.02 | 1.79E-234 | 4 | MSRB3 |
| **237** | 6.10E-237 | 1.65812521 | 0.464 | 0.037 | 1.63E-232 | 4 | HMCN1 |
| **238** | 1.94E-234 | 1.08329721 | 0.255 | 0.006 | 5.18E-230 | 4 | AC079298.3 |
| **239** | 3.43E-234 | 1.24836191 | 0.316 | 0.013 | 9.15E-230 | 4 | NXN |
| **240** | 3.80E-234 | 1.51833176 | 0.435 | 0.032 | 1.01E-229 | 4 | TEAD1 |
| **241** | 9.98E-232 | 1.61946949 | 0.456 | 0.037 | 2.66E-227 | 4 | DPYSL3 |
| **242** | 4.40E-231 | 1.16550967 | 0.297 | 0.011 | 1.17E-226 | 4 | GULP1 |
| **243** | 1.10E-230 | 1.9109292 | 0.594 | 0.069 | 2.93E-226 | 4 | KIAA1217 |
| **244** | 2.44E-229 | 1.41885739 | 0.355 | 0.019 | 6.52E-225 | 4 | PRICKLE1 |
| **245** | 1.86E-228 | 1.61736111 | 0.374 | 0.023 | 4.97E-224 | 4 | ROR2 |
| **246** | 1.24E-222 | 1.573611 | 0.454 | 0.038 | 3.32E-218 | 4 | LAMB1 |
| **247** | 2.06E-221 | 1.1059089 | 0.279 | 0.01 | 5.50E-217 | 4 | MEIS1 |
| **248** | 4.89E-219 | 1.22382902 | 0.292 | 0.012 | 1.31E-214 | 4 | FHOD3 |
| **249** | 1.40E-211 | 2.24276492 | 0.597 | 0.079 | 3.73E-207 | 4 | ADAM12 |
| **250** | 2.59E-208 | 2.1776395 | 0.517 | 0.059 | 6.92E-204 | 4 | COL8A1 |
| **251** | 3.66E-208 | 1.50825676 | 0.297 | 0.014 | 9.77E-204 | 4 | NTM |
| **252** | 4.60E-205 | 1.75959805 | 0.228 | 0.006 | 1.23E-200 | 4 | LINC00922 |
| **253** | 1.76E-202 | 1.82302477 | 0.629 | 0.094 | 4.70E-198 | 4 | APBB2 |
| **254** | 3.40E-199 | 1.17334404 | 0.22 | 0.006 | 9.07E-195 | 4 | SLC24A2 |
| **255** | 1.50E-198 | 2.5352901 | 0.512 | 0.062 | 4.00E-194 | 4 | KIF26B |
| **256** | 5.79E-197 | 1.33645604 | 0.334 | 0.021 | 1.54E-192 | 4 | DCLK1 |
| **257** | 1.31E-194 | 2.23881559 | 0.751 | 0.147 | 3.51E-190 | 4 | COL5A2 |
| **258** | 9.77E-194 | 1.14246422 | 0.31 | 0.018 | 2.61E-189 | 4 | DDAH1 |
| **259** | 2.10E-193 | 1.16044511 | 0.379 | 0.03 | 5.61E-189 | 4 | MSC-AS1 |
| **260** | 1.54E-191 | 1.42330779 | 0.35 | 0.025 | 4.11E-187 | 4 | SEMA5A |
| **261** | 1.50E-190 | 1.3352841 | 0.401 | 0.035 | 4.00E-186 | 4 | LAMA4 |
| **262** | 2.34E-188 | 1.96671278 | 0.432 | 0.044 | 6.25E-184 | 4 | ITGBL1 |
| **263** | 8.22E-187 | 1.13157693 | 0.255 | 0.011 | 2.20E-182 | 4 | CARMN |
| **264** | 5.33E-181 | 1.08568357 | 0.241 | 0.01 | 1.42E-176 | 4 | AC004160.1 |
| **265** | 1.41E-179 | 1.30334906 | 0.432 | 0.045 | 3.76E-175 | 4 | RBFOX2 |
| **266** | 4.58E-178 | 1.27820937 | 0.43 | 0.044 | 1.22E-173 | 4 | GLI3 |
| **267** | 4.79E-176 | 2.03901501 | 0.69 | 0.137 | 1.28E-171 | 4 | PALLD |
| **268** | 1.33E-174 | 1.04814674 | 0.355 | 0.029 | 3.54E-170 | 4 | SH3D19 |
| **269** | 1.82E-174 | 1.60199836 | 0.531 | 0.074 | 4.85E-170 | 4 | PARD3B |

| | | | | | | | |
|---|---|---|---|---|---|---|---|
| **270** | 1.25E-172 | 1.33685592 | 0.398 | 0.039 | 3.33E-168 | 4 | WWTR1 |
| **271** | 1.80E-170 | 1.12824157 | 0.271 | 0.015 | 4.80E-166 | 4 | TENM3 |
| **272** | 5.68E-169 | 1.38309164 | 0.411 | 0.043 | 1.52E-164 | 4 | RBPMS |
| **273** | 5.80E-169 | 1.29142534 | 0.207 | 0.007 | 1.55E-164 | 4 | AADACL2-AS1 |
| **274** | 2.93E-168 | 1.48463292 | 0.496 | 0.066 | 7.81E-164 | 4 | PPFIBP1 |
| **275** | 6.99E-168 | 1.14677089 | 0.366 | 0.033 | 1.87E-163 | 4 | CEP112 |
| **276** | 2.53E-165 | 1.09217487 | 0.233 | 0.011 | 6.76E-161 | 4 | LRIG3 |
| **277** | 2.95E-165 | 1.93302857 | 0.634 | 0.119 | 7.87E-161 | 4 | RBMS3 |
| **278** | 2.33E-161 | 1.34440108 | 0.326 | 0.027 | 6.22E-157 | 4 | MAGI2 |
| **279** | 1.06E-160 | 1.20367167 | 0.361 | 0.034 | 2.82E-156 | 4 | RNF144A |
| **280** | 1.55E-160 | 1.86779822 | 0.549 | 0.09 | 4.14E-156 | 4 | NAV3 |
| **281** | 2.50E-156 | 1.37271842 | 0.342 | 0.031 | 6.69E-152 | 4 | INHBA |
| **282** | 7.13E-156 | 1.50024665 | 0.332 | 0.03 | 1.90E-151 | 4 | ADAMTS6 |
| **283** | 5.21E-154 | 1.12571316 | 0.292 | 0.022 | 1.39E-149 | 4 | SORBS1 |
| **284** | 3.02E-152 | 1.73649356 | 0.56 | 0.094 | 8.07E-148 | 4 | THBS2 |
| **285** | 4.12E-151 | 1.21053265 | 0.289 | 0.022 | 1.10E-146 | 4 | ITGA11 |
| **286** | 4.71E-149 | 1.34821884 | 0.379 | 0.041 | 1.26E-144 | 4 | RAI14 |
| **287** | 9.40E-149 | 1.01331388 | 0.294 | 0.023 | 2.51E-144 | 4 | EGFR |
| **288** | 2.14E-146 | 1.11293652 | 0.244 | 0.015 | 5.70E-142 | 4 | DCLK2 |
| **289** | 2.32E-146 | 1.02589957 | 0.292 | 0.023 | 6.18E-142 | 4 | OSBPL10 |
| **290** | 7.27E-145 | 1.44887913 | 0.507 | 0.08 | 1.94E-140 | 4 | ENAH |
| **291** | 3.55E-143 | 1.40927285 | 0.488 | 0.075 | 9.48E-139 | 4 | SVIL |
| **292** | 3.66E-143 | 1.34008859 | 0.345 | 0.036 | 9.78E-139 | 4 | NFIB |
| **293** | 4.53E-141 | 2.01575595 | 0.544 | 0.097 | 1.21E-136 | 4 | SULF1 |
| **294** | 9.94E-141 | 1.57141005 | 0.382 | 0.046 | 2.65E-136 | 4 | NEGR1 |
| **295** | **8.24E-117** | **3.75980423** | **0.893** | **0.457** | **2.20E-112** | **5** | **APOE** |
| **296** | 5.31E-116 | 2.64078959 | 0.785 | 0.262 | 1.42E-111 | 5 | C1QC |
| **297** | 5.20E-109 | 2.2483878 | 0.981 | 0.875 | 1.39E-104 | 5 | CD74 |
| **298** | 1.09E-105 | 2.23799137 | 0.659 | 0.174 | 2.92E-101 | 5 | CD68 |
| **299** | 4.10E-98 | 2.50769685 | 0.915 | 0.574 | 1.09E-93 | 5 | HLA-DRA |
| **300** | 7.72E-97 | 2.44462867 | 0.659 | 0.193 | 2.06E-92 | 5 | C1QA |
| **301** | 1.32E-94 | 2.41216777 | 0.959 | 0.744 | 3.53E-90 | 5 | PSAP |
| **302** | 2.24E-87 | 2.26683202 | 0.911 | 0.653 | 5.98E-83 | 5 | HLA-DRB1 |
| **303** | 3.20E-81 | 2.39180149 | 0.748 | 0.309 | 8.54E-77 | 5 | GRN |
| **304** | 4.13E-80 | 2.20243137 | 0.848 | 0.515 | 1.10E-75 | 5 | HLA-DPA1 |
| **305** | 4.12E-79 | 2.0047248 | 0.893 | 0.612 | 1.10E-74 | 5 | HLA-A |
| **306** | 1.14E-76 | 2.23356844 | 0.811 | 0.391 | 3.05E-72 | 5 | IFI30 |
| **307** | 1.50E-75 | 1.93665027 | 0.867 | 0.505 | 4.01E-71 | 5 | HLA-DPB1 |
| **308** | 1.26E-72 | 2.6088178 | 0.585 | 0.187 | 3.36E-68 | 5 | LYZ |

| | | | | | | | |
|---|---|---|---|---|---|---|---|
| **309** | 8.35E-71 | 2.7701533 | 0.819 | 0.433 | 2.23E-66 | 5 | CTSD |
| **310** | 1.42E-70 | 1.79248291 | 0.911 | 0.687 | 3.80E-66 | 5 | HLA-B |
| **311** | 3.09E-69 | 2.15427075 | 0.693 | 0.28 | 8.25E-65 | 5 | CTSZ |
| **312** | 4.77E-68 | 2.33237489 | 0.593 | 0.201 | 1.27E-63 | 5 | C1QB |
| **313** | 1.25E-63 | 2.37102052 | 0.874 | 0.523 | 3.35E-59 | 5 | FTL |
| **314** | 8.57E-62 | 1.63967472 | 0.944 | 0.815 | 2.29E-57 | 5 | B2M |
| **315** | 1.13E-61 | 1.78616132 | 0.674 | 0.277 | 3.00E-57 | 5 | CD81 |
| **316** | 3.81E-61 | 1.80648032 | 0.793 | 0.444 | 1.02E-56 | 5 | HLA-DRB5 |
| **317** | 1.08E-60 | 1.69018608 | 0.881 | 0.647 | 2.87E-56 | 5 | HLA-C |
| **318** | 3.61E-58 | 1.39903059 | 0.981 | 0.919 | 9.65E-54 | 5 | MT-CO1 |
| **319** | **3.89E-58** | **2.35443345** | **0.396** | **0.099** | **1.04E-53** | **5** | **APOC1** |
| **320** | 1.21E-57 | 1.99862481 | 0.73 | 0.37 | 3.23E-53 | 5 | HLA-DQA1 |
| **321** | 2.59E-56 | 1.83152973 | 0.741 | 0.374 | 6.91E-52 | 5 | CST3 |
| **322** | 5.49E-51 | 1.89169267 | 0.585 | 0.238 | 1.47E-46 | 5 | ITGB2 |
| **323** | 8.52E-46 | 2.16493928 | 0.244 | 0.046 | 2.28E-41 | 5 | MMP9 |
| **324** | 3.47E-43 | 1.2647492 | 0.911 | 0.693 | 9.26E-39 | 5 | MT-CO2 |
| **325** | 5.91E-43 | 1.3352616 | 0.267 | 0.058 | 1.58E-38 | 5 | DNASE2 |
| **326** | 6.20E-43 | 1.49239962 | 0.867 | 0.661 | 1.65E-38 | 5 | ACTB |
| **327** | 4.11E-42 | 1.67432146 | 0.519 | 0.208 | 1.10E-37 | 5 | TYROBP |
| **328** | 1.64E-41 | 1.69316301 | 0.563 | 0.257 | 4.37E-37 | 5 | CTSS |
| **329** | 5.02E-39 | 1.49099748 | 0.467 | 0.176 | 1.34E-34 | 5 | NPC2 |
| **330** | 2.47E-38 | 1.51765581 | 0.363 | 0.112 | 6.59E-34 | 5 | SERPINA1 |
| **331** | 4.02E-38 | 1.70281326 | 0.681 | 0.394 | 1.07E-33 | 5 | TMSB4X |
| **332** | 1.59E-37 | 1.35120296 | 0.815 | 0.542 | 4.25E-33 | 5 | MT-ND5 |
| **333** | 2.93E-37 | 1.23695103 | 0.822 | 0.587 | 7.82E-33 | 5 | ITM2B |
| **334** | 2.13E-36 | 1.13427359 | 0.9 | 0.736 | 5.69E-32 | 5 | MT-CO3 |
| **335** | 3.19E-36 | 1.64931792 | 0.43 | 0.164 | 8.51E-32 | 5 | PLD3 |
| **336** | 1.68E-35 | 1.06851181 | 0.878 | 0.655 | 4.49E-31 | 5 | MT-CYB |
| **337** | 1.71E-34 | 1.55593824 | 0.659 | 0.386 | 4.56E-30 | 5 | A2M |
| **338** | 4.60E-34 | 1.39723929 | 0.674 | 0.4 | 1.23E-29 | 5 | HLA-E |
| **339** | 6.44E-34 | 1.40362746 | 0.363 | 0.119 | 1.72E-29 | 5 | TMEM176B |
| **340** | 5.68E-33 | 1.94877763 | 0.507 | 0.24 | 1.52E-28 | 5 | GPNMB |
| **341** | 6.01E-33 | 1.31120892 | 0.363 | 0.123 | 1.60E-28 | 5 | ATP6AP2 |
| **342** | 1.02E-32 | 1.18181627 | 0.563 | 0.26 | 2.72E-28 | 5 | MT-ND3 |
| **343** | 1.85E-32 | 1.4360641 | 0.596 | 0.322 | 4.94E-28 | 5 | CYBA |
| **344** | 5.62E-31 | 1.74267456 | 0.278 | 0.082 | 1.50E-26 | 5 | ACP5 |
| **345** | 7.34E-31 | 1.50212116 | 0.63 | 0.37 | 1.96E-26 | 5 | HLA-DQB1 |
| **346** | 4.29E-30 | 1.12738457 | 0.774 | 0.53 | 1.15E-25 | 5 | MT-ND4L |
| **347** | 1.07E-29 | 1.19713334 | 0.722 | 0.453 | 2.86E-25 | 5 | MT-ND1 |

| | | | | | | | |
|---|---|---|---|---|---|---|---|
| **348** | 7.46E-29 | 1.30837647 | 0.574 | 0.302 | 1.99E-24 | 5 | CD63 |
| **349** | 8.64E-29 | 1.18485559 | 0.778 | 0.548 | 2.31E-24 | 5 | MT-ATP6 |
| **350** | 1.31E-28 | 1.30390111 | 0.241 | 0.066 | 3.50E-24 | 5 | IGSF6 |
| **351** | 1.62E-28 | 1.00472967 | 0.778 | 0.557 | 4.33E-24 | 5 | MTRNR2L12 |
| **352** | 8.59E-28 | 1.19706074 | 0.622 | 0.368 | 2.29E-23 | 5 | LAPTM5 |
| **353** | 2.70E-27 | 1.17511491 | 0.244 | 0.07 | 7.21E-23 | 5 | GM2A |
| **354** | 3.74E-27 | 1.32295587 | 0.585 | 0.331 | 9.98E-23 | 5 | CALR |
| **355** | 5.49E-27 | 1.24312202 | 0.278 | 0.088 | 1.47E-22 | 5 | SH3BGRL3 |
| **356** | 7.94E-26 | 1.26896477 | 0.441 | 0.196 | 2.12E-21 | 5 | CFL1 |
| **357** | **1.49E-25** | **1.41124722** | **0.244** | **0.074** | **3.99E-21** | **5** | **TREM2** |
| **358** | 2.81E-25 | 1.29042445 | 0.444 | 0.208 | 7.49E-21 | 5 | LAMP1 |
| **359** | 3.52E-25 | 1.49521544 | 0.378 | 0.161 | 9.40E-21 | 5 | FCGR3A |
| **360** | 4.16E-25 | 1.31609986 | 0.815 | 0.653 | 1.11E-20 | 5 | CTSB |
| **361** | 6.25E-25 | 1.11069147 | 0.637 | 0.397 | 1.67E-20 | 5 | MT-ND4 |
| **362** | 2.08E-24 | 1.57485065 | 0.519 | 0.281 | 5.55E-20 | 5 | C3 |
| **363** | 2.25E-24 | 1.29737631 | 0.344 | 0.133 | 6.00E-20 | 5 | CD14 |
| **364** | **1.25E-23** | **1.67391272** | **0.322** | **0.125** | **3.34E-19** | **5** | **LIPA** |
| **365** | 1.38E-23 | 1.21648566 | 0.419 | 0.191 | 3.68E-19 | 5 | ATP6V0C |
| **366** | 1.97E-23 | 1.28282183 | 0.511 | 0.265 | 5.26E-19 | 5 | S100A11 |
| **367** | 5.78E-23 | 1.13642391 | 0.574 | 0.334 | 1.54E-18 | 5 | CANX |
| **368** | 2.02E-22 | 1.12526311 | 0.593 | 0.36 | 5.39E-18 | 5 | CYBB |
| **369** | 9.23E-22 | 1.40355363 | 0.215 | 0.066 | 2.46E-17 | 5 | FUCA1 |
| **370** | 1.85E-21 | 1.22710793 | 0.289 | 0.108 | 4.94E-17 | 5 | PRDX1 |
| **371** | 2.37E-21 | 1.19317567 | 0.167 | 0.043 | 6.33E-17 | 5 | PLD4 |
| **372** | 4.39E-21 | 1.17347772 | 0.5 | 0.271 | 1.17E-16 | 5 | MT-ATP8 |
| **373** | 1.19E-20 | 1.15339275 | 0.367 | 0.164 | 3.17E-16 | 5 | ADA2 |
| **374** | 1.93E-20 | 1.23363589 | 0.407 | 0.195 | 5.15E-16 | 5 | MT-ND6 |
| **375** | 2.06E-20 | 1.00284524 | 0.389 | 0.176 | 5.49E-16 | 5 | SYNGR2 |
| **376** | 3.63E-20 | 1.07237472 | 0.237 | 0.08 | 9.68E-16 | 5 | IL4I1 |
| **377** | 5.72E-20 | 1.07961201 | 0.389 | 0.178 | 1.53E-15 | 5 | MAN2B1 |
| **378** | 6.04E-20 | 1.0703754 | 0.53 | 0.303 | 1.61E-15 | 5 | TMBIM6 |
| **379** | 1.68E-19 | 1.10538557 | 0.4 | 0.193 | 4.48E-15 | 5 | TLN1 |
| **380** | 2.16E-19 | 1.17339543 | 0.278 | 0.108 | 5.77E-15 | 5 | MARCKS |
| **381** | 2.19E-19 | 1.20573253 | 0.456 | 0.25 | 5.85E-15 | 5 | PFN1 |
| **382** | 3.55E-19 | 1.02639773 | 0.548 | 0.321 | 9.48E-15 | 5 | RPS19 |
| **383** | 4.59E-19 | 1.34718073 | 0.333 | 0.147 | 1.22E-14 | 5 | FCER1G |
| **384** | 2.70E-18 | 1.00324935 | 0.456 | 0.242 | 7.20E-14 | 5 | PPIB |
| **385** | 3.00E-18 | 1.11917197 | 0.548 | 0.354 | 8.02E-14 | 5 | PDIA3 |
| **386** | 5.90E-18 | 1.27827899 | 0.5 | 0.304 | 1.58E-13 | 5 | CTSC |

| | | | | | | | |
|---|---|---|---|---|---|---|---|
| **387** | 7.72E-18 | 1.39039264 | 0.296 | 0.125 | 2.06E-13 | 5 | IFI27 |
| **388** | 1.02E-17 | 1.19918935 | 0.426 | 0.23 | 2.73E-13 | 5 | FCGRT |
| **389** | 5.44E-17 | 1.09134574 | 0.304 | 0.131 | 1.45E-12 | 5 | LGALS3BP |
| **390** | 6.78E-17 | 1.20715946 | 0.489 | 0.289 | 1.81E-12 | 5 | HSPA5 |
| **391** | 7.49E-17 | 1.02342354 | 0.459 | 0.25 | 2.00E-12 | 5 | FGL2 |
| **392** | 8.59E-17 | 1.00796584 | 0.411 | 0.213 | 2.29E-12 | 5 | HSPA8 |
| **393** | 2.14E-16 | 1.14764057 | 0.167 | 0.051 | 5.72E-12 | 5 | CXCL9 |
| **394** | 3.90E-16 | 1.16720429 | 0.4 | 0.212 | 1.04E-11 | 5 | CTSL |
| **395** | 1.22E-191 | 2.82934661 | 0.678 | 0.093 | 3.25E-187 | 6 | SKAP1 |
| **396** | 1.52E-163 | 2.67938687 | 0.694 | 0.118 | 4.06E-159 | 6 | ETS1 |
| **397** | 1.51E-152 | 2.37400004 | 0.506 | 0.061 | 4.04E-148 | 6 | CD96 |
| **398** | 1.96E-152 | 2.34637823 | 0.473 | 0.052 | 5.22E-148 | 6 | CAMK4 |
| **399** | 1.57E-150 | 2.72787322 | 0.502 | 0.061 | 4.19E-146 | 6 | THEMIS |
| **400** | 7.80E-142 | 2.81444277 | 0.996 | 0.493 | 2.08E-137 | 6 | COL1A2 |
| **401** | 1.37E-138 | 2.95885311 | 0.996 | 0.547 | 3.66E-134 | 6 | COL1A1 |
| **402** | 3.66E-134 | 2.56755597 | 0.727 | 0.161 | 9.77E-130 | 6 | FYN |
| **403** | 3.38E-133 | 2.68099944 | 0.98 | 0.464 | 9.03E-129 | 6 | COL3A1 |
| **404** | 1.48E-130 | 2.76540763 | 0.571 | 0.095 | 3.95E-126 | 6 | IL7R |
| **405** | 2.71E-130 | 2.55805783 | 0.776 | 0.198 | 7.24E-126 | 6 | COL6A1 |
| **406** | 4.54E-130 | 2.22315455 | 0.469 | 0.061 | 1.21E-125 | 6 | ITK |
| **407** | 7.58E-127 | 2.61006527 | 0.502 | 0.072 | 2.02E-122 | 6 | TOX |
| **408** | 1.45E-115 | 2.11159802 | 0.396 | 0.048 | 3.86E-111 | 6 | BCL11B |
| **409** | 2.87E-108 | 2.32642013 | 0.788 | 0.239 | 7.67E-104 | 6 | COL6A3 |
| **410** | 6.77E-106 | 2.19684464 | 0.38 | 0.048 | 1.81E-101 | 6 | LINC01934 |
| **411** | 1.36E-89 | 2.32857574 | 0.792 | 0.309 | 3.62E-85 | 6 | SFRP2 |
| **412** | 1.12E-88 | 1.77023763 | 0.31 | 0.038 | 3.00E-84 | 6 | PRKCQ |
| **413** | 2.78E-88 | 1.99145717 | 0.371 | 0.056 | 7.41E-84 | 6 | SLAMF6 |
| **414** | 3.15E-85 | 2.06660617 | 0.551 | 0.13 | 8.40E-81 | 6 | COL5A1 |
| **415** | 9.37E-85 | 2.02119037 | 0.971 | 0.651 | 2.50E-80 | 6 | FN1 |
| **416** | 1.43E-84 | 2.24269691 | 0.633 | 0.182 | 3.83E-80 | 6 | COL6A2 |
| **417** | 7.34E-84 | 2.06120759 | 0.298 | 0.037 | 1.96E-79 | 6 | SCML4 |
| **418** | 2.23E-83 | 2.00910908 | 0.616 | 0.165 | 5.95E-79 | 6 | COL5A2 |
| **419** | 5.37E-82 | 2.15047179 | 0.453 | 0.09 | 1.43E-77 | 6 | TC2N |
| **420** | 8.50E-81 | 1.82017449 | 0.339 | 0.05 | 2.27E-76 | 6 | CD247 |
| **421** | 1.74E-79 | 1.95712686 | 0.49 | 0.107 | 4.64E-75 | 6 | THBS2 |
| **422** | 8.15E-78 | 1.78449457 | 0.31 | 0.043 | 2.18E-73 | 6 | ZNF831 |
| **423** | 2.00E-76 | 2.03533792 | 0.829 | 0.403 | 5.33E-72 | 6 | SPARC |
| **424** | 2.50E-72 | 1.52896161 | 0.22 | 0.023 | 6.68E-68 | 6 | TESPA1 |
| **425** | 2.46E-71 | 2.06652498 | 0.437 | 0.096 | 6.56E-67 | 6 | STAT4 |

| | | | | | | | |
|---|---|---|---|---|---|---|---|
| **426** | 3.13E-71 | 2.00162381 | 0.567 | 0.161 | 8.35E-67 | 6 | AEBP1 |
| **427** | 1.45E-68 | 1.90336782 | 0.478 | 0.118 | 3.88E-64 | 6 | SYNE2 |
| **428** | 1.71E-67 | 1.87907429 | 0.461 | 0.11 | 4.57E-63 | 6 | RIPOR2 |
| **429** | 1.08E-66 | 2.14291155 | 0.763 | 0.355 | 2.88E-62 | 6 | POSTN |
| **430** | 1.37E-66 | 1.93295589 | 0.543 | 0.158 | 3.66E-62 | 6 | INPP4B |
| **431** | 2.00E-65 | 1.85406239 | 0.637 | 0.221 | 5.34E-61 | 6 | VCAN |
| **432** | 2.51E-65 | 1.88681809 | 0.473 | 0.119 | 6.70E-61 | 6 | FBN1 |
| **433** | 3.31E-65 | 1.80909584 | 0.306 | 0.05 | 8.84E-61 | 6 | SLFN12L |
| **434** | 2.56E-64 | 1.49963268 | 0.249 | 0.033 | 6.83E-60 | 6 | CD3E |
| **435** | 5.24E-64 | 1.71439037 | 0.4 | 0.085 | 1.40E-59 | 6 | MXRA5 |
| **436** | 1.26E-60 | 1.85081585 | 0.531 | 0.16 | 3.38E-56 | 6 | MMP2 |
| **437** | 9.02E-60 | 1.5778626 | 0.229 | 0.03 | 2.41E-55 | 6 | GRAP2 |
| **438** | 1.93E-59 | 2.09007109 | 0.261 | 0.04 | 5.16E-55 | 6 | IFNG-AS1 |
| **439** | 8.38E-59 | 1.38084229 | 0.22 | 0.028 | 2.24E-54 | 6 | ZAP70 |
| **440** | 4.53E-58 | 1.49821309 | 0.253 | 0.038 | 1.21E-53 | 6 | PYHIN1 |
| **441** | 2.81E-57 | 1.79933257 | 0.314 | 0.06 | 7.49E-53 | 6 | PPP1R16B |
| **442** | 3.29E-57 | 2.08944814 | 0.51 | 0.158 | 8.79E-53 | 6 | ANK3 |
| **443** | 5.71E-57 | 1.74323035 | 0.58 | 0.194 | 1.52E-52 | 6 | COL12A1 |
| **444** | 6.53E-56 | 1.51853447 | 0.241 | 0.036 | 1.74E-51 | 6 | SAMD3 |
| **445** | 5.05E-53 | 1.63512532 | 0.245 | 0.039 | 1.35E-48 | 6 | CD2 |
| **446** | 1.32E-51 | 1.67660277 | 0.506 | 0.164 | 3.51E-47 | 6 | BGN |
| **447** | 1.75E-50 | 1.44950132 | 0.22 | 0.033 | 4.68E-46 | 6 | FBLN2 |
| **448** | 4.12E-50 | 1.84860597 | 0.4 | 0.108 | 1.10E-45 | 6 | MLLT3 |
| **449** | 4.71E-49 | 1.36842919 | 0.216 | 0.032 | 1.26E-44 | 6 | RASGRP1 |
| **450** | 4.85E-49 | 1.50488804 | 0.257 | 0.046 | 1.30E-44 | 6 | IKZF3 |
| **451** | 1.16E-48 | 1.43499111 | 0.184 | 0.024 | 3.11E-44 | 6 | AC006369.1 |
| **452** | 9.25E-48 | 1.42007941 | 0.229 | 0.037 | 2.47E-43 | 6 | LINC00861 |
| **453** | 1.04E-47 | 1.7864388 | 0.527 | 0.192 | 2.78E-43 | 6 | NCK2 |
| **454** | 3.38E-47 | 1.54725125 | 0.543 | 0.197 | 9.03E-43 | 6 | LUM |
| **455** | 4.46E-46 | 1.64734488 | 0.286 | 0.06 | 1.19E-41 | 6 | CCDC88C |
| **456** | 2.39E-45 | 1.58682238 | 0.188 | 0.026 | 6.37E-41 | 6 | DTHD1 |
| **457** | 2.82E-45 | 1.68365816 | 0.261 | 0.05 | 7.52E-41 | 6 | NELL2 |
| **458** | 7.90E-45 | 1.49828805 | 0.69 | 0.347 | 2.11E-40 | 6 | PRKCH |
| **459** | 8.81E-45 | 1.7174521 | 0.702 | 0.4 | 2.35E-40 | 6 | PARP8 |
| **460** | 4.40E-44 | 1.51795266 | 0.294 | 0.065 | 1.18E-39 | 6 | FBLN1 |
| **461** | 1.40E-43 | 1.34795142 | 0.192 | 0.029 | 3.73E-39 | 6 | SPOCK2 |
| **462** | 3.93E-42 | 1.54509437 | 0.184 | 0.027 | 1.05E-37 | 6 | KLRK1 |
| **463** | 9.95E-42 | 1.3797721 | 0.163 | 0.022 | 2.66E-37 | 6 | ICOS |
| **464** | 3.52E-41 | 1.75393064 | 0.314 | 0.078 | 9.41E-37 | 6 | SFRP4 |

| | | | | | | | |
|---|---|---|---|---|---|---|---|
| **465** | 3.77E-41 | 1.6228875 | 0.514 | 0.204 | 1.01E-36 | 6 | CASK |
| **466** | 6.61E-41 | 1.86500091 | 0.424 | 0.143 | 1.77E-36 | 6 | CDC14A |
| **467** | 8.16E-41 | 1.49176905 | 0.257 | 0.053 | 2.18E-36 | 6 | CILP |
| **468** | 1.24E-39 | 1.66014826 | 0.555 | 0.249 | 3.31E-35 | 6 | RNF19A |
| **469** | 1.48E-39 | 1.4898587 | 0.457 | 0.16 | 3.95E-35 | 6 | SLC38A1 |
| **470** | 2.45E-39 | 1.78499023 | 0.461 | 0.172 | 6.55E-35 | 6 | PITPNC1 |
| **471** | 3.06E-39 | 1.57834896 | 0.343 | 0.096 | 8.16E-35 | 6 | C1S |
| **472** | 2.07E-38 | 1.49475463 | 0.184 | 0.03 | 5.52E-34 | 6 | PPP2R2B |
| **473** | 3.97E-38 | 1.59122573 | 0.404 | 0.135 | 1.06E-33 | 6 | KLF12 |
| **474** | 4.03E-38 | 1.48977169 | 0.241 | 0.05 | 1.08E-33 | 6 | FAM102A |
| **475** | 4.43E-38 | 1.29772942 | 0.188 | 0.031 | 1.18E-33 | 6 | CD6 |
| **476** | 5.02E-38 | 1.61481648 | 0.506 | 0.198 | 1.34E-33 | 6 | CCN2 |
| **477** | 5.51E-38 | 1.46290706 | 0.233 | 0.047 | 1.47E-33 | 6 | LEF1 |
| **478** | 1.05E-37 | 1.02734342 | 0.127 | 0.014 | 2.79E-33 | 6 | TRAT1 |
| **479** | 1.61E-36 | 1.65941793 | 0.49 | 0.202 | 4.30E-32 | 6 | TNFAIP8 |
| **480** | 2.12E-36 | 1.62910895 | 0.318 | 0.089 | 5.65E-32 | 6 | OXNAD1 |
| **481** | 4.05E-36 | 1.62409169 | 0.498 | 0.203 | 1.08E-31 | 6 | DCN |
| **482** | 3.13E-35 | 1.58769425 | 0.478 | 0.197 | 8.37E-31 | 6 | CNOT6L |
| **483** | 1.69E-34 | 1.14380571 | 0.151 | 0.022 | 4.51E-30 | 6 | SLAMF1 |
| **484** | 1.82E-34 | 1.58388612 | 0.367 | 0.117 | 4.85E-30 | 6 | CD69 |
| **485** | 3.28E-34 | 1.04884803 | 0.151 | 0.023 | 8.76E-30 | 6 | SEPTIN1 |
| **486** | 2.03E-33 | 1.44543602 | 0.327 | 0.097 | 5.43E-29 | 6 | TNIK |
| **487** | 1.48E-32 | 1.20555703 | 0.2 | 0.04 | 3.95E-28 | 6 | CYFIP2 |
| **488** | 7.46E-32 | 1.37807691 | 0.224 | 0.051 | 1.99E-27 | 6 | PCAT1 |
| **489** | 1.05E-31 | 1.52148415 | 0.4 | 0.141 | 2.80E-27 | 6 | RORA |
| **490** | 1.11E-31 | 1.76890445 | 0.441 | 0.178 | 2.97E-27 | 6 | NIBAN1 |
| **491** | 1.30E-31 | 1.43628131 | 0.249 | 0.062 | 3.48E-27 | 6 | TSPAN5 |
| **492** | 1.45E-31 | 1.42453832 | 0.624 | 0.356 | 3.86E-27 | 6 | CBLB |
| **493** | 1.88E-31 | 1.19203631 | 0.212 | 0.047 | 5.02E-27 | 6 | ACAP1 |
| **494** | 3.68E-31 | 1.3865753 | 0.347 | 0.116 | 9.82E-27 | 6 | STK17A |
| **495** | 1.13E-235 | 1.66159053 | 0.299 | 0.005 | 3.00E-231 | 7 | SPOCD1 |
| **496** | 3.31E-233 | 2.45480922 | 0.372 | 0.011 | 8.84E-229 | 7 | TM4SF19 |
| **497** | 2.36E-206 | 2.58842055 | 0.36 | 0.012 | 6.29E-202 | 7 | FABP4 |
| **498** | 1.41E-153 | 1.31583343 | 0.159 | 0.002 | 3.77E-149 | 7 | ACTN2 |
| **499** | 1.56E-153 | 4.19599177 | 0.683 | 0.086 | 4.16E-149 | 7 | CD36 |
| **500** | 9.76E-149 | 1.52251052 | 0.335 | 0.017 | 2.61E-144 | 7 | ZMIZ1-AS1 |
| **501** | 9.51E-132 | 2.40282356 | 0.451 | 0.039 | 2.54E-127 | 7 | LPL |
| **502** | 8.29E-126 | 2.2252671 | 0.396 | 0.031 | 2.21E-121 | 7 | DOCK3 |
| **503** | 1.35E-114 | 1.29066793 | 0.165 | 0.004 | 3.61E-110 | 7 | ATP6V0D2 |

| | | | | | | | |
|---|---|---|---|---|---|---|---|
| **504** | 6.52E-112 | 1.48540734 | 0.354 | 0.026 | 1.74E-107 | 7 | AQP9 |
| **505** | 2.26E-110 | 1.9959885 | 0.409 | 0.037 | 6.02E-106 | 7 | MMP19 |
| **506** | 3.96E-107 | 1.87327405 | 0.463 | 0.05 | 1.06E-102 | 7 | FABP5 |
| **507** | 2.46E-102 | 1.35567046 | 0.128 | 0.002 | 6.57E-98 | 7 | LINC02725 |
| **508** | 2.86E-97 | 2.92590648 | 0.329 | 0.027 | 7.63E-93 | 7 | CHI3L1 |
| **509** | 1.24E-95 | 2.06580573 | 0.433 | 0.049 | 3.31E-91 | 7 | SLC39A8 |
| **510** | 4.59E-81 | 1.80230543 | 0.494 | 0.077 | 1.23E-76 | 7 | FAM20C |
| **511** | 4.18E-77 | 1.75404183 | 0.36 | 0.041 | 1.12E-72 | 7 | ANPEP |
| **512** | 1.50E-76 | 2.72936727 | 0.256 | 0.021 | 4.00E-72 | 7 | ZNF385D |
| **513** | 1.76E-69 | 2.38064968 | 0.683 | 0.166 | 4.71E-65 | 7 | SPP1 |
| **514** | 1.98E-67 | 1.11716355 | 0.256 | 0.023 | 5.28E-63 | 7 | AC023282.1 |
| **515** | 1.73E-65 | 1.95493129 | 0.457 | 0.078 | 4.61E-61 | 7 | CD109 |
| **516** | 3.90E-62 | 2.63304728 | 0.177 | 0.012 | 1.04E-57 | 7 | SLC9B2 |
| **517** | 5.48E-60 | 1.72683079 | 0.171 | 0.011 | 1.46E-55 | 7 | AK5 |
| **518** | 8.78E-59 | 1.13102404 | 0.366 | 0.055 | 2.35E-54 | 7 | ME3 |
| **519** | 1.86E-57 | 1.24852473 | 0.384 | 0.061 | 4.95E-53 | 7 | MREG |
| **520** | 1.57E-56 | 1.71560384 | 0.287 | 0.036 | 4.19E-52 | 7 | ANO5 |
| **521** | 1.61E-56 | 1.73221114 | 0.518 | 0.114 | 4.31E-52 | 7 | MGLL |
| **522** | 1.06E-55 | 4.09133789 | 0.323 | 0.048 | 2.84E-51 | 7 | MMP9 |
| **523** | 8.00E-54 | 1.79907852 | 0.89 | 0.391 | 2.14E-49 | 7 | MITF |
| **524** | 1.35E-52 | 1.32317314 | 0.372 | 0.062 | 3.61E-48 | 7 | CYP27A1 |
| **525** | 5.53E-50 | 1.45024079 | 0.53 | 0.127 | 1.48E-45 | 7 | PLIN2 |
| **526** | 6.79E-50 | 1.02145543 | 0.22 | 0.024 | 1.81E-45 | 7 | ZNF462 |
| **527** | 1.18E-48 | 1.14670754 | 0.177 | 0.016 | 3.14E-44 | 7 | IL1RN |
| **528** | 1.57E-48 | 2.46569845 | 0.677 | 0.249 | 4.20E-44 | 7 | TPRG1 |
| **529** | 4.02E-48 | 1.30382192 | 0.348 | 0.059 | 1.07E-43 | 7 | PHLDA1 |
| **530** | 1.87E-46 | 1.98206421 | 0.238 | 0.03 | 4.99E-42 | 7 | ITGB3 |
| **531** | 7.81E-45 | 1.58779317 | 0.787 | 0.306 | 2.08E-40 | 7 | ARHGAP10 |
| **532** | 2.45E-43 | 1.4913874 | 0.28 | 0.044 | 6.54E-39 | 7 | MSC-AS1 |
| **533** | 8.69E-43 | 1.68634047 | 0.75 | 0.318 | 2.32E-38 | 7 | CCDC88A |
| **534** | 2.20E-41 | 1.84782937 | 0.433 | 0.104 | 5.87E-37 | 7 | SDC2 |
| **535** | 5.09E-40 | 1.50426532 | 0.53 | 0.158 | 1.36E-35 | 7 | NCEH1 |
| **536** | 1.75E-38 | 1.45930731 | 0.213 | 0.029 | 4.68E-34 | 7 | JAKMIP2 |
| **537** | 6.71E-38 | 1.41243727 | 0.134 | 0.012 | 1.79E-33 | 7 | LINC00511 |
| **538** | 9.31E-37 | 1.34740506 | 0.183 | 0.022 | 2.49E-32 | 7 | ALDH1A2 |
| **539** | 4.87E-36 | 1.66235321 | 0.75 | 0.344 | 1.30E-31 | 7 | MYO1E |
| **540** | 1.78E-35 | 1.46665469 | 0.36 | 0.082 | 4.76E-31 | 7 | RGCC |
| **541** | 8.96E-35 | 1.29244897 | 0.451 | 0.125 | 2.39E-30 | 7 | LIPA |
| **542** | 1.57E-34 | 2.81714515 | 0.354 | 0.083 | 4.20E-30 | 7 | ACP5 |

| | | | | | | | |
|---|---|---|---|---|---|---|---|
| **543** | 2.80E-34 | 1.36239761 | 0.14 | 0.014 | 7.46E-30 | 7 | SCD5 |
| **544** | 3.00E-34 | 1.51704301 | 0.604 | 0.214 | 8.01E-30 | 7 | TCIRG1 |
| **545** | 4.55E-34 | 1.32745009 | 0.366 | 0.087 | 1.21E-29 | 7 | TNFRSF11A |
| **546** | 1.44E-33 | 1.07452919 | 0.457 | 0.127 | 3.83E-29 | 7 | PAPSS1 |
| **547** | 4.71E-32 | 1.20680403 | 0.421 | 0.115 | 1.26E-27 | 7 | ST18 |
| **548** | 3.95E-31 | 1.58994132 | 0.415 | 0.12 | 1.05E-26 | 7 | NR1H3 |
| **549** | 7.15E-31 | 1.41707108 | 0.524 | 0.177 | 1.91E-26 | 7 | PPARG |
| **550** | 5.13E-30 | 1.24218964 | 0.476 | 0.152 | 1.37E-25 | 7 | SEMA3C |
| **551** | 1.51E-29 | 1.03689474 | 0.195 | 0.031 | 4.02E-25 | 7 | NOS1AP |
| **552** | 2.13E-29 | 1.50610429 | 0.537 | 0.182 | 5.68E-25 | 7 | SNTB1 |
| **553** | 4.18E-28 | 1.89553043 | 0.787 | 0.519 | 1.12E-23 | 7 | ALCAM |
| **554** | 5.02E-28 | 1.01903984 | 0.22 | 0.04 | 1.34E-23 | 7 | TPST1 |
| **555** | 1.38E-27 | 1.05376062 | 0.262 | 0.056 | 3.69E-23 | 7 | RAI14 |
| **556** | 5.62E-27 | 1.02865143 | 0.591 | 0.234 | 1.50E-22 | 7 | ATP13A3 |
| **557** | 6.11E-27 | 1.36300819 | 0.72 | 0.357 | 1.63E-22 | 7 | CBLB |
| **558** | 2.49E-26 | 1.23800208 | 0.378 | 0.109 | 6.66E-22 | 7 | PLA2G7 |
| **559** | 7.67E-26 | 2.06519391 | 0.195 | 0.036 | 2.05E-21 | 7 | AKAP6 |
| **560** | 1.14E-25 | 1.43145162 | 0.165 | 0.026 | 3.05E-21 | 7 | GPC4 |
| **561** | 4.38E-25 | 1.3029323 | 0.817 | 0.48 | 1.17E-20 | 7 | ASAP1 |
| **562** | 6.34E-25 | 1.07015922 | 0.451 | 0.151 | 1.69E-20 | 7 | ANKRD28 |
| **563** | 6.44E-25 | 1.04766378 | 0.488 | 0.173 | 1.72E-20 | 7 | WDR11 |
| **564** | 1.36E-23 | 1.23466085 | 0.463 | 0.167 | 3.62E-19 | 7 | SLC11A1 |
| **565** | 1.44E-23 | 1.02041727 | 0.366 | 0.11 | 3.83E-19 | 7 | CSTB |
| **566** | 5.89E-23 | 1.0056561 | 0.604 | 0.261 | 1.57E-18 | 7 | PDE8A |
| **567** | 4.46E-22 | 1.23656679 | 0.372 | 0.121 | 1.19E-17 | 7 | THRB |
| **568** | 2.62E-21 | 1.02292039 | 0.22 | 0.05 | 6.99E-17 | 7 | PLPP3 |
| **569** | 2.75E-20 | 1.19391005 | 0.646 | 0.332 | 7.34E-16 | 7 | FNIP2 |
| **570** | 3.60E-20 | 1.28752909 | 0.634 | 0.349 | 9.61E-16 | 7 | NUMB |
| **571** | 1.91E-19 | 1.00805883 | 0.713 | 0.392 | 5.11E-15 | 7 | RASAL2 |
| **572** | 2.40E-19 | 1.03544912 | 0.61 | 0.299 | 6.42E-15 | 7 | NRP1 |
| **573** | 1.37E-18 | 1.23194853 | 0.39 | 0.147 | 3.66E-14 | 7 | TDRD3 |
| **574** | 1.55E-17 | 1.00190083 | 0.427 | 0.171 | 4.13E-13 | 7 | MYO1D |
| **575** | 4.70E-17 | 1.32350854 | 0.445 | 0.186 | 1.25E-12 | 7 | GBE1 |
| **576** | 1.00E-16 | 1.00989266 | 0.598 | 0.309 | 2.68E-12 | 7 | DENND4C |
| **577** | 6.31E-16 | 1.10235706 | 0.183 | 0.046 | 1.69E-11 | 7 | MIR222HG |
| **578** | 8.94E-16 | 1.08231283 | 0.573 | 0.295 | 2.39E-11 | 7 | ZNF804A |
| **579** | 2.75E-15 | 1.11591075 | 0.396 | 0.161 | 7.34E-11 | 7 | BICD1 |
| **580** | 6.16E-15 | 1.30567114 | 0.122 | 0.024 | 1.65E-10 | 7 | ITGA2 |
| **581** | 1.33E-14 | 1.01181117 | 0.146 | 0.033 | 3.56E-10 | 7 | GLDN |

| | | | | | | | |
|---|---|---|---|---|---|---|---|
| **582** | 3.58E-14 | 1.21336757 | 0.152 | 0.037 | 9.56E-10 | 7 | RUFY4 |
| **583** | 2.02E-11 | 1.09806315 | 0.22 | 0.078 | 5.39E-07 | 7 | NFATC1 |
| **584** | 6.05E-09 | 2.01327756 | 0.207 | 0.084 | 0.00016167 | 7 | CTSK |
| **585** | 2.61E-08 | 1.03755717 | 0.378 | 0.205 | 0.00069805 | 7 | MAP4K4 |
| **586** | 0.00605815 | 1.85796251 | 0.28 | 0.21 | 1 | 7 | EXT1 |
| **587** | 1.18E-102 | 2.65509106 | 0.874 | 0.205 | 3.14E-98 | 8 | COL6A1 |
| **588** | 4.36E-86 | 2.94590462 | 1 | 0.554 | 1.16E-81 | 8 | COL1A1 |
| **589** | 3.36E-85 | 2.66976679 | 1 | 0.501 | 8.96E-81 | 8 | COL1A2 |
| **590** | 9.49E-83 | 2.64691784 | 0.979 | 0.473 | 2.53E-78 | 8 | COL3A1 |
| **591** | 7.64E-74 | 2.21050611 | 0.65 | 0.135 | 2.04E-69 | 8 | COL5A1 |
| **592** | 1.38E-73 | 1.97984082 | 0.734 | 0.17 | 3.69E-69 | 8 | COL5A2 |
| **593** | 2.78E-69 | 2.20151875 | 0.811 | 0.248 | 7.42E-65 | 8 | COL6A3 |
| **594** | 1.05E-68 | 2.24155358 | 0.72 | 0.188 | 2.80E-64 | 8 | COL6A2 |
| **595** | 8.60E-65 | 1.7855387 | 0.58 | 0.111 | 2.30E-60 | 8 | THBS2 |
| **596** | 2.44E-63 | 1.74055056 | 0.503 | 0.088 | 6.51E-59 | 8 | MXRA5 |
| **597** | 1.44E-62 | 2.02958287 | 0.783 | 0.224 | 3.86E-58 | 8 | VCAN |
| **598** | 4.74E-58 | 2.0766537 | 0.881 | 0.359 | 1.27E-53 | 8 | POSTN |
| **599** | 3.50E-55 | 1.91970274 | 0.979 | 0.656 | 9.33E-51 | 8 | FN1 |
| **600** | 9.70E-53 | 1.85717013 | 0.692 | 0.198 | 2.59E-48 | 8 | COL12A1 |
| **601** | 2.83E-51 | 2.0756653 | 0.804 | 0.316 | 7.55E-47 | 8 | SFRP2 |
| **602** | 3.01E-51 | 1.98850941 | 0.622 | 0.167 | 8.04E-47 | 8 | AEBP1 |
| **603** | 1.25E-46 | 1.68878792 | 0.524 | 0.124 | 3.35E-42 | 8 | FBN1 |
| **604** | 6.87E-46 | 1.99928095 | 0.86 | 0.41 | 1.84E-41 | 8 | SPARC |
| **605** | 2.74E-43 | 1.29732358 | 0.231 | 0.027 | 7.32E-39 | 8 | LRRC15 |
| **606** | 7.83E-42 | 1.84438738 | 0.273 | 0.039 | 2.09E-37 | 8 | COMP |
| **607** | 1.29E-40 | 1.73938265 | 0.629 | 0.2 | 3.44E-36 | 8 | LUM |
| **608** | 6.89E-38 | 1.0053396 | 0.217 | 0.027 | 1.84E-33 | 8 | EMILIN1 |
| **609** | 1.02E-37 | 1.80948608 | 0.545 | 0.166 | 2.73E-33 | 8 | MMP2 |
| **610** | 3.66E-36 | 1.48474217 | 0.343 | 0.067 | 9.77E-32 | 8 | FBLN1 |
| **611** | 1.41E-35 | 1.57260135 | 0.413 | 0.099 | 3.77E-31 | 8 | C1S |
| **612** | 2.02E-35 | 1.41250921 | 0.378 | 0.08 | 5.40E-31 | 8 | COL8A1 |
| **613** | 6.04E-31 | 1.25551378 | 0.343 | 0.076 | 1.61E-26 | 8 | COL18A1 |
| **614** | 8.03E-31 | 1.5152745 | 0.58 | 0.206 | 2.14E-26 | 8 | DCN |
| **615** | 5.13E-30 | 1.42517169 | 0.517 | 0.169 | 1.37E-25 | 8 | BGN |
| **616** | 6.21E-28 | 1.61278783 | 0.559 | 0.202 | 1.66E-23 | 8 | CCN2 |
| **617** | 1.67E-27 | 1.11376802 | 0.301 | 0.066 | 4.45E-23 | 8 | HSPG2 |
| **618** | 4.74E-27 | 1.29626532 | 0.573 | 0.208 | 1.27E-22 | 8 | HTRA1 |
| **619** | 9.11E-27 | 1.22601036 | 0.315 | 0.074 | 2.43E-22 | 8 | COL14A1 |
| **620** | 1.42E-26 | 1.2035898 | 0.315 | 0.073 | 3.79E-22 | 8 | ASPN |

| | | | | | | | |
|---|---|---|---|---|---|---|---|
| **621** | 3.90E-25 | 1.40133188 | 0.497 | 0.168 | 1.04E-20 | 8 | THBS1 |
| **622** | 7.95E-25 | 1.04943422 | 0.224 | 0.042 | 2.12E-20 | 8 | COL16A1 |
| **623** | 1.88E-24 | 1.23019291 | 0.357 | 0.096 | 5.03E-20 | 8 | CTHRC1 |
| **624** | 1.20E-23 | 1.59008671 | 0.301 | 0.075 | 3.20E-19 | 8 | MMP11 |
| **625** | 6.75E-23 | 1.56716024 | 0.385 | 0.118 | 1.80E-18 | 8 | SULF1 |
| **626** | 8.79E-23 | 1.09506756 | 0.315 | 0.082 | 2.35E-18 | 8 | CTSK |
| **627** | 3.47E-22 | 1.46432858 | 0.497 | 0.189 | 9.27E-18 | 8 | TIMP3 |
| **628** | 2.77E-21 | 1.23097251 | 0.245 | 0.057 | 7.39E-17 | 8 | CILP |
| **629** | 1.54E-20 | 1.32956124 | 0.301 | 0.082 | 4.10E-16 | 8 | SFRP4 |
| **630** | 1.61E-20 | 1.51172756 | 0.594 | 0.274 | 4.30E-16 | 8 | IGFBP7 |
| **631** | 8.10E-20 | 1.27567744 | 0.322 | 0.096 | 2.16E-15 | 8 | FSTL1 |
| **632** | 1.71E-18 | 1.30265649 | 0.357 | 0.119 | 4.58E-14 | 8 | SPARCL1 |
| **633** | 4.15E-17 | 1.12514586 | 0.28 | 0.081 | 1.11E-12 | 8 | CCDC80 |
| **634** | 1.86E-15 | 1.04479243 | 0.231 | 0.063 | 4.96E-11 | 8 | ITGBL1 |
| **635** | 5.52E-14 | 1.09748895 | 0.308 | 0.111 | 1.48E-09 | 8 | SERPINF1 |
| **636** | 1.28E-13 | 1.14479729 | 0.224 | 0.066 | 3.41E-09 | 8 | TNC |
| **637** | 4.29E-12 | 1.07480983 | 0.294 | 0.113 | 1.15E-07 | 8 | MMP14 |
| **638** | 1.87E-11 | 1.01136677 | 0.336 | 0.14 | 4.99E-07 | 8 | CCN1 |
| **639** | 3.79E-11 | 1.0037764 | 0.182 | 0.054 | 1.01E-06 | 8 | ABI3BP |
| **640** | 6.75E-11 | 1.2183692 | 0.231 | 0.08 | 1.80E-06 | 8 | COL4A1 |
| **641** | 0 | 4.74652653 | 0.946 | 0.05 | 0 | 9 | CCL4 |
| **642** | 0 | 3.87569119 | 0.853 | 0.044 | 0 | 9 | CCL3 |
| **643** | 4.78E-281 | 3.76209285 | 0.752 | 0.041 | 1.28E-276 | 9 | CCL4L2 |
| **644** | 1.80E-277 | 3.34162963 | 0.643 | 0.028 | 4.80E-273 | 9 | CCL3L1 |
| **645** | 1.18E-160 | 1.82333022 | 0.395 | 0.018 | 3.14E-156 | 9 | EGR2 |
| **646** | 1.00E-108 | 1.7627277 | 0.14 | 0.002 | 2.68E-104 | 9 | TNFSF18 |
| **647** | 1.14E-96 | 1.61759963 | 0.295 | 0.017 | 3.04E-92 | 9 | EGR3 |
| **648** | 2.07E-77 | 2.21870415 | 0.628 | 0.109 | 5.53E-73 | 9 | EGR1 |
| **649** | 1.86E-75 | 2.11188867 | 0.659 | 0.121 | 4.97E-71 | 9 | NR4A1 |
| **650** | 5.14E-72 | 2.3465366 | 0.853 | 0.241 | 1.37E-67 | 9 | CD83 |
| **651** | 6.61E-69 | 1.30549479 | 0.163 | 0.007 | 1.77E-64 | 9 | TNF |
| **652** | 1.41E-52 | 1.17144487 | 0.132 | 0.006 | 3.76E-48 | 9 | CH25H |
| **653** | 6.59E-50 | 1.35947094 | 0.209 | 0.018 | 1.76E-45 | 9 | OTUD1 |
| **654** | 1.16E-44 | 1.9150039 | 0.783 | 0.286 | 3.11E-40 | 9 | RASGEF1B |
| **655** | 1.17E-37 | 1.59874077 | 0.372 | 0.071 | 3.13E-33 | 9 | FOSB |
| **656** | 2.68E-37 | 1.8636923 | 0.605 | 0.184 | 7.15E-33 | 9 | ATF3 |
| **657** | 7.92E-35 | 1.82713073 | 0.729 | 0.285 | 2.11E-30 | 9 | BTG2 |
| **658** | 2.13E-33 | 1.61079634 | 0.481 | 0.127 | 5.69E-29 | 9 | NR4A2 |
| **659** | 2.55E-33 | 1.54761014 | 0.899 | 0.538 | 6.81E-29 | 9 | FOS |

| | | | | | | | |
|---|---|---|---|---|---|---|---|
| **660** | 1.40E-28 | 1.81369542 | 0.481 | 0.14 | 3.74E-24 | 9 | NR4A3 |
| **661** | 2.71E-28 | 1.42904175 | 0.496 | 0.149 | 7.24E-24 | 9 | C3AR1 |
| **662** | 3.20E-27 | 1.3968813 | 0.333 | 0.076 | 8.54E-23 | 9 | CXorf21 |
| **663** | 4.11E-27 | 1.67492704 | 0.202 | 0.03 | 1.10E-22 | 9 | CCL2 |
| **664** | 2.52E-26 | 1.41511435 | 0.465 | 0.141 | 6.73E-22 | 9 | GBP2 |
| **665** | 8.66E-26 | 1.20248195 | 0.465 | 0.139 | 2.31E-21 | 9 | IER2 |
| **666** | 7.71E-25 | 1.53428698 | 0.442 | 0.133 | 2.06E-20 | 9 | IER3 |
| **667** | 8.03E-25 | 1.08791618 | 0.14 | 0.016 | 2.15E-20 | 9 | IL1B |
| **668** | 1.09E-24 | 1.22134915 | 0.38 | 0.102 | 2.90E-20 | 9 | NFKBID |
| **669** | 1.57E-24 | 1.52713212 | 0.488 | 0.164 | 4.18E-20 | 9 | GADD45B |
| **670** | 1.67E-24 | 1.20291343 | 0.907 | 0.616 | 4.45E-20 | 9 | ZFP36 |
| **671** | 5.83E-24 | 1.29891935 | 0.76 | 0.388 | 1.56E-19 | 9 | KLF6 |
| **672** | 3.96E-22 | 1.19711677 | 0.442 | 0.135 | 1.06E-17 | 9 | CD14 |
| **673** | 1.69E-21 | 1.15522615 | 0.837 | 0.492 | 4.50E-17 | 9 | DUSP1 |
| **674** | 2.32E-21 | 1.01070618 | 0.124 | 0.015 | 6.19E-17 | 9 | PTGS2 |
| **675** | 6.00E-21 | 1.20314078 | 0.721 | 0.359 | 1.60E-16 | 9 | SPRED1 |
| **676** | 1.29E-20 | 1.1643231 | 0.791 | 0.451 | 3.45E-16 | 9 | SRGN |
| **677** | 1.53E-20 | 2.11605986 | 0.55 | 0.248 | 4.09E-16 | 9 | PDK4 |
| **678** | 1.90E-20 | 1.5410545 | 0.434 | 0.153 | 5.06E-16 | 9 | AC020916.1 |
| **679** | 2.28E-20 | 1.02806576 | 0.682 | 0.311 | 6.08E-16 | 9 | CLEC7A |
| **680** | 1.76E-19 | 1.12281525 | 0.775 | 0.44 | 4.69E-15 | 9 | SGK1 |
| **681** | 4.44E-19 | 1.21346568 | 0.674 | 0.326 | 1.18E-14 | 9 | PEAK1 |
| **682** | 2.89E-18 | 1.61733504 | 0.535 | 0.231 | 7.71E-14 | 9 | ANKH |
| **683** | 4.85E-17 | 1.02080495 | 0.341 | 0.108 | 1.29E-12 | 9 | SOCS6 |
| **684** | 1.78E-16 | 1.02685679 | 0.636 | 0.312 | 4.75E-12 | 9 | FCGR2A |
| **685** | 6.04E-16 | 1.21002861 | 0.457 | 0.185 | 1.61E-11 | 9 | NFKBIZ |
| **686** | 1.22E-15 | 1.4324923 | 0.302 | 0.094 | 3.27E-11 | 9 | AC100849.1 |
| **687** | 1.53E-15 | 1.14459461 | 0.357 | 0.122 | 4.10E-11 | 9 | MB21D2 |
| **688** | 4.42E-15 | 1.23722916 | 0.434 | 0.173 | 1.18E-10 | 9 | LMNA |
| **689** | 4.68E-15 | 1.2458292 | 0.45 | 0.181 | 1.25E-10 | 9 | PPARG |
| **690** | 1.86E-14 | 1.15923023 | 0.434 | 0.177 | 4.97E-10 | 9 | TSC22D2 |
| **691** | 7.30E-14 | 1.06041951 | 0.411 | 0.167 | 1.95E-09 | 9 | FAM20A |
| **692** | 8.59E-12 | 1.6766105 | 0.403 | 0.181 | 2.29E-07 | 9 | PLEK |
| **693** | 1.17E-09 | 1.06937038 | 0.287 | 0.115 | 3.14E-05 | 9 | PAPSS2 |
| **694** | 6.16E-09 | 1.41234246 | 0.271 | 0.112 | 0.00016453 | 9 | RCAN1 |
| **695** | 6.80E-09 | 1.25942308 | 0.178 | 0.058 | 0.00018165 | 9 | FGF13 |
| **696** | 1.91E-08 | 1.05745675 | 0.496 | 0.293 | 0.00051073 | 9 | NFKB1 |
| **697** | 1.53E-07 | 1.0866865 | 0.349 | 0.175 | 0.004078 | 9 | CEMIP2 |
| **698** | 3.18E-07 | 1.00897159 | 0.171 | 0.06 | 0.00849542 | 9 | CLDN1 |

| | | | | | | | |
|---|---|---|---|---|---|---|---|
| **699** | 2.33E-06 | 1.50931361 | 0.349 | 0.201 | 0.06209445 | 9 | SOD2 |
| **700** | 9.73E-06 | 1.37643059 | 0.217 | 0.099 | 0.25988297 | 9 | CD36 |
| **701** | 2.78E-05 | 1.15499712 | 0.395 | 0.244 | 0.7423831 | 9 | ANXA1 |
